# A quantitative synthesis of soil microbial effects on plant species coexistence

**DOI:** 10.1101/2021.11.12.467958

**Authors:** Xinyi Yan, Jonathan M. Levine, Gaurav S. Kandlikar

## Abstract

Soil microorganisms play a major role in shaping plant diversity, not only through their direct effects as pathogens, mutualists, and decomposers, but also by altering interactions between plants. In particular, previous research has shown that the soil community often generates frequency-dependent feedback loops among plants that can either destabilize species interactions, or generate stabilizing niche differences that promote species coexistence. However, recent insights from modern coexistence theory have shown that microbial effects on plant coexistence depend not only on these stabilizing or destabilizing effects, but also on the degree to which they generate competitive fitness differences. While many previous experiments have generated the data necessary for evaluating microbially mediated fitness differences, these effects have rarely been quantified in the literature. Here we present a meta-analysis of data from 50 studies, which we used to quantify the microbially mediated (de)stabilization and fitness differences derived from a classic plant-soil feedback model. Across 518 pairwise comparisons, we found that soil microbes generated both stabilization (or destabilization) and fitness differences, but also that the microbially mediated fitness differences dominated. As a consequence, if plants are otherwise equivalent competitors, the balance of soil microbe-generated (de)stabilization and fitness differences drives species exclusion much more frequently than coexistence or priority effects. Our work shows that microbially mediated fitness differences are an important but overlooked effect of soil microbes on plant coexistence. This finding paves the way for a more complete understanding of the processes that maintain plant biodiversity.

## Introduction

Over the past few decades, it has become clear that soil microorganisms exert substantial controls over biodiversity in plant communities (1). Much of our understanding of how soil microbes affect plant coexistence comes from empirical studies facilitated by the plant-soil feedback models of Bever and colleagues (2, 3). This framework captures the dynamics of plant species that cultivate distinct soil microorganisms, and whose performance is in turn affected by the composition of the soil community. Plant-microbe interactions can give rise to frequency-dependent feedback loops among competing plants, with positive feedbacks arising when plant species cultivate the soil microbial community in a way that advantages the cultivating species over heterospecific plants, and negative feedbacks arising when plant species are disfavored by the microbial community they cultivate. Many previous studies have compared the performance of plants experimentally grown with soil communities cultivated by conspecifics vs. heterospecifics to test for evidence of plant-soil feedbacks (reviewed in 4). On average, soil microbes generate negative rather than positive frequency dependence, especially among plant species that associate with the same mycorrhizal guild, are distantly related, have different root functional traits, or interact with each other in their native ranges (4, 5).

While we now know that soil microbes frequently generate frequency-dependence in plant population dynamics (2, 4), recent integration of plant-microbe interactions into modern coexistence theory (6–8) has highlighted that this effect only partially captures the soil community’s effects on plant coexistence (9, 10). Specifically, the frequency-dependent effects of soil microbes can destabilize plant coexistence or generate stabilizing niche differences (X-axis in Fig. 1), but the outcome of plant interactions also depends on the degree to which soil microbes generate competitive fitness differences among plants (Y-axis in Fig. 1). These fitness differences arise when the focal plant species differ in the average degree to which they benefit from mutualistic soil microbes and/or suffer from pathogenic taxa, causing one plant species to be disproportionately favored over the other, regardless of its abundance in the community. While the sign of the microbially-mediated frequency-dependence predicts whether soil microbes stabilize or destabilize the system (11, 12), whether or not soil microbes drive coexistence is ultimately determined by the balance of the stabilization (or destabilization) and fitness differences they generate (9, 10). For example, negative plant soil feedbacks are commonly inferred to mean that the microbial communities help maintain plant diversity. However, even if microbes have a stabilizing effect on plant dynamics, if they also drive large fitness differences, their net effect will be to harm plant diversity. Thus, one cannot infer the effects of plant-microbe interactions from their stabilizing or destabilizing effects alone.

**Figure 1:**
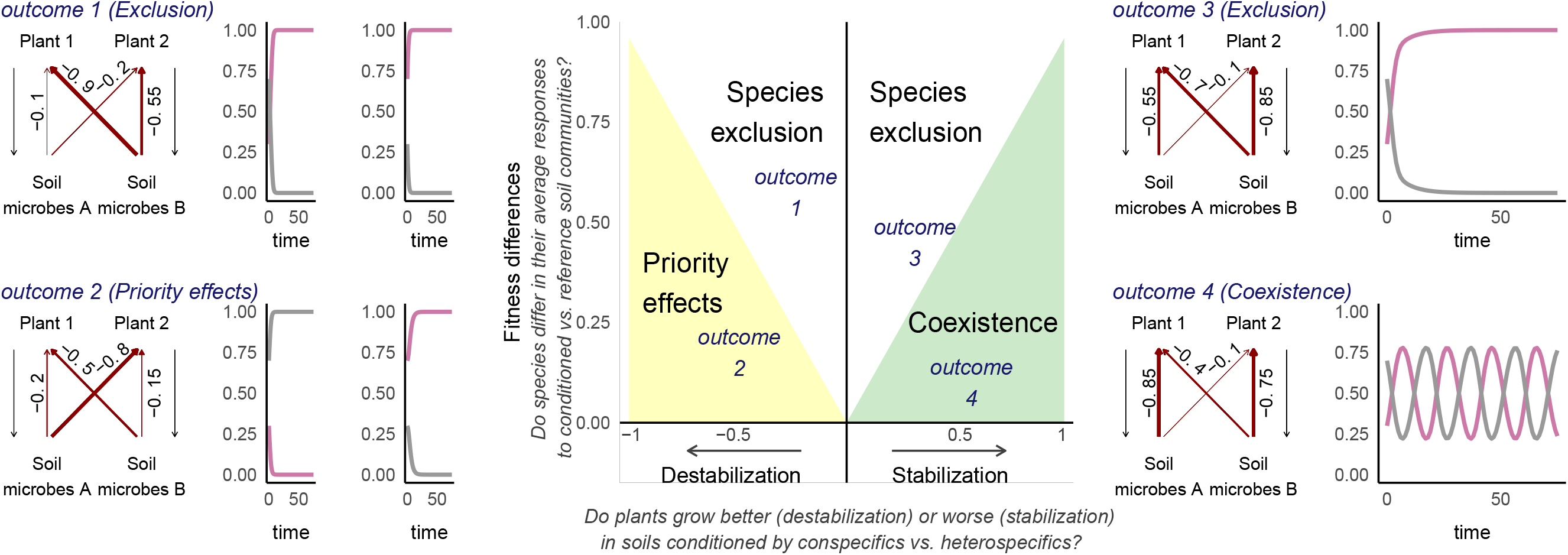
The range of possible competitive outcomes in the classic plant-soil feedback model (Bever et al. 1997), where plants only differ in their interactions with soil microbes. The middle panel shows how the balance of microbially mediated (de)stabilization and fitness difference result in coexistence, priority effects, or species exclusion. The four outcomes illustrate the population dynamics of species pairs experiencing different combinations of microbial effects, as labeled on the arrows (*m*_*iX*_ terms, which represent the effect of microbial community *X* on plant *i*). Microbially mediated (de)stabilization is calculated by comparing how plant growth is affected by conspecific- vs. heterospecific-cultivated soil microbes 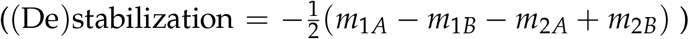, while fitness differences are calculated by comparing plant growth in either cultivated soil to growth with the reference soil community 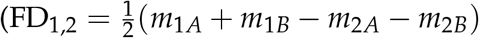; see Eqns 1-2 in Methods). Negative values of the fitness difference are possible when plant 2 is the fitness superior, but for simplicity we always set plant 1 as the superior competitor in our analyses. All outcomes are also possible if microbes have positive rather than negative effects on plant growth; see Fig S.1.

Importantly, while stabilization (or destabilization) and fitness differences have distinct effects on plant dynamics, both can arise from the same microbial players (13–15). For example, the accumulation of species-specific pathogens can stabilize plant interactions by generating negative frequency dependence, but can simultaneously create fitness differences that favor whichever plant is on average less sensitive to its pathogens. The degree to which soil microbes drive (de)stabilization vs. fitness differences is influenced by the degree of host specificity in plant-microbe interactions (10, 16, 17), and may vary with other biotic interactions and the environmental context (18, 19). In the classic plant-soil feedback framework (2), where plants are equivalent competitors except in their interactions with soil microbes, competitive outcomes between plants are solely driven by the balance between microbially mediated stabilization (or destabilization) and fitness differences (10, Fig. 1). In more complex models that incorporate other differences between plants (e.g. competition), plant competitive outcomes are determined by the joint effects of multiple processes (10). For example, microbially-mediated fitness differences can hasten competitive exclusion if they amplify other competitive asymmetries, or can favor coexistence if they equalize competitive imbalances by favoring the weaker plant species (20).

Although a few studies have recently begun testing how microbes influence the fitness differences between plant species (21–23), there have been no quantitative analyses of the literature to systematically quantify these effects, or to compare them to microbially mediated (de)stabilization. Fortunately, many experimental designs in the literature produce the data necessary to test for microbially mediated fitness differences. One simply needs results from the same two-phase experimental protocol used to quantify stabilization (or destabilization) (Fig. 2A-B), along with an additional treatment to capture plant growth in reference, uncultivated, soil communities (Fig. 2C) (23). Many studies in fact include such a reference soil community, even if they never evaluated the microbially driven fitness difference. Moreover, the microbial composition of this reference soil determines how we interpret the microbially mediated fitness differences. When the goal is to quantify the fitness differences generated specifically by the plant-soil feedback process (effects of microbial communities cultivated by either plant species), the microbial community of the reference soil should reflect the unconditioned (naive) microbial community in field soils when none of the focal species are dominant (2, 12). When the goal is to quantify the fitness differences generated by the soil microbial community as a whole, including the microbes cultivated by focal plants as well as any other microbes that are simply present in the soil irrespective of the cultivation process, plant growth in soils that lack a microbial community (sterile soils) provides the appropriate reference (Fig. 2C.ii-v).

**Figure 2:**
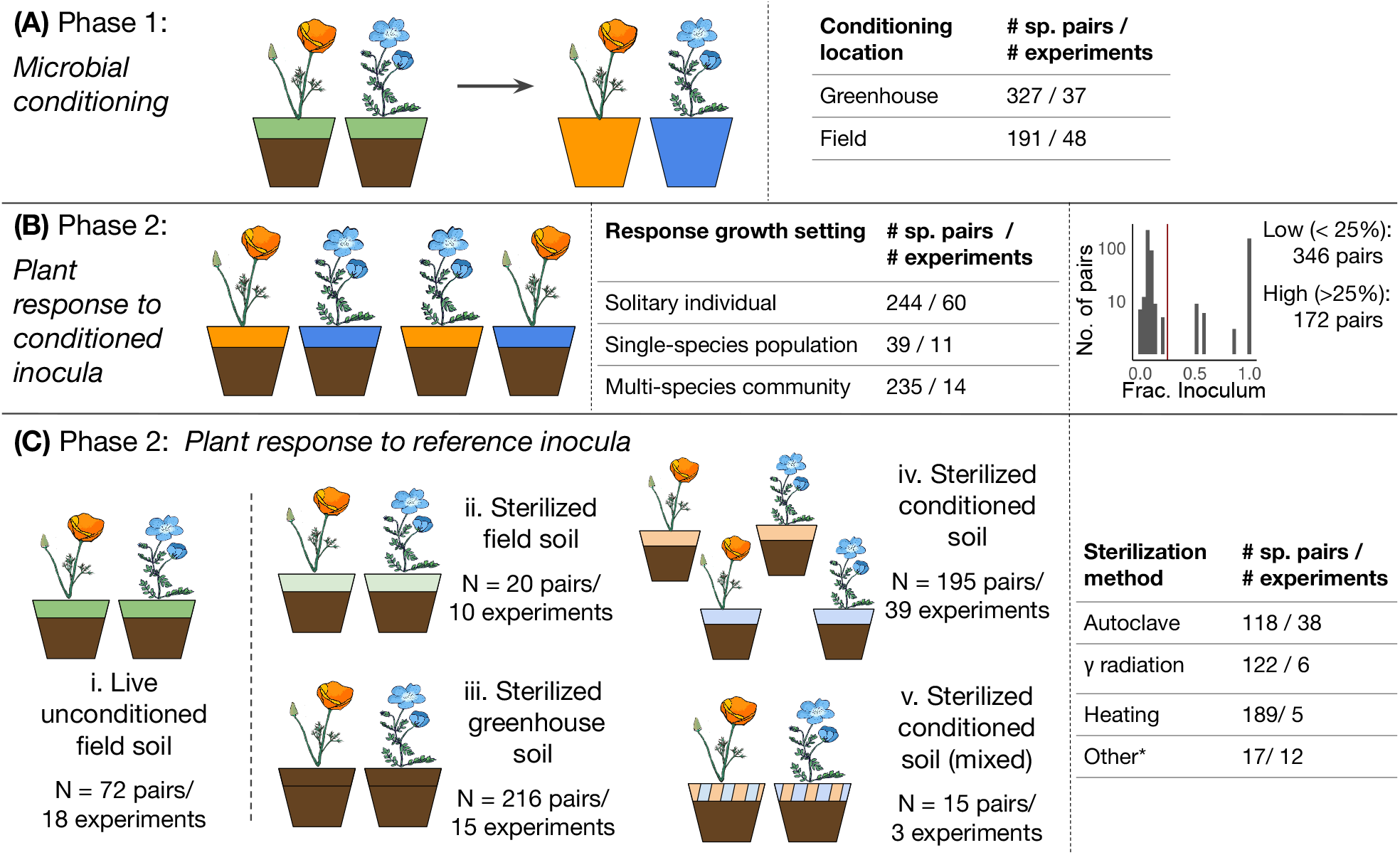
Schematic of the two-phase experimental design for studying plant-soil feedbacks and overview of the experiments in our meta-analysis. In the microbial conditioning phase (Panel A), plants grow in common sterilized background soils (brown) that contain a live field-collected soil inoculum (green). As they grow, plants impart species-specific legacies on abiotic and biotic soil properties, resulting in distinct soils (orange or blue). Quantifying microbially mediated stabilization (or destabilization) only requires measuring growth of the focal plants in the plant response phase in soils inoculated with conspecific- and heterospecific-cultivated microbes (Panel B). Quantifying the microbially mediated fitness difference also requires quantifying plant growth in soils that lack a species-specific soil community (unconditioned or sterilized soil, Panel C). The inoculum for this phase can be the same live, unconditioned field soil used to start phase 1 (green, option i), or a sterile inoculum. In practice, researchers have sterilized various types of soil to use as reference inocula, including field soil (light green, option ii), greenhouse soil (brown, option iii), or soil conditioned by the focal plants (light orange or light blue, options iv-v). The numbers in each panel summarize the experimental methodology of the studies in our meta-dataset, see Methods for details.

Here we synthesize data from the plant soil feedback literature with recently derived theoretical metrics (10) to present a meta-analysis evaluating how soil microbes affect plant coexistence through both the (de)stabilization and the fitness differences they generate. We frame our analyses around two main questions: **Q1)** What is the relative magnitude of microbially driven stabilization (or destabilization) versus fitness differences, and how is it affected by experimental treatments? **Q2)** Across species pairs, what is the net effect of microbes on competitive outcomes? In other words, how frequently does the balance of microbially-mediated stabilization (or destabilization) and fitness differences predict species coexistence, exclusion, or priority effects? We explored these questions separately for species pairs whose growth in live unconditioned soil communities provided the reference condition for calculating microbially mediated fitness difference, and for pairs where growth in sterile soils provided the reference. This lets us evaluate both the specific effects of microbes accumulating in soils specifically due to the cultivation process (using reference soils with live unconditioned microbial communities), and the more general effect of soil microbes on plant coexistence (using sterile reference soils).

## Results

### Data overview

Our meta-analysis included data for 263 unique species pairs across 85 experiments (from 50 studies) with all treatments required to calculate the microbially mediated stabilization (or destabilization) and fitness differences in Bever et al.’s model of plant-soil feedback (see Appendix S1 for PRISMA chart). After accounting for multiple experiments within study (e.g. under low or high watering, under different light regimes), we quantified 72 effect sizes (from 18 experiments) for these metrics with growth in a live unconditioned soil community serving as the reference condition, and 446 effect sizes (from 67 experiments) for the pairwise (de)stabilization and fitness difference metrics with growth in sterile soil serving as the reference condition.

### Q1: Relative strength of microbially mediated (de)stabilization and fitness differences

Across all species pairs, microbes generated both (de)stabilization and fitness differences (all mean effect sizes > 0 with p < 0.001, Fig. 3). The strength of stabilization (or destabilization) was calculated by comparing how the growth of each species is affected by soil conditioned by conspecifics vs. heterospecifics (absolute value of Eqn. 1 in Methods), and fitness differences were calculated by comparing how each plant’s growth is on average affected by the two cultivated soil communities vs. the reference soil (Eqn. 2). We further found that the fitness differences generated by soil microbes were substantially larger than the magnitude of their (de)stabilizing effects in both reference soil datasets (live reference: F = 21.1, n = 72, df = 1, 8.3; p = 0.0016; sterile reference: F = 33.1, n = 446, df = 1, 50.7; p < 0.001). There was also substantial heterogeneity in both effects, regardless of reference soil type (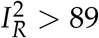 for all effect sizes, Fig. 3). Among experiments with growth in sterile soil as a reference, a larger microbially mediated fitness difference than stabilization or destabilization was consistent across experiments that used sterilized greenhouse soil or sterilized single-species cultivated soil as the reference inoculum, measured plants grown as solitary individuals or in multispecies communities, or sterilized soils by autoclaving, gamma irradiating, or heating (Fig. S.3A). Among species pairs for which growth in live unconditioned soil provided the reference condition, a larger fitness difference than (de)stabilization was consistent across experiments which conducted the conditioning phase in a greenhouse, used a high fraction of inoculum, or grew plants as individuals in the response phase (Fig. S.3B). None of these results changed with a univariate analyses approach (Fig. S.4), different specifications of the variance-covariance matrices for the multivariate models (Fig. S.6), or with the exclusion of influential experiments (Fig S.5).

**Figure 3:**
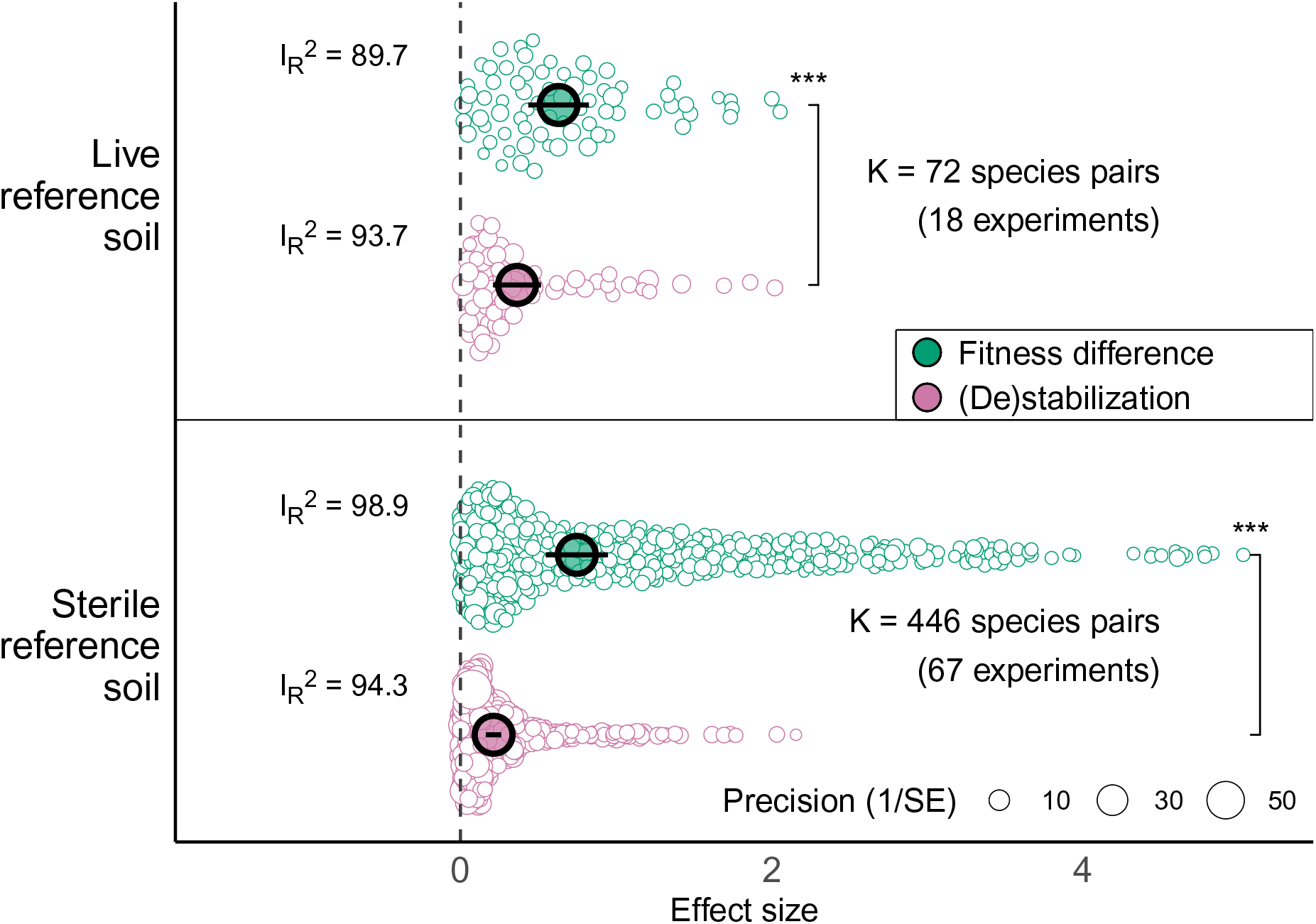
Magnitude (absolute value) of pairwise (de)stabilization and fitness differences mediated by soil microbes. Solid points show the mean effect sizes from multivariate meta-analyses, and black lines show their 95% confidence intervals. Hollow point shows the effect sizes for each pairwise comparison, and their size indicates the precision of the estimate. Brackets show that within each reference type, fitness differences were significantly larger than the strength of microbially mediated (de)stabilization (Wald tests; p = 0.0016 for live reference and p < 0.001 for steril reference).

### Q2: How frequently do microbes predict different competitive outcomes among plants

Our next goal was to evaluate whether the balance of microbially driven (de)stabilization and fitness differences generally predicts plant coexistence, priority effects, or species exclusion. We addressed this goal by sampling 10000 times from probability distributions reflecting the mean values and uncertainties of plant biomass in different soil types from the original studies to quantify pairwise stabilization (or destabilization) and fitness differences and then predicting coexistence outcomes (see Eqns. 4-5 in Methods). From this sampling approach, we found that soil microbes were predicted to drive species exclusion much more frequently than coexistence or priority effects. This result was consistent regardless of whether we focused on species pairs with reference growth on live or sterile soils (Fig. 4, live reference: 69.4% of sample draws predict exclusion; sterile reference: 81.2% of sample draws predict exclusion). Specifically, competitive exclusion was the most common outcome, followed by coexistence and priority effects. This was found in all methodological subgroups except among experiments that used conditioned soils that had been mixed prior to Phase 2 as the reference inoculum; here, coexistence was the dominant outcome (Tables S1-S2).

**Figure 4:**
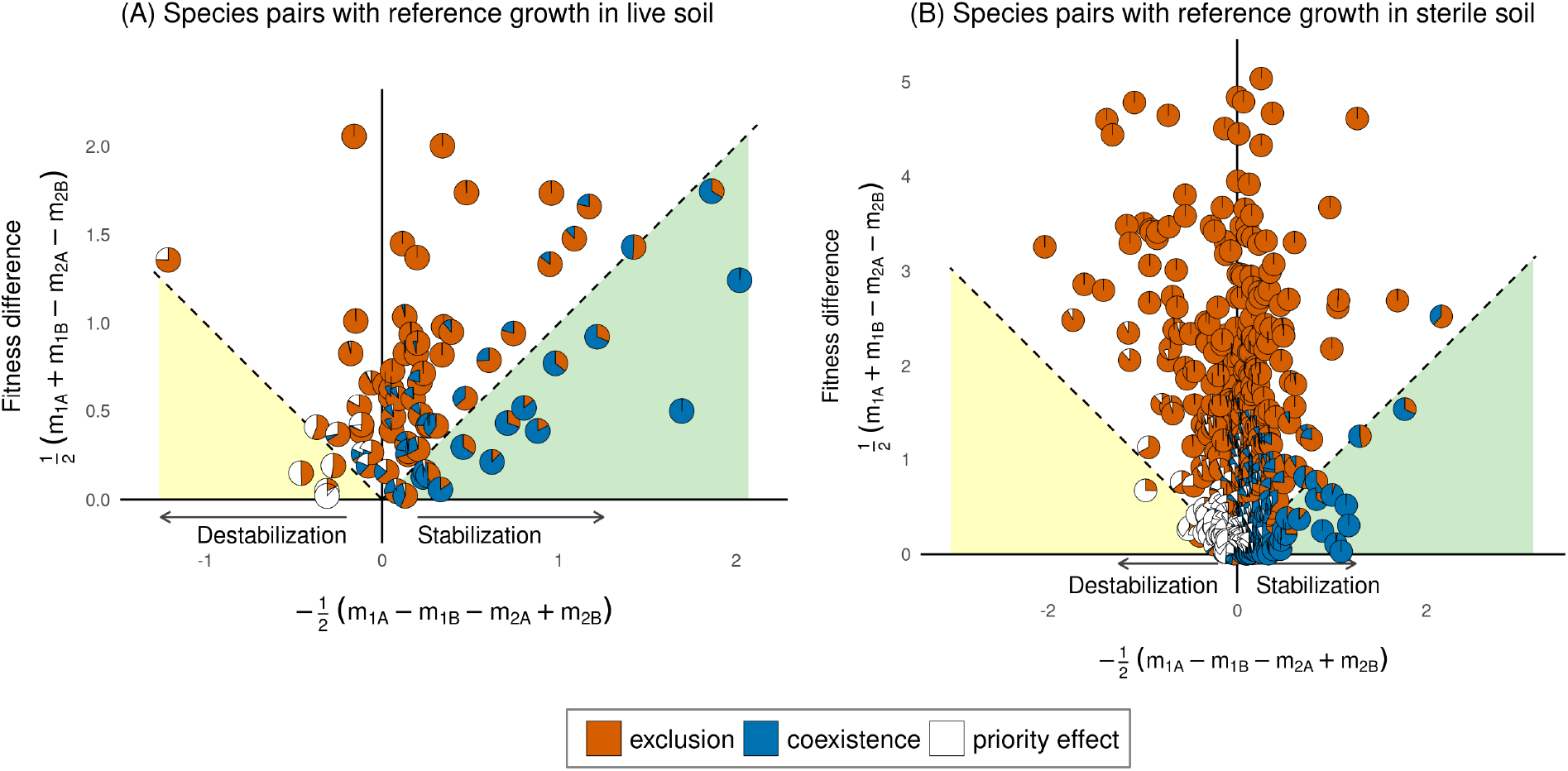
Outcomes of microbially-mediated pairwise plant interactions among pairs with plant growth in live, unconditioned soils (panel A, n = 72) or sterile soils (panel B, n = 446) providing the reference. Each species pair is represented by a pie chart, whose x and y coordinates show the mean strength of (de)stabilization and fitness difference, and whose colors indicate the proportion of sample draws that result in each coexistence outcome.

Consistent with these findings on the predicted competitive outcome, among the subset of 319 species pairs in which soil microbes were predicted to generate stabilizing niche differences, the overall strength of this stabilization was weaker than the microbially mediated fitness difference, irrespective of whether growth on sterile or live soil provided the reference (Fig. S.7A, live reference: F = 22.1, n = 55, df = 1, 6.66; p = 0.0025; sterile reference: F = 25.9, n = 264, df = 1, 39.9; p <0.001). Among the 199 pairs whose interactions were destabilized by soil microbes, the destabilization was significantly weaker than microbially mediated fitness difference for pairs where growth on sterile but not live soils provided the reference (Fig. S.7B, live reference: F = 2.71.1, n = 17, df = 1, 5.72; p = 0.153; sterile reference: F = 36.3, n = 182, df = 1, 34; p <0.001). Both lines of evidence suggest that the balance of microbially driven (de)stabilization and fitness differences primarily drives plant species exclusion rather than coexistence or priority effects.

## Discussion

Modern coexistence theory highlights that biological differences between species can simultaneously affect their coexistence by generating stabilizing or destabilizing frequency dependence, and by equalizing or exaggerating fitness inequalities (6, 13). Despite growing evidence that soil microbes often generate feedback loops that stabilize or destabilize plant dynamics (4), whether plant-microbe interactions cause coexistence, priority effects, or exclusion among plants has remained unclear because microbially mediated fitness differences have been very rarely quantified. Our synthesis integrated 518 pairwise stabilization (or destabilization) and fitness difference comparisons from 50 studies and found that microbially mediated fitness differences represent a dominant consequence of soil microbes on plant competitive outcomes. Specifically, we found that across all species pairs, microbes drove large fitness differences (effect sizes > 0 with p-values < 0.001) that favor one plant over the other (Fig. 3). Moreover, these fitness differences tend to be more consequential for plant population dynamics than any stabilizing or destabilizing effects (Figs. 3 and S.7).

Our analysis underscores the importance of precisely framing the question of how microbes influence plant coexistence. Answering if plant-soil feedbacks generate negative or positive frequency-dependence in plant interactions only requires quantifying the microbially mediated stabilization or destabilization; no reference soil treatment is needed. This has been the primary goal of previous plant-soil feedback theory and experiments (2, 11, 24). Consistent with previous meta-analyses (4, 25), we found that microbes have an overall stabilizing effect on plant interactions (Fig. S.8). But to answer whether microbes drive plant *coexistence* requires comparing the strengths of (de)stabilization and fitness differences. When the goal is to specifically assess how *soil microbes cultivated in the plant-soil feedback process* influence coexistence, we need measures of (de)stabilization and the fitness difference with respect to plant growth in an unconditioned soil community (i.e. live soil reference) (2, 12). Alternatively, when the goal is to more broadly assess how the *soil microbial community as a whole* influences coexistence, we need measures of the (de)stabilization, and the fitness difference with respect to plant growth in soils lacking a microbial community (i.e. sterile reference soil).

Regardless of whether one is interested in the effects of the soil community as a whole, or of only those microbes that are specifically conditioned during the feedback process, our results indicate that plant-microbe interactions largely drive species exclusion rather than coexistence or priority effects. This tendency is slightly more pronounced when we consider the coexistence consequences of the microbial community as a whole, than only of the soil communities cultivated by the plants. Interactions between plants and the whole soil community result in coexistence in only 12.8% of species pairs, about half as frequently as interactions between plants and the soil communities they cultivate (22.2%) (Fig. 4). It is possible that the fitness differences generated by the background microbial community either overturn or further exaggerate the effects of microbes cultivated in the feedback process. Quantifying these relationships requires data from studies that grew plants on both live and sterile references, but we could not perform this analysis because we only had 4 such studies in our meta-dataset. Our analysis also revealed substantial variation in the strength of microbially mediated fitness differences among species pairs (high 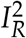 for all effect sizes). Exploring how variation in abiotic conditions and functional traits of plant and soil microorganisms contribute to this heterogeneity is a key next step (26–28).

While our study was not designed to identify the mechanisms through which soil microbes generate fitness differences among plants, previous studies suggest a few likely avenues through which these differences arise. First, co-occurring plant species can vary greatly in their responsiveness to soil mutualists (30). In such cases, the soil microbial community will give a fitness advantage to the more responsive species, an effect that might be most evident when growth of both plant species in sterile soils is the reference. In nature, the growth of small-seeded plant species and of many invasive species tends to be more responsive to mycorrhizal fungi with large host ranges (31, 32), suggesting that the soil community likely generates fitness advantages in favor of such plants. Similarly, soilborne pathogens can generate fitness differences among plants that vary in their defense against microbial enemies (33). In such cases, microbially mediated fitness differences should be evident even when using plant growth in live unconditioned soil as the reference, especially if the plants differ in their sensitivity to both generalist and specialist pathogens (10). Furthermore, rather than directly affecting plant performance as mutualists or pathogens, soil microbes may also differentially affect plant species fitness by altering soil nutrient dynamics, for example by directly competing with plant species for nitrogen (34) or by altering litter decomposition processes (35). Finally, plant fitness differences may also arise through interactive effects of plants on the biotic and abiotic properties of the soil, a possibility that has received relatively little attention in empirical studies (36). In all of these scenarios, microbially mediated fitness differences may act to equalize or exaggerate other competitive asymmetries among the focal species, a possibility that we discuss next.

The fitness differences we have identified drive species exclusion in simple models of plant-microbe interactions where plants are otherwise equal competitors (2), but their joint effects with other competitive asymmetries may enable coexistence in more complex settings (9, 10). For example, microbes may equalize fitness among plants that experience a tradeoff between their competitive ability and their defense against microbial pathogens. Evidence is mounting that such a tradeoff may be captured by the root economics spectrum, such that soil microbes might equalize fitness between species with contrasting strategies on this spectrum (5, 37–39). Mycorrhizal fungi may also equalize plant competitive imbalances via the formation of common mycelial networks if nutrients are transferred from dominant to inferior plants (14, 40). On the other hand, it is also possible for soil microbes to amplify other fitness inequalities among plants. This may happen if, for example, plant species that are inferior competitors are also more susceptible to attack by generalist pathogens because they are less able to defend themselves than plant species with superior competitive ability. Future studies that adopt slightly more complex designs like those outlined by Ke and Wan (9) offer an ideal path forward to build this understanding.

Here we have synthesized data from decades of research on plant-microbe interactions with the more recent theoretical insight that coexistence is influenced by the relative strength of interspecific (de)stabilization and fitness differences. We find that microbially mediated fitness differences are a major and prevailing effect of soil microbes on plant coexistence, whose mechanisms and consequences need to be further explored in future empirical studies. These fitness differences often drive species exclusion when plants are otherwise equivalent competitors, but may favor diversity if they generate tradeoffs that equalize plant fitness. Regardless of which of these outcomes is ultimately supported by future empirical investigation, our work suggests that the overwhelming effects of microbes on plant competitive outcomes is through the fitness differences, not the stabilization or destabilization as classically assumed. More generally, our work highlights the value of linking empirical data with theoretical models of species interactions to study how individual mechanisms scale up to affect the dynamics of whole ecological communities.

## Methods

### Study selection

The list of publications that we screened for our meta-analysis came from two sources (See Appendix S1 for PRISMA chart). The first source was the meta-dataset compiled in Crawford et al. (2019) (4), which included 69 studies with all the relevant treatments to calculate stabilization or destabilization mediated by plant-soil feedbacks. To compile these data, Crawford et al. had conducted a Web of Science search through 17 January 2018 using the keyword ‘plant AND soil AND feedback’ within the ‘Ecology’ and ‘Environmental Science’ categories, and from secondary citations. They filtered the resulting studies following four criteria: (1) The study included at least two plant species (or two genotypes as in one study, which for simplicity we include as heterospecifics) growing on soils that were previously cultivated by conspecifics and heterospecifics. (2) The study used a factorial design with each species in a pair grown in conspecific and heterospecific-cultivated soil. (3) Each soil inoculum reflected the conditioning effects of a single plant species, such that studies that pooled soils from multiple species were excluded. (4) Measures of mean, error, and sample size for plant performance in the different soil treatments could be acquired from the publication or by contacting the authors. We further screened the resulting 69 studies from Crawford et al’s search with one additional criterion: (5) the study also measured plant growth in a reference soil environment that had either been inoculated with a microbial community that was not specifically cultivated by conspecifics or heterospecifics (e.g. soil collected in the field, away from the focal species; Fig. 2C.i), or was lacking a microbial community (e.g. autoclaved greenhouse soil; Fig. 2C.ii-v). 32 out of the 69 studies passed this filter.

We supplemented this dataset with a search from Web of Science Core Collection on 13 July 2021 using the same keywords and identified 391 studies published since 2018. We filtered these studies following the five criteria above. This yielded 18 studies, for a total of 50 papers from which we extracted information (21, 23, 31, 36, 41–86). If a study had multiple experiments across different soil sources, resource gradients, or competition intensities, we compiled data separately for each experiment.

### Data, metadata, and moderator extraction

We used whole-plant biomass, above-ground biomass, or other proxies of biomass as measures for plant growth. For the 32 relevant studies included in Crawford et al.’s dataset, which already included plant growth in conspecific- and heterospecific-trained soil, we simply extracted data for each plant species’ growth in the reference soil microbial community. For the 18 additional studies we identified in our follow-up literature search, we extracted data for plants growing in conspecific-conditioned, heterospecific-conditioned, and reference soils. We directly extracted the mean, standard error, and sample size of the plant growth data from text, tables, or supplemental datasets when available, or by requesting data from authors. When biomass data were only available in figures, we manually extracted them using Webplot digitizer (87). We verified that the biomass estimates from our manual data extraction were consistent with Crawford et al.’s.

We also recorded methodological factors that could influence microbial effects on plant growth. First, we recorded whether the reference soil contained a live microbial community that was not specific to a single plant species (e.g. live field-collected inoculum), or no microbial community (e.g. sterilized field-collected or greenhouse soil, see details below), and analyzed species pairs with live vs. sterile reference soils separately. In addition, we collected the following information (Fig. 2): (1) the environment where the first (‘conditioning’) phase took place (greenhouse or field), (2) whether pots in the second (‘response’) phase of the experiment received a low (<=25%) or high (>25%) fraction of previously-conditioned inoculum, (3) the setting for the response phase (whether plants were grown as individuals, in populations of single species, or in mixed species communities). For experiments that use the sterile reference soil, we also recorded (4) the source of sterilized reference inoculum (mixed-species cultivated soil, species specific cultivated soil, field soil, or greenhouse soil), and (5) soil sterilization method (autoclaving, gamma irradiation, heating, or other minority methods).

### Quantifying pairwise (de)stabilization and fitness difference

We used metrics derived from the classic plant-soil feedback model (2) to calculate microbially driven (de)stabilization and fitness difference (10):

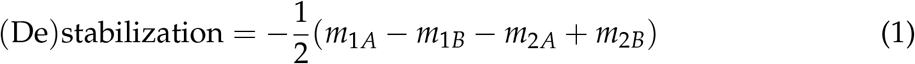

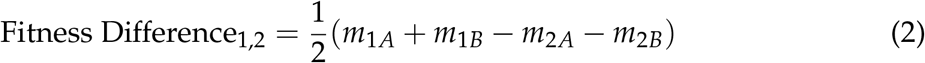

Each *m*_*iX*_ term represents the performance of plant *i* in soil *X* relative to its performance in the reference soil, and is calculated as shown in the next paragraph. Positive values of the (de)stabilization metric (Eqn 1) correspond to stabilizing effects of soil microbes, while negative values indicate destabilizing effects. The absolute value of this metric indicates the magnitude of microbially-mediated frequency dependence, regardless of whether it stabilizes or destabilizes plant interactions. Note that the term in Eqn 1 is equal to 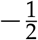*I*_*S*_, the feedback metric originally derived by Bever and colleagues, which was used in Crawford et al.’s meta-analysis (4). To calculate fitness differences, we always set species 1 as the fitness superior and species 2 as the inferior competitor; thus, the mean fitness difference for every species pair has a positive value. Microbes result in plant coexistence when they generate stronger stabilizing niche differences than fitness differences:

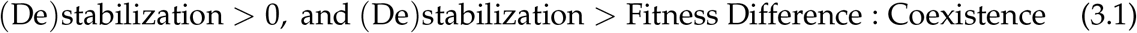

Priority effects arise when microbes generate stronger destabilizing effects than fitness differences:

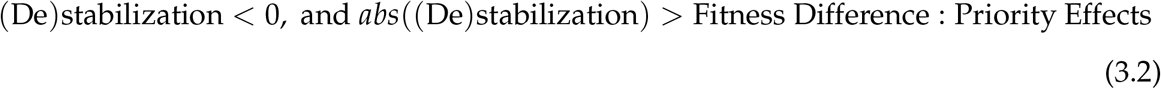

Finally, soil microbes drive species exclusion when they drive stronger fitness differences than (de)stabilization:

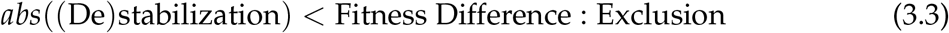

For our synthesis, we operationalized Eqns 1-2 by quantifying each *m*_*iX*_ term as *ln*(*G*_*iX*_) − *ln*(*G*_*i,Re f*_), where *G*_*iX*_ represents the mean growth of a focal plant species *i* with soil community *X*, and *G*_*i,Re f*_ is the mean growth of plant *i* in the reference community. Due to arithmetic, *G*_*i,Re f*_ drops out from stabilization, but is required for quantifying fitness differences (23). The effect sizes can also be rearranged into a log response ratio form (Appendix S2):

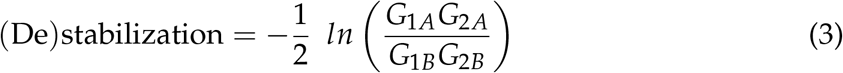

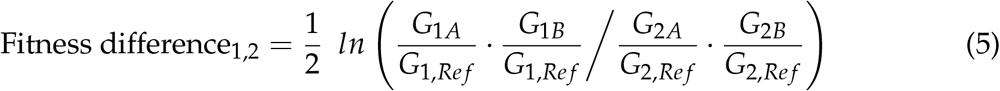

Following Lajeunesse (2011) (88), we also derived the variances equations of each effect size (see Appendix S2 for details):

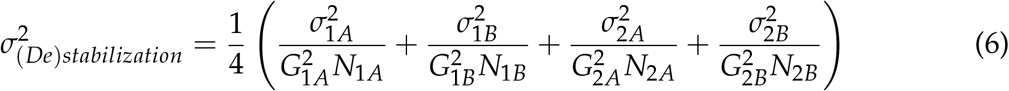

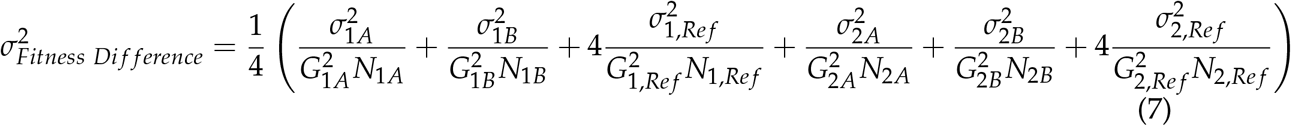

Where 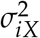 the variance and *N*_*iX*_ is the sample size for *G*_*iX*_.

### Analysis

#### Analysis for Q1

The goal of our first analysis was to evaluate the relative magnitude of microbially mediated (de)stabilization and fitness differences among plant competitors, and how it depends on experimental methodologies. We addressed this question with a multilevel multivariate metaanalysis of the absolute values of (de)stabilization and fitness difference metrics (Eqn. 4-5) using `metafor::rma.mv` (89). The model included a fixed effect for the effect size type ((de)stabilization/fitness differences). The random effects were for each effect size type within an experiment and within a species pair (such that each effect size measurement had its own random effect). The model also weighed the effect size estimates based on their variances (Eqn. 6-7), such that more uncertain estimates carried less weight. Because we lacked information about the covariance between effect sizes in the original studies, we imputed a covariance matrix using `clubSandwich::impute_covariance_matrix` (90), assuming that within experiments, (de)stabilization and fitness differences of each species pair are have an intermediate correlation (r = 0.5). We verified that our results were not sensitive to this decision by repeating the analyses for covariance matrices generated with assumed correlations of −0.5, 0.1 and 0.9. We fit separate models for species pairs where the reference soil for growth was either live or sterile. We quantified heterogeneity using the 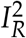 metric (91), and evaluated the likelihood profiles to ensure we had not overfit models (Fig. S.2). We did not test for publication bias because the substantial heterogeneity and non-independence of effect sizes violates assumptions of relevant tests (92). We used Cook’s distance to identify influential effect sizes, and verified that our results were robust against excluding influential points.

We evaluated the models using tools for robust variance estimation, which are well suited to handle multiple dependent effect sizes within meta-analytical models (93, 94). In each model, we first evaluated whether the magnitude of each effect size ((de)stabilization and fitness difference) was different from 0 using t-tests implemented in `clubSandwich::coef_test`. This function evaluates significance using sandwich estimators for the standard errors, and Satterthwaite-corrected degrees of freedom (90). Next, we tested whether soil microbes generated stronger (de)stabilization or fitness differences using linear contrasts implemented in `clubSandwich::Wald_test`, using bias reduced linearization estimate to correct for small sample errors (`vcov = “CR2”`) (90). We also evaluated the robustness of our results with equivalent univariate analyses of (de)stabilization and fitness differences.

We also conducted moderator analyses to evaluate if variation in the effect size estimates was driven by methodological differences. We separately tested whether the magnitude of (de)stabilization vs fitness differences varied according to differences in the conditioning phase environment (greenhouse/field), the fraction of inoculum used in the response phase (low (<25%)/high), the response phase setting (individual/population/community), the source of the sterile soil reference (greenhouse soil, field soil, previously-conditioned soil), or the sterilization method (autoclaving, gamma irradiation, heating, or ‘other’ (pasteurization, steaming, or unspecified sterilization)). We built the meta-analytical models as above, each with one additional fixed effect that captured the relevant methodological factor. This resulted in five models for the sterile reference dataset, and three models for the live reference dataset. We again used linear contrasts to test whether microbes generated stronger (de)stabilization or fitness differences under different experimental conditions.

#### Analysis for Q2

The goal of our second question was to evaluate how frequently the balance of microbially mediated (de)stabilization and fitness differences drives plant species coexistence, priority effects, or competitive exclusion in the classic plant-soil feedback model (2). We addressed this question with a sampling approach that incorporated the mean estimates of plant growth in different microbial contexts, as well as the uncertainty in these estimates. In each sample draw, we drew one value for each of the 6 *G*_*iX*_ terms per species pair from a normal distribution with the mean and standard deviation as reported in the original experiment. Following Eqns 4-5, we used these draws to calculate the (de)stabilization and fitness differences for each species pair, which we used to render a coexistence outcome (Eqns 3). We repeated this procedure 10000 times per species pair, and summarized the frequency of each outcome (species coexistence/priority effects/exclusion) across all sample draws.

We supplemented this sampling approach with a pair of multivariate meta-analyses similar to those in Q1, conducted separately for species pairs with stabilizing niche differences (mean value of Eqn. 1 > 0) and those where microbes destabilize plant interactions (mean value of Eqn. 1 < 0). This lets us evaluate whether plant-soil interactions that stabilize plant dynamics do so strongly enough to overcome microbially mediated fitness differences to result in coexistence rather than exclusion in the classic plant-soil feedback model. Similarly, we can evaluate whether soil microbes with destabilizing effects generally result in priority effects among plants, or if destabilizing effects of microbes are weaker than the fitness differences they generate.

The analyses were conducted in `R` version 4.1.2 (95). Code to recreate all analyses is available in Appendix S3 and will be archived along with the meta-dataset upon publication.

## Acknowledgements

We respectfully acknowledge that the University of Missouri is located on the traditional, ancestral lands of the Osage, Omaha, and Kaw peoples, among others; and that the University of Texas is on the indigenous land of the Tonkawa, Comanche, and Apache communities, among others. We thank all authors who made their data available for this analysis. We acknowledge Farrior and Wolf Labs at UT, Amarasekare Lab at UCLA, Sullivan, Halsey, and Brown Lab at UCLA, Sullivan, Halsey, and Brown Labs at MU, Madeline Cowen, Po-Ju Ke, and the MU DataPhiles group for help at various stages. We also thank Bindu Viswanathan and Michael Mahometa for conversations on the statistical problem. XY is partially supported by UT Enhanced Support Fellowship

## Supplemental figures

**Figure S.1:**
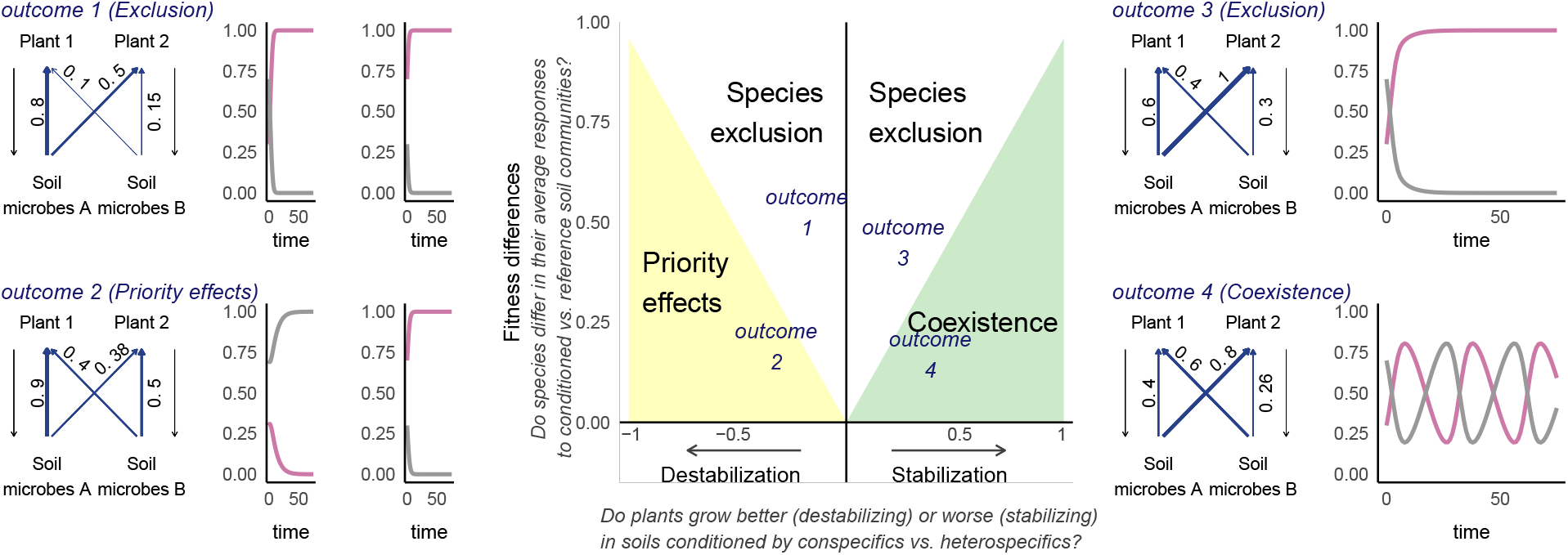
In the four outcomes of the Main Text Fig. 1, we show that plant-soil feedbacks can result in priotity effects, coexistence, or competitive exclusion when soil microbes generally suppress plant growth (negative values of *m*_*iX*_). Here, we show that all four outcomes even soil microbes promote the growth of both plant species (positive values of *m*_*iX*_).

**Figure S.2:**
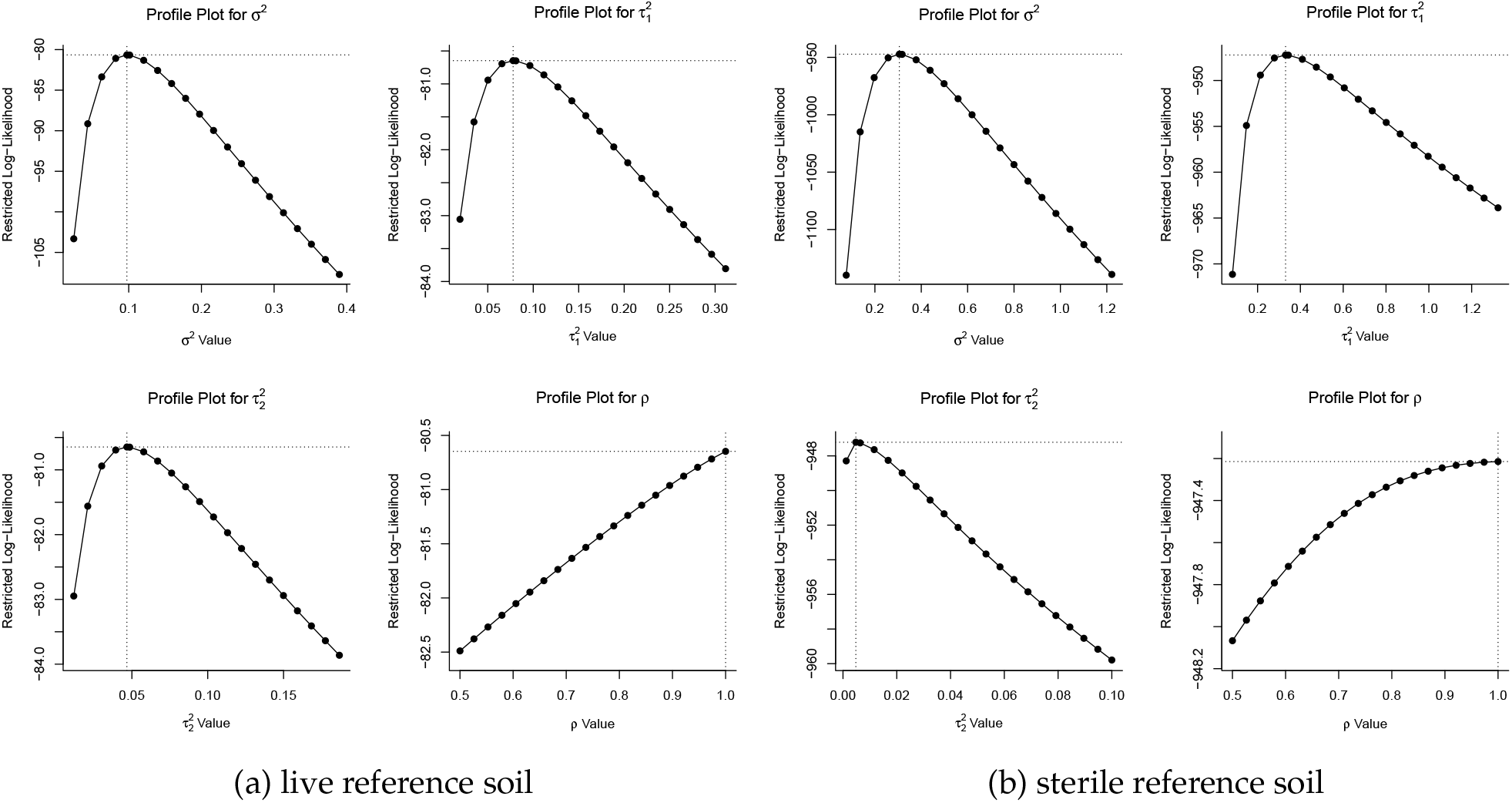
Likelihood profiles for Q1 model parameters, generated with `metafor::profile()`

**Figure S.3:**
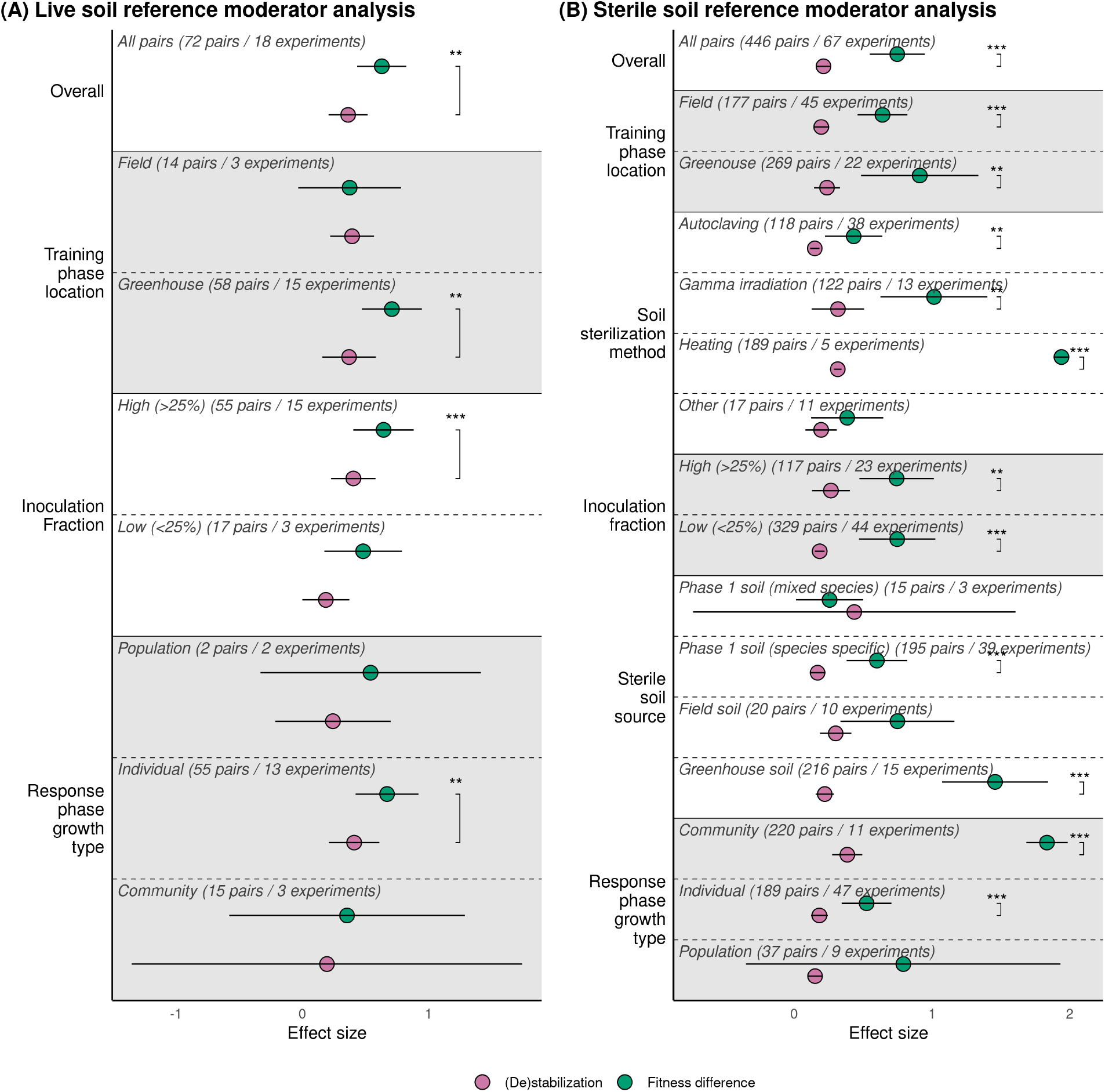
Moderator analyses for Q1. Soil points indicate the overall effect sizes for pairs that were studied using the relevant methodological approach, and whiskers indicate the 95% confidence intervals. Significance brackets indicate those pairwise (De)stabilization/Fitness difference comparisons that are significantly different from one another.

**Figure S.4:**
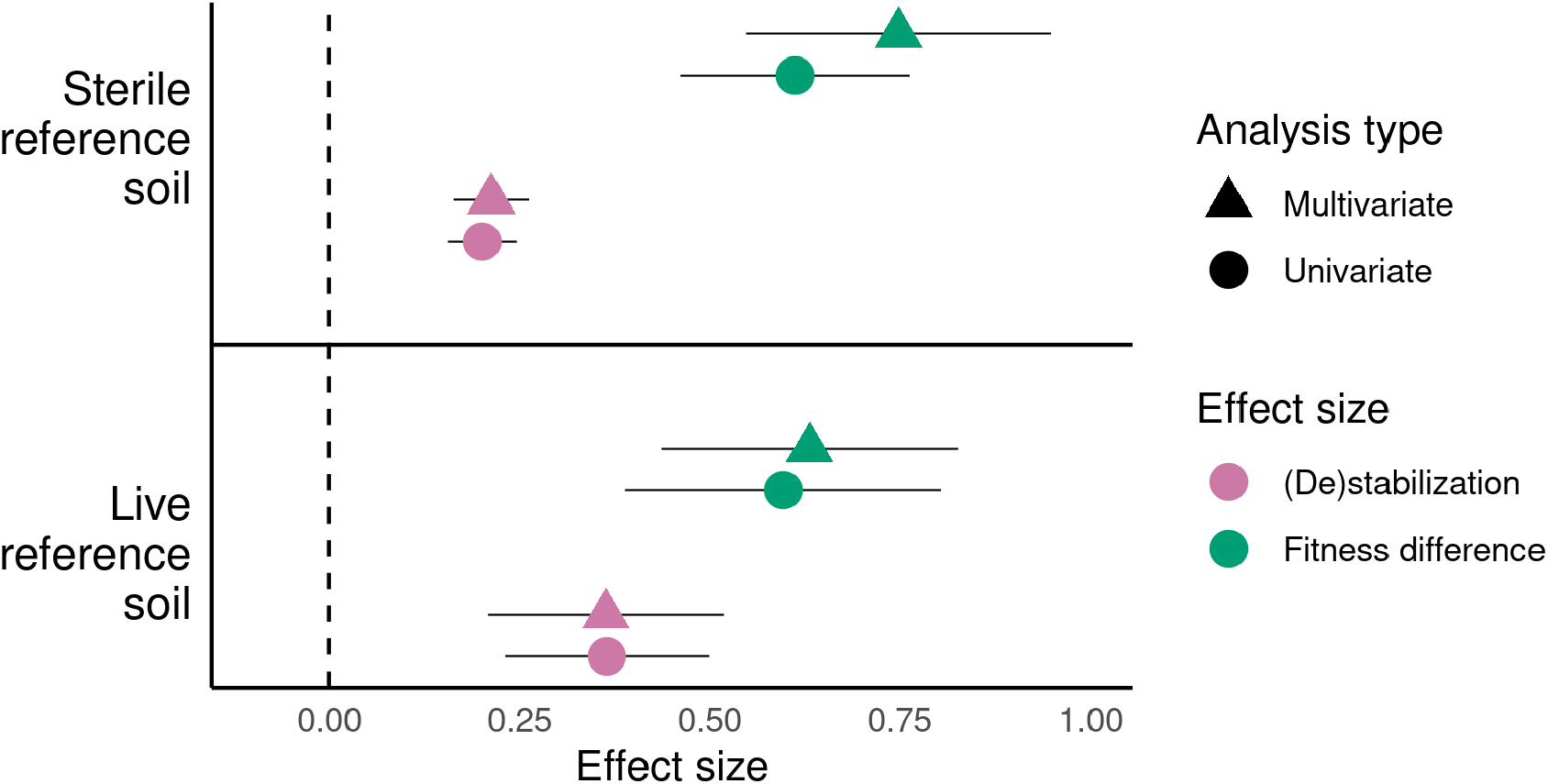
Comparison of (de)stabilization and fitness difference effect sizes (Main text Q1) when analyzed through multivariate models vs. through separate univariate models of the two effect sizes.

**Figure S.5:**
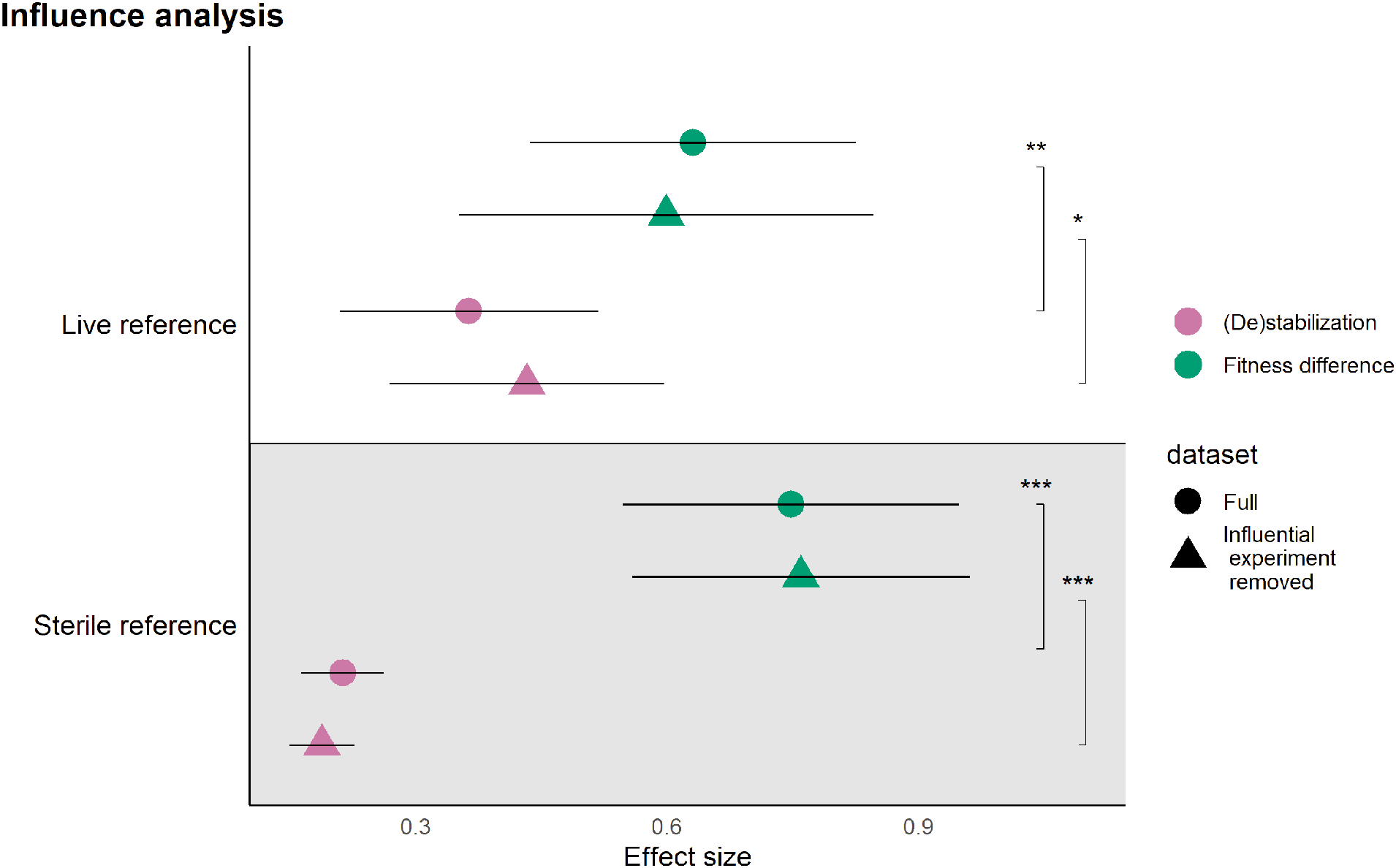
The effect of excluding influential studies. Kandlikar et al. 2019 and Miller et al. 2015 were identified as influential experiments in the live reference dataset, and Hendriks et al. 2013 in the sterile reference dataset. Fitness difference remains significantly larger than (de)stabilization in both reference datasets after removing these studies.

**Figure S.6:**
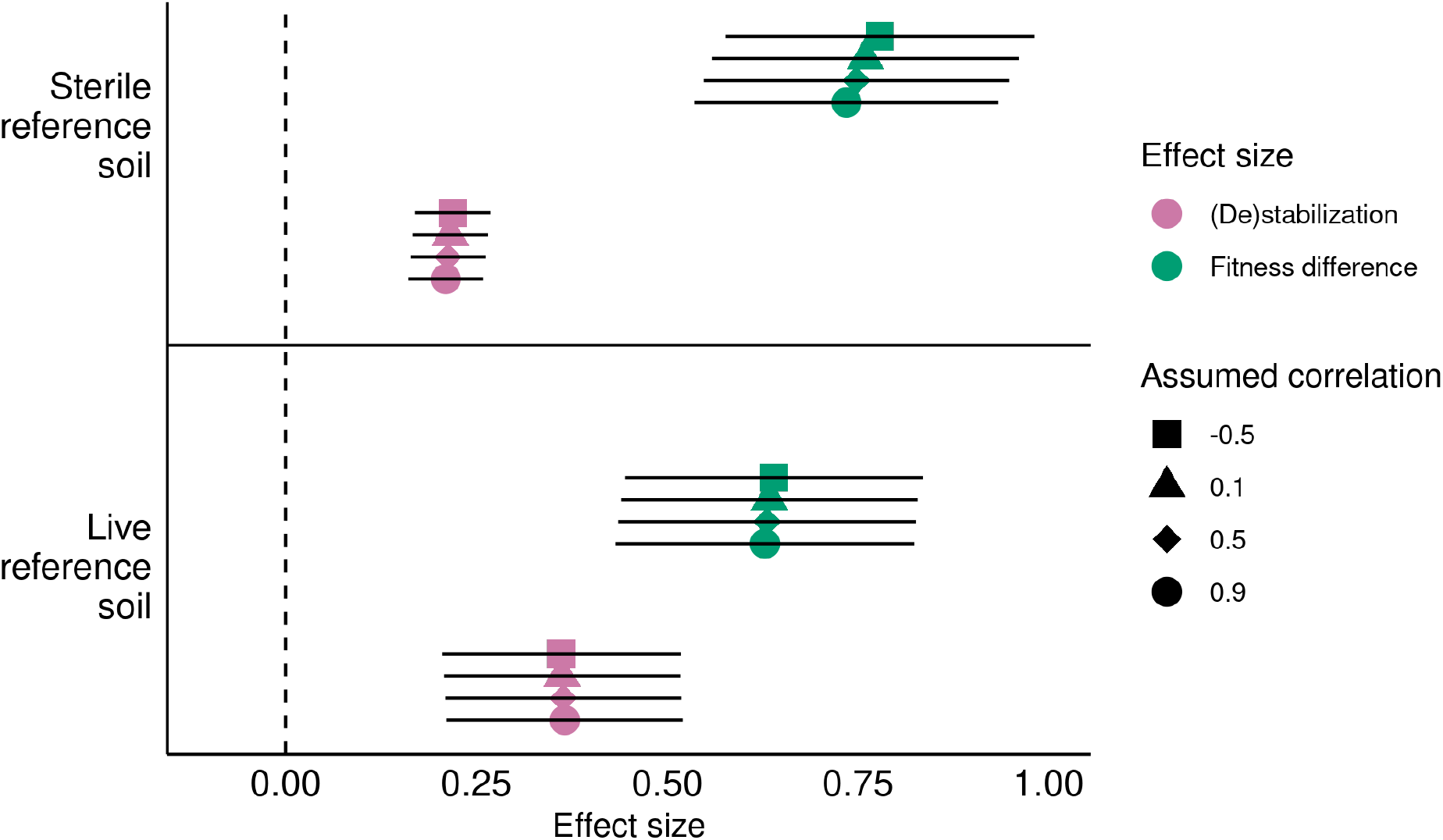
Comparison of (de)stabilization and fitness difference effect sizes (Main text Q1) when analyzed with variance-covariance matrices derived assuming different strengths of correlations between (de)stabilization and fitness differences among species pairs.

**Figure S.7:**
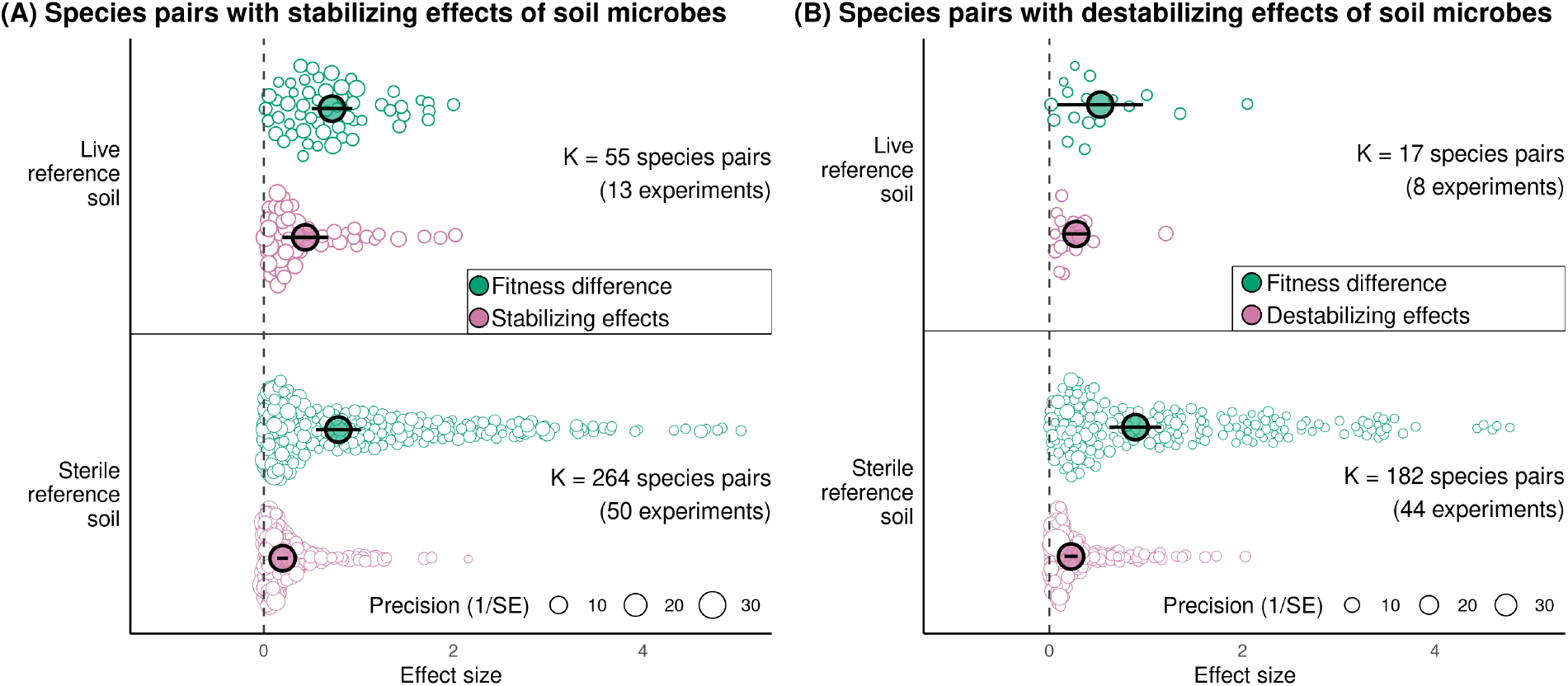
Effect sizes from meta-analyses of species pairs with stabilizing (A) or destabilizing (B) effects of microbes. Microbially mediated fitness differences are stronger than stabilizing effects, regardless of whether the reference growth was of plants in sterile or live soils (panel A). Among species pairs where microbes destabilize interactions, fitness differences were stronger than destabilizing effects only among plants whose growth in steriles soils provided the reference comparison (panel B). Large points indicate the overall effect sizes; black lines show the 95% confidence interval.

**Figure S.8:**
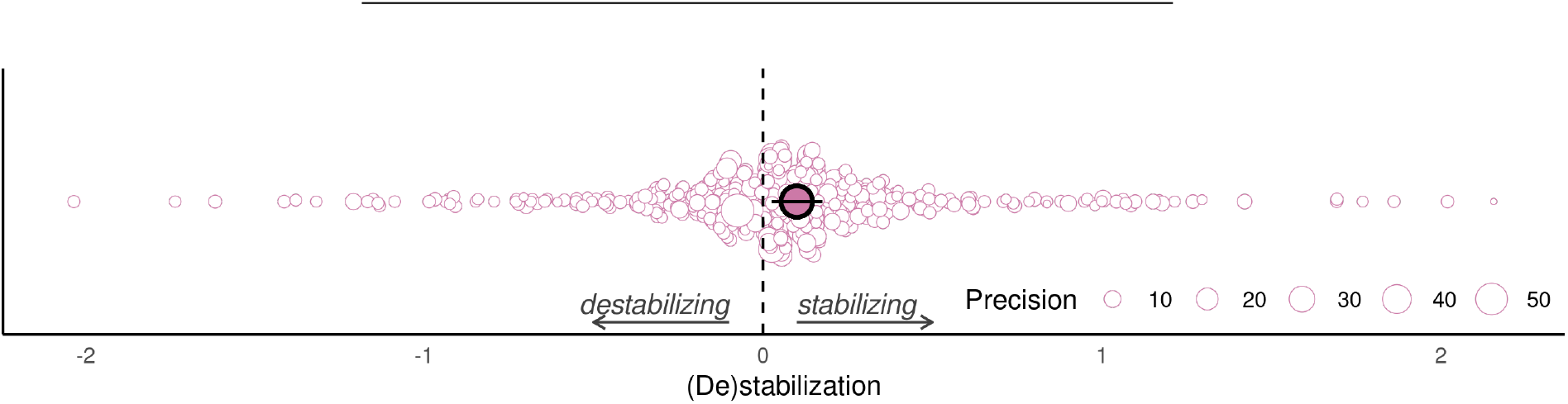
Orchard plot showing the results from a meta-analysis of the destabilizing and stabilizing microbial effects across the entire dataset.

**Table S1:**
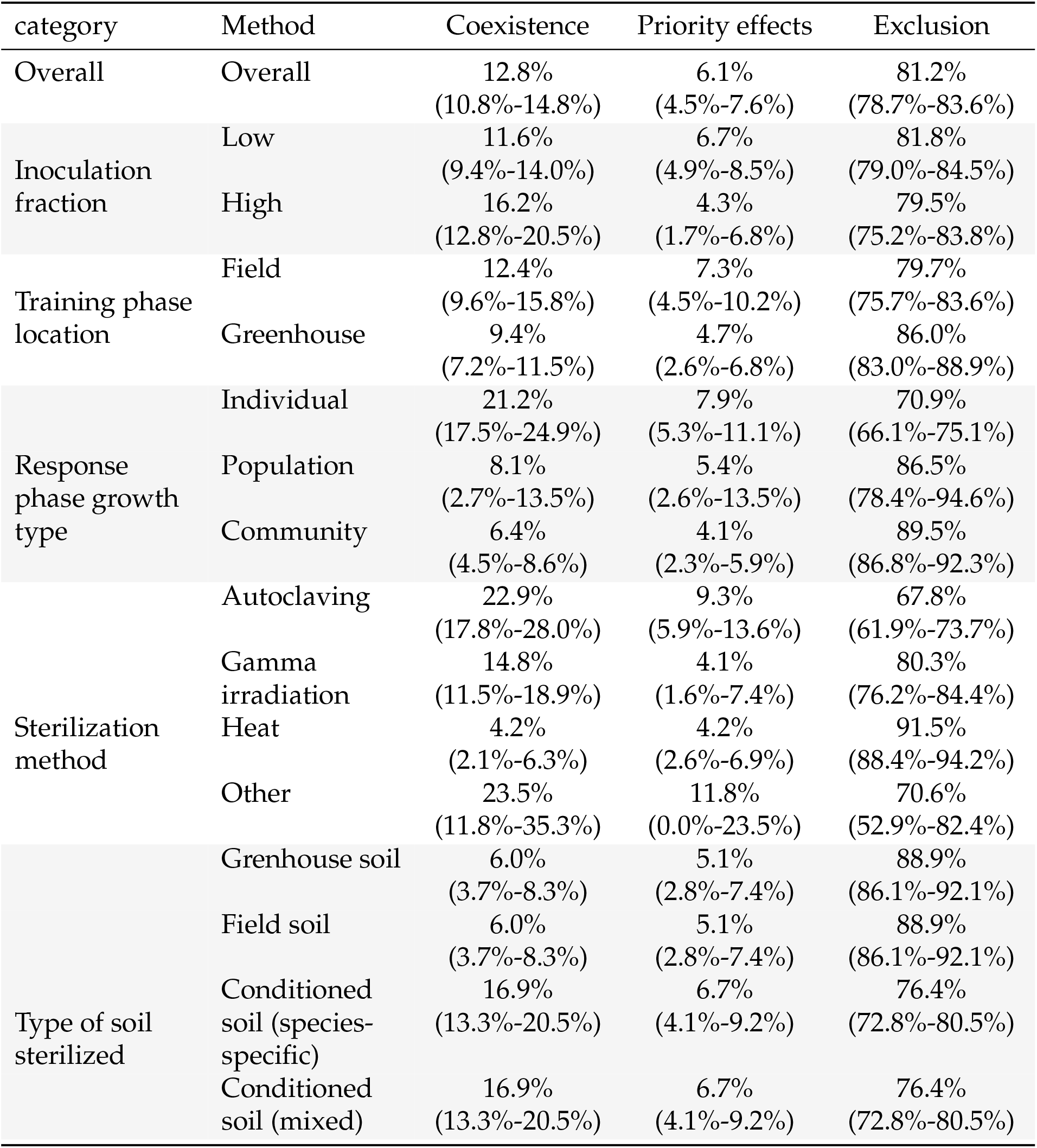
Proportion of simulation runs (Q2) that resulted in Coexistence/Priority effects/Exclusion among pairs for which growth in sterile soils provide the reference. Top number indicates median, and the range indicates the 95% confidence interval.

**Table S2:**
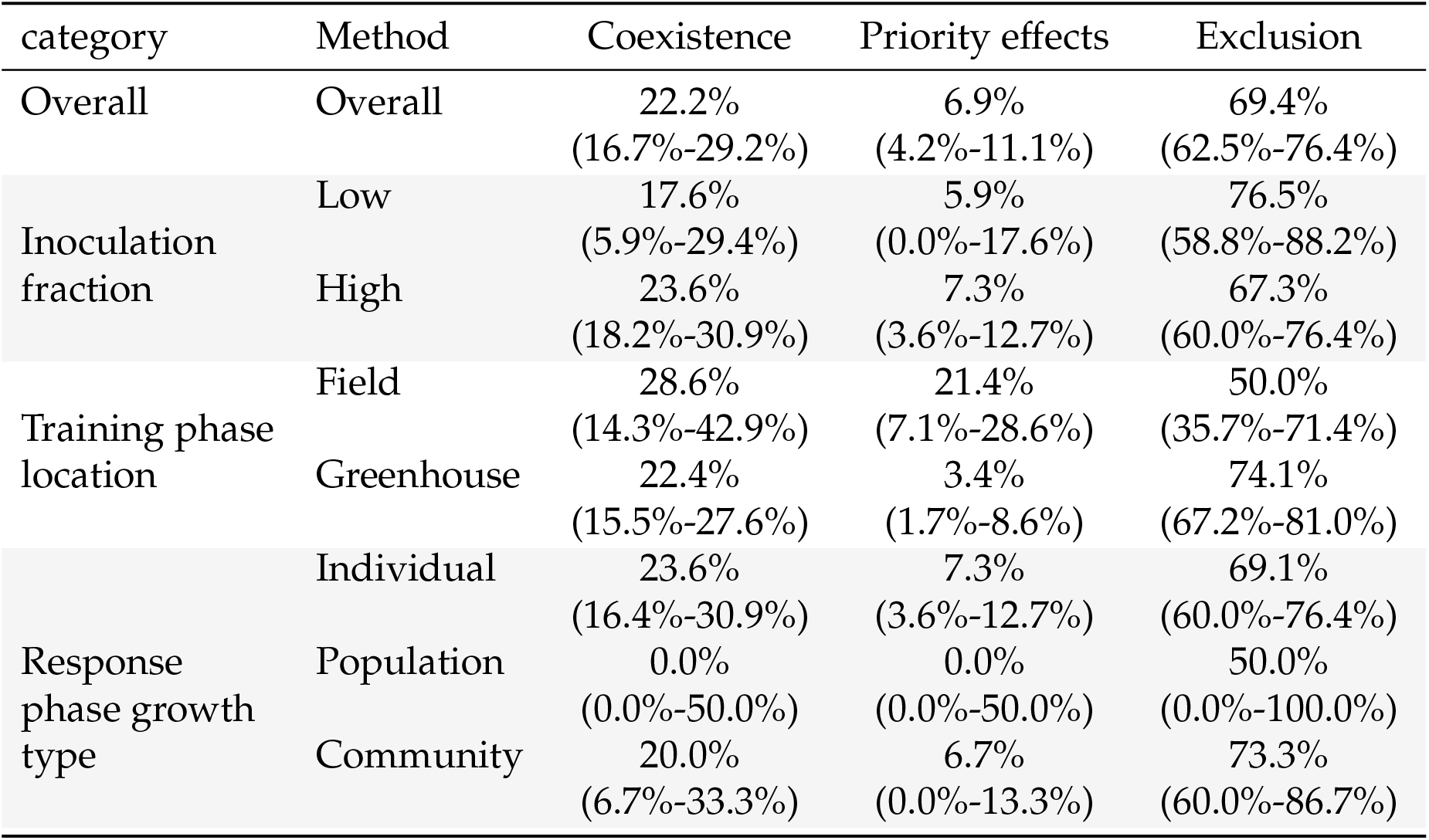
Proportion of simulation runs (Q2) that resulted in Coexistence/Priority effects/Exclusion among pairs for which growth in uncultivated live soils provide the reference. Top number indicates median, and the range indicates the 95% confidence interval.

## Notes

### Competing Interest Statement

The authors have declared no competing interest.

### Summary of Updates

Corrected a mistake in Fig. 3 legend

