## Supplemental materials and code for "A quantitative synthesis of soil microbial effects on plant species coexistence": appendices.html

Supplementary information for ‘A quantitative synthesis of soil microbial effects on plant species coexistence’


Code 

- Show All Code
- Hide All Code

### Supplementary information for ‘A quantitative synthesis of soil microbial effects on plant species coexistence’

###### Xinyi Yan, Jonathan Levine, and Gaurav Kandlikar

###### Last updated 10 Nov 2021

### Appendix 1: PRISMA chart

PRISMA chart

From: Page MJ, McKenzie JE, Bossuyt PM, Boutron I, Hoffmann TC, Mulrow CD, et al. The PRISMA 2020 statement: an updated guideline for reporting systematic reviews. BMJ 2021;372:n71. doi: 10.1136/bmj.n71. For more information, visit: http://www.prisma-statement.org/

---


---

### Appendix 2: Derivation of effect sizes and variance terms

Here we show the derivation of the microabilly mediated (de)stabilization and fitness difference effect sizes and their variances.

##### Effect sizes: (de)stabilization & fitness differences

Following (1), the (de)stabilization and fitness difference of plant species 1 and 2 mediated by their microbial communities A and B can be expressed as the following: \[\begin{equation}
\mathrm{(De)stabilization} = -\frac{1}{2} (m\_{1A}-m\_{1B}-m\_{2A}+m\_{2B}) \end{equation}\] \[\begin{equation}
\mathrm{Fitness\ Difference} = \frac{1}{2}[(m\_{1A}+m\_{1B})-(m\_{2A}+m\_{2B})] \end{equation}\]

By operationalizing each microbial effect (\(m\_{iX}\)) as the natural log growth of plant species \(i\) on microbial community \(X\) mimus its log growth in a reference soil community (\(\ln(G\_{iX})-\ln(G\_{iR})\)), the above equations can be expanded. The reference growth terms cancel out when calculating the (de)stabilization, but not for the fitness difference:

\[\begin{equation}
\begin{aligned}
\mathrm{(De)stabilization} &= -\frac{1}{2}
[(\ln(G\_{1A})-\ln(G\_{1R}))-(\ln(G\_{1B})-\ln(G\_{1R}))-\\
&\ \ \ \ \ \ \ \ \ \ \ \ \ (\ln(G\_{2A})-\ln(G\_{2R}))+(\ln(G\_{2B})-\ln(G\_{2R}))]\\
&= -\frac{1}{2} [\ln(G\_{1A})-\ln(G\_{1B})-\ln(G\_{2A})+\ln(G\_{2B})]
\end{aligned}
\end{equation}\]

\[\begin{equation}
\begin{aligned}
\mathrm{Fitness\ Difference} &= \frac{1}{2}
[(\ln(G\_{1A})-\ln(G\_{1R}))+(\ln(G\_{1B})-\ln(G\_{1R}))-\\
&\ \ \ \ \ \ \ \ \ \ (\ln(G\_{2A})-\ln(G\_{2R}))-(\ln(G\_{2B})-\ln(G\_{2R}))]\\
&= \frac{1}{2}[\ln(G\_{1A})+\ln(G\_{1B})-\ln(G\_{2A})-\ln(G\_{2B})]-\ln(G\_{1R})+\ln(G\_{2R})
\end{aligned}
\end{equation}\]

Both summations can also be written in the log response ratio form:

\[\begin{equation}
\begin{aligned}
\mathrm{(De)stabilization} &= -\frac{1}{2} [\ln(G\_{1A})-\ln(G\_{1B})-\ln(G\_{2A})+\ln(G\_{2B})]\\
&= -\frac{1}{2}[(\ln(\frac{G\_{1A}}{G\_{1B}})+\ln(\frac{G\_{2B}}{G\_{2A}})]\\
&= -\frac{1}{2}\ln(\frac{(G\_{1A}G\_{2B}}{G\_{1B}G\_{2A}})
\end{aligned}
\end{equation}\]

\[\begin{equation}
\begin{aligned}
\mathrm{Fitness\ Difference} &= \frac{1}{2}
[(\ln(G\_{1A})-\ln(G\_{1R}))+(\ln(G\_{1B})-\ln(G\_{1R})) -
(\ln(G\_{2A})-\ln(G\_{2R}))-(\ln(G\_{2B})-\ln(G\_{2R}))]\\
&= \frac{1}{2}[\ln(\frac{G\_{1A}}{G\_{1R}})+\ln(\frac{G\_{1B}}{G\_{1R}})-\ln(\frac{G\_{2A}}{G\_{2R}})-\ln(\frac{G\_{2B}}{G\_{2R}})]\\
&= \frac{1}{2}\ln(\frac{\frac{G\_{1A}}{G\_{1R}}\frac{G\_{1B}}{G\_{1R}}} {\frac{G\_{2A}}{G\_{2R}}\frac{G\_{2B}}{G\_{2R}}} )
\end{aligned}
\end{equation}\]

Note that the \(\mathrm{rr((De)stabilization)}\) is \(-\frac{1}{2}\) of the \({rr(I\_{S})}\) effect size used by Crawford et a., and can likewise be interpreted as the difference of the two plant’s species in their responses to conspecific and heterospecific soils. The \(\mathrm{rr(Fitness\ Difference)}\) is the ratio of the average mictobially mediated fitness of plant species 1 over that of plant species 2.

##### Variance of the (de)stabilization & fitness differences

The variance of the effect sizes can be calculated using the following equation ((2), (3)):  
\[\begin{equation}
\sigma^2(\mathrm{(De)stabilization/Fitness\ Difference}) = A^{T}\Sigma A
\end{equation}\]

where \(A\) is the column vector of partial derivatives of \(\mathrm{(de)stabilization}\) or \(\mathrm{Fitness\ Difference}\) (FD) with respect to each of its \(G\) terms.

Therefore,  
\[\begin{equation}
\begin{aligned} A\_{\mathrm{(de)st}}^T &= \frac{1}{2}[\frac{\partial (de)st}{\partial G\_{1A}},\ \frac{\partial (de)st}{\partial G\_{1B}},\ \frac{\partial FD}{\partial G\_{2A}},\ \frac{\partial FD}{\partial G\_{2B}}]\\
&= [\frac{1}{G\_1A},\ -\frac{1}{G\_1B},\ -\frac{1}{G\_2A},\ \frac{1}{G\_2R},\ \frac{1}{G\_2B}]
\end{aligned}
\end{equation}\]

\[\begin{equation}
\begin{aligned} A\_{\mathrm{FD}}^T &= \frac{1}{2}[\frac{\partial FD}{\partial G\_{1A}},\ \frac{\partial FD}{\partial G\_{1R}},\ \frac{\partial FD}{\partial G\_{1B}},\ \frac{\partial FD}{\partial G\_{1R}},\ ... ,\ \frac{\partial FD}{\partial G\_{2B}}]\\
&= [\frac{1}{G\_1A},\ -\frac{1}{G\_1R},\ \frac{1}{G\_1B},\ -\frac{1}{G\_1R},\ -\frac{1}{G\_2A}, \frac{1}{G\_2R},\ -\frac{1}{G\_2B},\ \frac{1}{G\_2R} ]
\end{aligned}
\end{equation}\]

Assuming that the growth terms (\(G\_{iX}\)) are independent of each other, the large-sample variance covariance matrix for niche difference \(\Sigma\_{(de)st}\) is simply a diagonal matrix; However, due to the repeated terms for growth in reference soils (\(G\_{iR}\)) in the eqution for fitness difference, \(\Sigma\_{FD}\) contains covariance terms between the duplicated controls (see (3) Appendix Fig. A1 (b)):

\(\Sigma\_{\mathrm{SD}} = \begin{bmatrix} \frac{\sigma\_{1A}^2}{N\_{1A}} & 0 & 0 & 0\\ 0 & \frac{\sigma\_{1B}^2}{N\_{1B}} & 0 & 0\\ 0 & 0 & \frac{\sigma\_{2A}^2}{N\_{2A}} & 0 \\ 0 & 0 & 0 & \frac{\sigma\_{2B}^2}{N\_{2B}}\\ \end{bmatrix}\), \(\Sigma\_{\mathrm{FD}} = \begin{bmatrix} \frac{\sigma\_{1A}^2}{N\_{1A}} & 0 & 0 & 0 & 0 & 0 & 0 & 0\\ 0 & \frac{\sigma\_{1R}^2}{N\_{1R}} & 0 & \frac{\sigma\_{1R}^2}{N\_{1R}} & 0 & 0 & 0 & 0\\ 0 & 0 & \frac{\sigma\_{1B}^2}{N\_{1B}} & 0 & 0 & 0 & 0 & 0\\ 0 & \frac{\sigma\_{1R}^2}{N\_{1R}} & 0 & \frac{\sigma\_{1R}^2}{N\_{1R}} & 0 & 0 & 0 & 0\\ 0 & 0 & 0 & 0 & \frac{\sigma\_{2A}^2}{N\_{2A}} & 0 & 0 & 0 \\ 0 & 0 & 0 & 0 & 0 & \frac{\sigma\_{2R}^2}{N\_{2R}} & 0 & \frac{\sigma\_{2R}^2}{N\_{2R}}\\ 0 & 0 & 0 & 0 & 0 & 0 & \frac{\sigma\_{2B}^2}{N\_{2B}} & 0\\ 0 & 0 & 0 & 0 & 0 & \frac{\sigma\_{2R}^2}{N\_{2R}} & 0 & \frac{\sigma\_{2R}^2}{N\_{2R}}\\ \end{bmatrix}\)

where each \(\sigma\_{iX}\) represents the standard error of the mean for plant growth, and \(N\_{iX}\) represents the sample size.

Putting together, the respective variance for the (de)stabilization and fitness difference are: \[\begin{equation}
\sigma\_{\mathrm{(de)st}}^2 = \frac{1}{4}( \frac{\sigma\_{1A}^2}{G\_{1A}^2N\_{1A}} + \frac{\sigma\_{1B}^2}{G\_{1B}^2N\_{1B}} + \frac{\sigma\_{2A}^2}{G\_{2A}^2N\_{2A}} + \frac{\sigma\_{2B}^2}{G\_{2B}^2N\_{2B}} )
\end{equation}\]

\[\begin{equation}
\sigma\_{\mathrm{FD}}^2 = \frac{1}{4}( \frac{\sigma\_{1A}^2}{G\_{1A}^2N\_{1A}} + \frac{\sigma\_{1B}^2}{G\_{1B}^2N\_{1B}} + 4\frac{\sigma\_{1R}^2}{G\_{1R}^2N\_{1R}} + \frac{\sigma\_{2A}^2}{G\_{2A}^2N\_{2A}} + \frac{\sigma\_{2B}^2}{G\_{2B}^2N\_{2B}} + 4\frac{\sigma\_{2R}^2}{G\_{2R}^2N\_{2R}} )
\end{equation}\]

Note that the above variance for (de)stabilization is \(\frac{1}{4}\) of that for \(I\_{S}\) used by Crawford et al. Those variances can also be calculated following the statistical definition of standard error of the mean. Here we show the derivation for fitness difference: \[\begin{equation}
\begin{aligned}
\sigma\_{\mathrm{FD}}^2 &= \sigma^2(\frac{1}{2}[\ln(G\_{1A}) + \ln(G\_{1B}) - 2\ln(G\_{1R}) - \ln(G\_{2A}) - \ln(G\_{2B}) + 2\ln(G\_{2R}) ])\\
&= \frac{1}{4}[\frac{\sigma^2(\ln(G\_{1A}))}{N\_{1A}} +\frac{\sigma^2(\ln(G\_{1B}))}{N\_{1B}} + 4\frac{\sigma^2(\ln(G\_{1R}))}{N\_{1R}} + \frac{\sigma^2(\ln(G\_{2A}))}{N\_{2A}} +\frac{\sigma^2(\ln(G\_{2B}))}{N\_{2B}} + 4\frac{\sigma^2(\ln(G\_{2R}))}{N\_{2R}}] \\
\\
&\text{ and using propagation error theory to get the variance of a natural log of a random variable:}\\
\\
&= \frac{1}{4}[\frac{\sigma^2(G\_{1A})}{N\_{1A}G\_{1A}^2} + \frac{\sigma^2(G\_{1B})}{N\_{1B}G\_{1B}^2} + 4\frac{\sigma^2(G\_{1R})}{N\_{1R}G\_{1R}^2} + \frac{\sigma^2(G\_{2A})}{N\_{2A}G\_{2A}^2} + \frac{\sigma^2(G\_{2B})}{N\_{2B}G\_{2B}^2} + 4\frac{\sigma^2(G\_{2R})}{N\_{2R}G\_{2R}^2}]]
\end{aligned}
\end{equation}\]

---


---

### Appendix 3: `R` code to recreate all results

#### Read in datasets

For this analysis, we are working with two complementary datasets. The first is the dataset of SD/FD effect sizes with field live soil as the reference; the second set of effect sizes are from studies with sterile soils as reference inoculua.

```
bind_rows(sterile_ref, live_ref, .id = "reference_type") %>%
  mutate(reference_type = ifelse(reference_type == 1, "sterile soil", "live soil")) %>%
  group_by(reference_type) %>%
  summarize(n_rows = n()) %>%
  kable() %>%
  kable_styling(full_width = FALSE)
```

| reference\_type | n\_rows |
| --- | --- |
| live soil | 72 |
| sterile soil | 449 |

Note that there are 449 comparisons in the sterile soil dataset right now, but there are three comparisons we will eventually need to exclude because of unreliable data, see below.

#### Explore data availability and patterns

We can first get a lay of the land on this dataset by checking how many species there are, from how many experiments, and from how many studies:

```
# Total number of effect sizes:
# FIElD LIVE REFERENCE
# first, across the whole dataset
cat(paste0("&emsp; Number of unique comparisons in Live ref dataset: ", 
           nrow(live_ref)))
```

  Number of unique comparisons in Live ref dataset: 72

```
# Number of unique species pairs
cat(paste0("&emsp; Number of unique species pairs in Live ref dataset: ", 
           live_ref$species_pair %>% unique %>% length))
```

  Number of unique species pairs in Live ref dataset: 69

```
# Number of unique Studies
cat(paste0("&emsp; Number of unique studies in Live ref dataset: ",
           live_ref$Study %>% unique %>% length))
```

  Number of unique studies in Live ref dataset: 16

```
# Number of unique Experiments
cat(paste0("&emsp; Number of unique experiments in Live ref dataset: ",
           live_ref$Experiment %>% unique %>% length))
```

  Number of unique experiments in Live ref dataset: 18

```
cat("----------")
```

---

```
# STERILE SOIL REFERENCE
# first, across the whole dataset

cat(paste0("&emsp; Number of unique comparisons in Sterile ref dataset: ", 
           nrow(sterile_ref)))
```

  Number of unique comparisons in Sterile ref dataset: 449

```
# Number of unique species pairs
cat(paste0("&emsp; Number of unique species pairs in Sterile ref dataset: ", 
           sterile_ref$species_pair %>% unique %>% length))
```

  Number of unique species pairs in Sterile ref dataset: 213

```
# Number of unique Studies
cat(paste0("&emsp; Number of unique studies in Sterile ref dataset: ",
           sterile_ref$Study %>% unique %>% length))
```

  Number of unique studies in Sterile ref dataset: 38

```
# Number of unique Experiments
cat(paste0("&emsp; Number of unique experiments in Sterile ref dataset: ",
           sterile_ref$Experiment %>% unique %>% length))
```

  Number of unique experiments in Sterile ref dataset: 67

We can also do some summaries of the 4-5 moderators we are going to use later on in the analysis. First for the **Live reference** dataset:

```
live_ref %>%
  group_by(`Training environment`) %>%
  summarize(n_rows = n()) %>%
  kable() %>% kable_styling(full_width = F)
```

| Training environment | n\_rows |
| --- | --- |
| Field | 14 |
| Lab | 58 |

```
live_ref %>%
  mutate(inoc_frac_binary = ifelse(`Fraction inoculum` < 0.251, "low", "high")) %>%
  group_by(inoc_frac_binary) %>%
  summarize(n_rows = n()) %>%
  kable() %>% kable_styling(full_width = F)
```

| inoc\_frac\_binary | n\_rows |
| --- | --- |
| high | 55 |
| low | 17 |

```
live_ref %>%
  group_by(`Testing community`) %>%
  summarize(n_rows = n()) %>%
  kable() %>% kable_styling(full_width = F)
```

| Testing community | n\_rows |
| --- | --- |
| Community | 15 |
| Individual | 55 |
| Population | 2 |

And next for the **Sterile reference** dataset:

```
sterile_ref %>%
  group_by(`Training environment`) %>%
  summarize(n_rows = n()) %>%
  kable() %>% kable_styling(full_width = F)
```

| Training environment | n\_rows |
| --- | --- |
| Field | 178 |
| Lab | 235 |
| LabField | 36 |

```
sterile_ref %>%
  mutate(inoc_frac_binary = ifelse(`Fraction inoculum` < 0.251, "low", "high")) %>%
  group_by(inoc_frac_binary) %>%
  summarize(n_rows = n()) %>%
  kable() %>% kable_styling(full_width = F)
```

| inoc\_frac\_binary | n\_rows |
| --- | --- |
| high | 118 |
| low | 331 |

```
sterile_ref %>%
  group_by(`Testing community`) %>%
  summarize(n_rows = n()) %>%
  kable() %>% kable_styling(full_width = F)
```

| Testing community | n\_rows |
| --- | --- |
| Community | 220 |
| Individual | 192 |
| Population | 37 |

```
cat("Note that in the following table, CmS = cultivated, mixed, sterile; CS = cultivated sterile, FS = field sterile soil, GS = greenhouse sterile soil")
```

Note that in the following table, CmS = cultivated, mixed, sterile; CS = cultivated sterile, FS = field sterile soil, GS = greenhouse sterile soil

```
sterile_ref %>%
  group_by(control_type) %>%
  summarize(n_rows = n()) %>%
  kable() %>% kable_styling(full_width = F)
```

| control\_type | n\_rows |
| --- | --- |
| CmS | 15 |
| CS | 197 |
| FS | 20 |
| GS | 217 |

```
sterile_ref %>%
  group_by(Study_sterilization_method) %>%
  summarize(n_rows = n()) %>%
  kable() %>% kable_styling(full_width = F)
```

| Study\_sterilization\_method | n\_rows |
| --- | --- |
| autoclaving | 120 |
| gamma irradiation | 122 |
| heating | 189 |
| not specified | 3 |
| pasteurization | 6 |
| steam-pasteurization | 1 |
| steaming | 8 |

Before starting on the meta-analysis models, we should do some more data exploration. First let us evaluate patterns of conspecific and heterospecific effects of soil microbes (boxplots of the conspecific effects of microbes (\(m\_{1A}\) and \(m\_{2B}\)) vs heterospecific effects (\(m\_{1B}\) and \(m\_{2A}\))):

```
live_ref %>%
  mutate(m1A = log(`AinA mean`) - log(`AinR mean`),
         m1B = log(`AinB mean`) - log(`AinR mean`),
         m2A = log(`BinA mean`) - log(`BinR mean`),
         m2B = log(`BinB mean`) - log(`BinR mean`)) %>%
  select(m1A:m2B) %>%
  pivot_longer(cols = m1A:m2B, names_to = "which_metric") %>%
  mutate(con_het = ifelse(which_metric %in% c("m1A", "m2B"), "con", "het")) %>%
  ggplot() +
  geom_violin(aes(y = value, x = con_het)) +
  labs(title = "Live ref") + 
  theme_classic()
```

```
# Slightly more positive (less negative) heterospecific effects than
# conspecific effects, consistent with stabilization

sterile_ref %>%
  mutate(m1A = log(`AinA mean`) - log(`AinR mean`),
         m1B = log(`AinB mean`) - log(`AinR mean`),
         m2A = log(`BinA mean`) - log(`BinR mean`),
         m2B = log(`BinB mean`) - log(`BinR mean`)) %>%
  select(m1A:m2B) %>%
  pivot_longer(cols = m1A:m2B, names_to = "which_metric") %>%
  mutate(con_het = ifelse(which_metric %in% c("m1A", "m2B"), "con", "het")) %>%
  ggplot() +
  geom_violin(aes(y = value, x = con_het)) +
  labs(title = "Sterile ref") + 
  theme_classic()
```

```
## Warning: Removed 2 rows containing non-finite values (stat_ydensity).
```

There is one species pair with an extreme outlier in the heterospecific effects column. We need to look into what is happening here. I am fairly sure that this is the pair from McCarthy Neumann’s study, where the paper has no value for plant growth, which was interpreted as zero growth in Crawford et al’s dataset.

```
sterile_ref %>%
  mutate(m1A = log(`AinA mean`) - log(`AinR mean`),
         m1B = log(`AinB mean`) - log(`AinR mean`),
         m2A = log(`BinA mean`) - log(`BinR mean`),
         m2B = log(`BinB mean`) - log(`BinR mean`)) %>%
   select(species_pair, `Pairwise comparison`,
          `AinA mean`,`AinB mean`, `AinR mean`, `BinA mean`, `BinB mean`, `BinR mean`,
          m1A:m2B) %>%
  arrange(m2A) # %>% View
```

```
## # A tibble: 449 × 12
##    species_pair `Pairwise compa… `AinA mean` `AinB mean` `AinR mean` `BinA mean`
##    <chr>                   <dbl>       <dbl>       <dbl>       <dbl>       <dbl>
##  1 colubrina_s…              133      33.8        20.1        32.9        0.001 
##  2 banksia_lep…               NA      10.7         9.77       22.8        0.631 
##  3 leucanthemu…              182       0.015       0.096       0.321      0.025 
##  4 anthoxanthu…              177       0.027       0.025       0.233      0.035 
##  5 festuca_rub…               NA       0.119       0.131       0.529      0.0893
##  6 lepidium_ca…               NA       0.121       0.098       0.729      0.197 
##  7 berteroa_in…               NA       0.116       0.352       0.374      0.401 
##  8 acacia_spat…               NA       2.60        2.67        6.42       0.38  
##  9 acacia_spat…               NA       2.60        2.82        6.42       1.41  
## 10 calothamnus…               NA       1.60        2.62       11.1        1.05  
## # … with 439 more rows, and 6 more variables: BinB mean <dbl>, BinR mean <dbl>,
## #   m1A <dbl>, m1B <dbl>, m2A <dbl>, m2B <dbl>
```

That does appear to be the case, the pair with the really low value is `colubrina_spinosa_iriartea_deltoidea`, for which no data for one species as interpreted as 0 biomass. From personal conversations with the author of the original study, this was just “no data” likely because of bad seeds rather than microbial effects.

There is also one species pair (`rumex_occidentalis_rumex_salicifolius`) for which there is always zero growth in reference soil, this is not usable because the value for SE is zero, making it impossible to use in the meta-analysis. We can remove those points here.

```
sterile_ref <- 
  sterile_ref %>% 
  filter(!(species_pair %in% c("colubrina_spinosa_iriartea_deltoidea",
                                 "rumex_occidentalis_rumex_salicifolius")))
```

Removing those unusable data points leaves us with **446** pairwise comparisons in the Sterile Ref dataset.

#### Question 1

Now that the data look in good shape, we are ready to quantify the effect sizes ((de)stabilization and fitness differences). To do so we use functions saved in the file `functions_effectSizes_coexOutcomes.R`.

```
# Source the functions
source("analysis/functions_effectSizes_coexOutcomes.R")
# The function makes a new dataframe, so let's use that to override the
# existing one.
live_ref <- calculate_effect_sizes(live_ref)
# The new columns are: 
# mean_SD, IS, var_SD, sem_SD, n_min_SD; mean_FD, var_FD, sem_FD,
# n_min_FD; abs_FD, mean_IGR, var_IGR, sem_IGR

sterile_ref <- calculate_effect_sizes(sterile_ref)
```

##### Prepare the datasets for meta-analysis

Right now the datasets are in “wide” format, with each row containing a species pair with it’s stabilization and fitness difference measures, and we need to make this into a long format dataframe with each species pair having two rows (one for stabilization, one for fitness differences).

```
# Make a dataset in 'long' format with all effect sizes
# in one column, a moderator column that specifies whether it is FD/SD, 
live_ref_long <-
  live_ref %>%
  select(Experiment, Study, species_pair, author,
         mean_SD, mean_FD, var_SD, var_FD, 
         `Fraction inoculum`, `Training environment`, `Testing environment`,
         Study_sterilization_method, `Testing community`) %>%
  # rename(FD_mean = mean_FD, #absFD_mean = abs(FD_mean),
  #        FD_var = var_FD, SD_mean = mean_SD, SD_var = var_SD) %>%
  pivot_longer(cols = mean_SD:var_FD, 
               names_to = c(".value","set"),
               names_pattern = "(.+)_(.+)")  %>%
  rename(mean_value = mean, var_value = var) %>%
  mutate(
    # Make a new column with absolute value of FD, and `real` value of SD
    absolute_FD_effect = ifelse(set == "FD", abs(mean_value), mean_value),
    # Make a new column with absolute value of both SD and FD
    absolute_FDSD_effects = abs(mean_value),
    # Make a new column with just effect size ID (to act as ranef)
    ES_ID = 1:nrow(.),
    # Make a new column for each species pair within a study
    Experiment_SpPair = paste0(Experiment, "_", species_pair)
  ) 


# Do the same process for the sterile reference dataset


# Make a dataset in 'long' format with all effect sizes
# in one column, a moderator column that specifies whether it is FD/SD, 

sterile_ref_long <-
  sterile_ref %>%
  select(Experiment, Study, author, species_pair,
         mean_SD, mean_FD, var_SD, var_FD, 
         `Fraction inoculum`, `Training environment`, `Testing environment`, control_type,
         Study_sterilization_method, `Testing community`) %>%
  # rename(FD_mean = mean_FD, #absFD_mean = abs(FD_mean),
  #        FD_var = var_FD, SD_mean = mean_SD, SD_var = var_SD) %>%
  pivot_longer(cols = mean_SD:var_FD, 
               names_to = c(".value","set"),
               names_pattern = "(.+)_(.+)")  %>%
  rename(mean_value = mean, var_value = var) %>%
  mutate(
    # Make a new column with absolute value of FD, and `real` value of SD
    absolute_FD_effect = ifelse(set == "FD", abs(mean_value), mean_value),
    # Make a new column with absolute value of both SD and FD
    absolute_FDSD_effects = abs(mean_value),
    # Make a new column with just effect size ID (to act as ranef)
    ES_ID = 1:nrow(.),
    # Make a new column for each species pair within a study
    Experiment_SpPair = paste0(Experiment, "_", species_pair)
  )
```

##### Run core models

For the first question, we need to run a multi-variate meta-analysis model. To do this we first need to define a new function, because otherwise we would have to keep repeating a bunch of code.

###### Define a function to run the meta-model

```
run_metamod <- function(which_df, which_response_column, 
                        moderators = " ~ (set-1)", impute_cov_r) {
  
  # Impute a variance-covariance matrix with assumed
  # correlation between effect sizes = impute_cov_r
  # This uses a function from clubSandwich
  imputed_Vlist <- impute_covariance_matrix(vi = which_df$var_value, 
                                            cluster = which_df$Experiment,
                                            subgroup = which_df$species_pair,
                                            r = impute_cov_r, return_list = F)
  
  # Run the model
  rma_formula <- formula(paste(which_response_column, moderators))
  model_out <- rma.mv(rma_formula,
                      V = imputed_Vlist,
                      random = list(~ set|Experiment_SpPair, # This is same as ES_ID
                                    ~ set|Experiment
                                    ), 
                      test = "t",
                      data = which_df, 
                      # allow for heterogeneity to differ between SD/FD
                      struct = "UN")
  return(model_out)
}
```

##### Run meta-analysis for the Live Reference dataset

With the function defined, we can now run the meta-analysis model on the live reference dataset.

```
# Run the meta-analysis model on the full live reference dataset ----
q1.lr.0.5 <- run_metamod(which_df = live_ref_long, 
                         which_response_column = "absolute_FDSD_effects",
                         moderators = " ~ (set-1)",
                         impute_cov_r = 0.5)

coef_test(q1.lr.0.5, vcov = "CR2") %>% # check if different from zero
  kable () %>%
  kable_styling(full_width = F)
```

|  | beta | SE | tstat | df | p\_Satt |
| --- | --- | --- | --- | --- | --- |
| setFD | 0.6308658 | 0.0909004 | 6.940190 | 14.07145 | 0.0000067 |
| setSD | 0.3638120 | 0.0721101 | 5.045232 | 13.98695 | 0.0001794 |

```
conf_int(q1.lr.0.5, vcov = "CR2") %>% # take look at the confidence interval
  kable () %>%
  kable_styling(full_width = F)
```

|  | beta | SE | df | CI\_L | CI\_U |
| --- | --- | --- | --- | --- | --- |
| setFD | 0.6308658 | 0.0909004 | 14.07145 | 0.4359967 | 0.8257348 |
| setSD | 0.3638120 | 0.0721101 | 13.98695 | 0.2091378 | 0.5184863 |

```
q1.lr.0.5_coefs <- data.frame(conf_int(q1.lr.0.5, vcov =  "CR2"))
# See https://cran.r-project.org/web/packages/clubSandwich/vignettes/Wald-tests-in-clubSandwich.html
# for explainer of Wald_test()
# (basically, check if values are different from one another)
q1.lr.wald <- Wald_test(q1.lr.0.5, constraints = constrain_equal(1:2), vcov = "CR2")


# Check the estimated correlation: 
# cov2cor(q1.lr.0.5$vb)

# Uncomment the following lines to re-run the profile plots -- commenting
# now because it's quite a slow step. 

# pdf("figures/q1-lr-profile.pdf", height = 8, width = 8)
# par(mfrow = c(2,2))
# profile(q1.lr.0.5)
# dev.off()
```

Now we can quantify the heterogeneity.

```
# To quantify heterogeneity

# Make an equivalent model with no random effects
# this is needed to calculate I^2_R, following
# Jackson et al. 2012. See also:
# http://www.metafor-project.org/doku.php/tips:i2_multilevel_multivariate
V_mat_lr <- impute_covariance_matrix(
  vi = live_ref_long$var_value,
  cluster = live_ref_long$Experiment,
  subgroup = live_ref_long$species_pair,
  r = 0.5, return_list = F)
V_mat_lr <- bldiag(V_mat_lr)

q1.lr.0.5.noRanefs <- rma.mv(absolute_FDSD_effects ~ (set - 1), V = V_mat_lr, data = live_ref_long)

I2R_LiveRef <- 
  c(100 * (vcov(q1.lr.0.5)[1,1] - vcov(q1.lr.0.5.noRanefs)[1,1]) / vcov(q1.lr.0.5)[1,1],
    100 * (vcov(q1.lr.0.5)[2,2] - vcov(q1.lr.0.5.noRanefs)[2,2]) / vcov(q1.lr.0.5)[2,2])

I2R_LiveRef # First for FD, then stabilization
```

```
## [1] 89.72426 93.66488
```

##### Run meta-analysis for the Sterile Reference dataset

```
# Run the meta-analysis model on the full sterile reference dataset ----
q1.xs.0.5 <- run_metamod(which_df = sterile_ref_long, 
                         which_response_column = "absolute_FDSD_effects", 
                         moderators = " ~ (set-1)",
                         impute_cov_r = 0.5)

q1.xs.coef <- coef_test(q1.xs.0.5, vcov = "CR2") # check if different from zero
q1.xs.0.5_coefs <- data.frame(conf_int(q1.xs.0.5, vcov =  "CR2"))
# conf_int(q1.xs.0.5, vcov = "CR2") # check if different from zero
q1.xs.wald <- Wald_test(q1.xs.0.5, constraints = constrain_equal(1:2), vcov = "CR2") # check if values are different from one another
cov2cor(q1.xs.0.5$vb)
```

```
##           setFD     setSD
## setFD 1.0000000 0.3909211
## setSD 0.3909211 1.0000000
```

```
# pdf("figures/q1-xs-profile.pdf", height = 8, width = 8)
# par(mfrow = c(2,2))
# profile(q1.xs.0.5)
# dev.off()
```

Now, quantify heterogeneity

```
# Quantify heterogeneity 
V_mat_xs <- impute_covariance_matrix(
  vi = sterile_ref_long$var_value,
  cluster = sterile_ref_long$Experiment,
  subgroup = sterile_ref_long$species_pair,
  r = 0.5, return_list = F)
V_mat_xs <- bldiag(V_mat_xs)

q1.xs.0.5.noRanefs <- rma.mv(absolute_FDSD_effects ~ (set - 1), V = V_mat_xs, data = sterile_ref_long)

I2R_SterileRef <- 
  c(100 * (vcov(q1.xs.0.5)[1,1] - vcov(q1.xs.0.5.noRanefs)[1,1]) / vcov(q1.xs.0.5)[1,1],
    100 * (vcov(q1.xs.0.5)[2,2] - vcov(q1.xs.0.5.noRanefs)[2,2]) / vcov(q1.xs.0.5)[2,2])
I2R_SterileRef
```

```
## [1] 98.94877 94.30221
```

###### Make plot of the effect sizes

```
q1_dfs <- bind_rows(q1.xs.0.5_coefs,
                    q1.lr.0.5_coefs)

q1_dfs <- q1_dfs %>%
  mutate(yval = c(1.2, 0.8, 2.2, 1.8),
         which_source = rep(c("XS", "LR"), each = 2),
         effect_type = rep(c("Fitness difference", 
                             "(De)stabilization"), 2),
         I2R = c(I2R_SterileRef, I2R_LiveRef))
prediction_intervals <- bind_rows(data.frame(predict.rma(q1.xs.0.5)[1:2,]),
                                  data.frame(predict.rma(q1.lr.0.5)[1:2,])) %>%
  mutate(which_source = rep(c("XS", "LR"), each = 2),
         effect_type = c("(De)stabilization", "Fitness difference", "(De)stabilization", "Fitness difference"))

q1_dfs <- left_join(x = q1_dfs, y = prediction_intervals, 
                    by = c("effect_type", "which_source"))

# add a new column to the long datasets to capture the "scale" 
# (1/precision) for each effect size
sterile_ref_long <-
  sterile_ref_long %>%
  mutate(scale = 1/sqrt(var_value),
         yval = ifelse(set == "FD", 1.2,0.8)) %>%
  mutate(effect_type = ifelse(set == "FD", "Fitness difference", "(De)stabilization"))


live_ref_long <-
  live_ref_long %>%
  mutate(scale = 1/sqrt(var_value),
         yval = ifelse(set == "FD", 2.2,1.8)) %>%
  mutate(effect_type = ifelse(set == "FD", "Frequency independent", "Frequency dependent"))

(q1_plot <- 
  ggplot(q1_dfs) +
  ggbeeswarm::geom_quasirandom(data = sterile_ref_long,
                               aes(y = absolute_FDSD_effects, x = yval, size = scale, color = effect_type),
                               alpha = 1, pch = 21, fill = "transparent", stroke = .25) +
  ggbeeswarm::geom_quasirandom(data = live_ref_long,
                               aes(y = absolute_FDSD_effects, x = yval, size = scale, color = effect_type),
                               alpha = 1, pch = 21, fill = "transparent", stroke = .25) +
  geom_point(aes(y = beta, x = yval, fill = effect_type),
             size = 5, stroke = 1.5, shape = 21) + 
  geom_errorbar(aes(x = yval, ymin = CI_L, ymax = CI_U), width = 0, size = 0.8) +
  # geom_errorbar(aes(x = yval, ymin = pi.lb, ymax = pi.ub), width = 0, size = 0.15) +
  geom_vline(xintercept = 1.5, size = .25) + 
  geom_hline(yintercept = 0, linetype = 2, size = .5, color = "grey25") +
  
  annotate("text", y = 4.5, x = 2, label = paste0("K = ", nrow(live_ref_long)/2, 
                                                  " species pairs\n (",length(unique(live_ref_long$Experiment))," experiments)"),
           hjust = 1, size = 4) + 
  annotate("text", y = 4.5, x = 1, label = paste0("K = ", nrow(sterile_ref_long)/2, 
                                                  " species pairs\n (",length(unique(sterile_ref_long$Experiment))," experiments)"), 
           hjust = 1, size = 4) + 
  scale_fill_manual(name = "", values = rev(c(alpha("#009E73",0.6), alpha("#CC79A7",0.75)))) +
  scale_color_manual(name = "", values = rev(c("#009E73", "#CC79A7", "#009E73","#CC79A7"))) +
  scale_x_continuous(name = "", breaks = c(1,2), labels = c("Sterile\nreference\nsoil", "Live\nreference\nsoil")) +
  scale_y_continuous(limits = c(-1.5, 5.15), breaks = c(0, 2,4)) +
  scale_size_continuous(name = "Precision (1/SE)", breaks = c(10, 30, 50)) + 
  ylab("Effect size") + 
  theme_plots() +

  annotate("segment", y = 2.8, yend = Inf, x = c(1.5, 1.7), xend = c(1.5,1.7), size = .25) +
  annotate("segment", y = c(2.8, Inf), yend = c(2.8, Inf), x = 1.5, xend = 1.7, size = .25)  + 
    
  geom_point(aes(y = 3, x = 1.65), size = 4, shape = 21, fill = c("#009E73")) + 
  geom_point(aes(y = 3, x = 1.55), size = 4, shape = 21, fill = c("#CC79A7")) + 
  annotate("text", y = 3.2, x = 1.65, label = "Fitness difference", hjust = 0, size = 4) +
  annotate("text", y = 3.2, x = 1.55, label = "(De)stabilization", hjust = 0, size = 4) +  
  
  ggtext::geom_richtext(aes(y = -0.75, x = yval,
                            label = paste0("I<sub>R</sub><sup>2</sup> = ", round(I2R, 1))),
                        size = 3.5, vjust = -0.25, label.color = NA) +
  guides(fill = "none", color = "none", text = "none") + 
  
  ggpubr::geom_bracket(xmin = 0.8, xmax = 1.2, y = 5.14, 
                       label = "***", tip.length = c(0.01, 0.01)) +
  ggpubr::geom_bracket(xmin = 1.8, xmax = 2.2, y = 2.3, 
                       label = "***", tip.length = c(0.01, 0.01)) +
  
  theme(axis.text.y = element_text(color = "black", size = 13),
        axis.text.x = element_text(size = 12),
        legend.position= c(1, 0), 
        legend.direction = "horizontal",
        legend.justification = c(1, 0),
        plot.title.position = "plot",
        plot.title = element_text(face = "bold")) +
  coord_flip())
```

```
if(save_figures) {
  ggsave("figures/q1_figure.png",q1_plot, bg = "white", width = 6.5, height = 4.5)
  ggsave("figures/q1_figure.pdf",q1_plot, bg = "white", width = 6.5, height = 4.5)
}
```

##### Check sensitivity (Influence analysis)

```
# We can run the Cook's distances to identify "influential" experiments
cd.exp.q1.lr.0.5 <- cooks.distance.rma.mv(q1.lr.0.5, 
                                          cluster = live_ref_long$Experiment, 
                                          parallel = "multicore")
21

# identify which study  is the influential one
which(cd.exp.q1.lr.0.5 > 0.45) # study 143
live_ref_long[live_ref_long$Experiment == 143,] # Kandlikar 2021

#
plot(cd.exp.q1.lr.0.5, type="o", pch=19, xlab="Experiment", ylab="Cook's Distance")

abline(0.45, 0, col = "red")

# re-do analysis without study 143
live_ref_trim <- live_ref_long %>% filter(Experiment != 143)
q1.lr.0.5.trim <- run_metamod(which_df = live_ref_trim, 
                              which_response_column = "absolute_FDSD_effects",
                              moderators = " ~ (set-1)",
                              impute_cov_r = 0.5)

cooks.distance.rma.mv(q1.lr.0.5.trim, 
                      cluster = live_ref_trim$Experiment, 
                      parallel = "multicore")
live_ref_long[live_ref_long$Experiment == 138,] # detect another influential exp

# delete this other influential exp.:
live_ref_trim <- live_ref_long %>% 
  filter(Experiment != 138 & Experiment != 143)
q1.lr.0.5.trim <- run_metamod(which_df = live_ref_trim, 
                         which_response_column = "absolute_FDSD_effects",
                         moderators = " ~ (set-1)",
                         impute_cov_r = 0.5)
cooks.distance.rma.mv(q1.lr.0.5.trim, 
                      cluster = live_ref_trim$Experiment, 
                      parallel = "multicore")

coef_test(q1.lr.0.5.trim, vcov = "CR2") # check if different from zero
conf_int(q1.lr.0.5.trim, vcov = "CR2") # take look at the confidence interval
q1.lr.trim.wald <- Wald_test(q1.lr.0.5.trim, constraints = constrain_equal(1:2), vcov = "CR2")


# find "influential points"
cd.points.q1.lr.0.5 <- cooks.distance.rma.mv(q1.lr.0.5, parallel = "multicore")
which(cd.points.q1.lr.0.5>0.45)

plot(cd.points.q1.lr.0.5, type = "o", pch = 19, xlab = "Points", ylab = "Cook's Distance")
abline(0.45, 0, col = "red")


# Now for sterile soils -------

# influence test
# find "influential experiments"
cd.exp.q1.xs.0.5<-cooks.distance.rma.mv(q1.xs.0.5, 
                      cluster = sterile_ref_long$Experiment, #whereas influential studies = 51, 172
                      reestimate = F,
                      parallel = "multicore")

which(cd.exp.q1.xs.0.5>0.45) # influential experiment = 33
sterile_ref_long[sterile_ref_long$Experiment ==33,] %>% view()

plot(cd.exp.q1.xs.0.5, type="o", pch=19, xlab="Experiment", ylab="Cook's Distance")
abline(0.45, 0, col = "red")

# remove influential experiment 33
sterile_ref_trim <- sterile_ref_long %>% filter(Experiment != 33)
q1.xs.0.5.trim <- run_metamod(which_df = sterile_ref_trim, 
                         which_response_column = "absolute_FDSD_effects",
                         moderators = " ~ (set-1)",
                         impute_cov_r = 0.5)
# check if there are new influence experiments:
cooks.distance.rma.mv(q1.xs.0.5.trim, 
                      cluster = sterile_ref_trim$Experiment, 
                      reestimate = F,
                      parallel = "multicore") # no other influence experiments

coef_test(q1.xs.0.5.trim, vcov = "CR2") # check if different from zero
conf_int(q1.xs.0.5.trim, vcov = "CR2") # take look at the confidence interval
q1.xs.trim.wald <- Wald_test(q1.xs.0.5.trim, constraints = constrain_equal(1:2), vcov = "CR2")


# find "influential points"
cd.points.q1.xs.0.5 <- cooks.distance.rma.mv(q1.xs.0.5, parallel = "multicore",
                                           reestimate = F)
which(cd.points.q1.xs.0.5 > 0.45) # no influential points

plot(cd.points.q1.xs.0.5, type = "o", pch = 19, xlab = "Points", ylab = "Cook's Distance")
abline(0.45, 0, col = "red")

cd.exp.q1.xs.0.5 <- cooks.distance.rma.mv(q1.xs.0.5, 
                                          cluster = sterile_ref_long$Experiment, #whereas influential studies = 51, 172
                                          reestimate = F,
                                          parallel = "multicore")
cd.points.q1.xs.0.5 <- cooks.distance.rma.mv(q1.xs.0.5, parallel = "multicore",
                                           reestimate = F)
```

###### Live reference influence analysis

In the influence analysis for live soil references we found two studies that were influential; we replicated all analyses without these studies (expand code chunk below for details).

```
# influence test
# find "influential experiments"
cd.exp.q1.lr.0.5 <- cooks.distance.rma.mv(q1.lr.0.5, 
                                          cluster = live_ref_long$Experiment, 
                                          parallel = "multicore")

# which(cd.exp.q1.lr.0.5 > 0.45) # exp.143
# live_ref_long[live_ref_long$Experiment == 143,] # Kandlikar et al

# plot(cd.exp.q1.lr.0.5, type="o", pch=19, xlab="Experiment", ylab="Cook's Distance")
# abline(0.45, 0, col = "red")

# re-do analysis without experiment 143
live_ref_no_143 <- live_ref_long %>% filter(Experiment != 143)
q1.lr.0.5.no143 <- run_metamod(which_df = live_ref_no_143, 
                               which_response_column = "absolute_FDSD_effects",
                               moderators = " ~ (set-1)",
                               impute_cov_r = 0.5)

cd.q1.lr.0.5.no143 <- cooks.distance.rma.mv(q1.lr.0.5.no143, 
                                            cluster = live_ref_no_143$Experiment, 
                                            parallel = "multicore")
# which(cd.q1.lr.0.5.no143 > 0.45) # detect another influential experiment: exp.138
# live_ref_long[live_ref_long$Experiment == 138,]

# delete this other influential exp.:
live_ref_no_143_138 <- live_ref_long %>% filter(Experiment != 138 & Experiment != 143)
q1.lr.0.5.no143.138 <- run_metamod(which_df = live_ref_no_143_138, 
                                   which_response_column = "absolute_FDSD_effects",
                                   moderators = " ~ (set-1)",
                                   impute_cov_r = 0.5)
cd.q1.lr.0.5.no143.138 <- cooks.distance.rma.mv(q1.lr.0.5.no143.138, 
                                                cluster = live_ref_no_143_138$Experiment, 
                                                parallel = "multicore")
# which(cd.q1.lr.0.5.no143.138 > 0.45) # now, no more influential studies

coef_test(q1.lr.0.5.no143.138, vcov = "CR2") # check if different from zero
```

```
##   Coef. Estimate    SE t-stat d.f. p-val (Satt) Sig.
## 1 setFD    0.599 0.114   5.24 12.8       <0.001  ***
## 2 setSD    0.433 0.075   5.77 11.6       <0.001  ***
```

```
conf_int(q1.lr.0.5.no143.138, vcov = "CR2") # take look at the confidence interval
```

```
##    Coef Estimate    SE d.f. Lower 95% CI Upper 95% CI
## 1 setFD    0.599 0.114 12.8        0.352        0.847
## 2 setSD    0.433 0.075 11.6        0.269        0.597
```

```
q1.lr.trim.wald <- Wald_test(q1.lr.0.5.no143.138, constraints = constrain_equal(1:2), vcov = "CR2")

# find "influential points"
# the code below was run and saved, here just being sourced to save time
# cd.points.q1.lr.0.5 <- cooks.distance.rma.mv(q1.lr.0.5, parallel = "multicore")
# write.csv(cd.points.q1.lr.0.5, "cd.points.q1.lr.0.5.csv")
cd.points.q1.lr.0.5 <- read.csv("cd.points.q1.lr.0.5.csv", row.names = 1)
# which(cd.points.q1.lr.0.5 > 0.45) # no "influential points"
```

###### Sterile reference influence analysis

In the influence analysis for live soil references we found one study that was influential; we replicated all analyses without these studies (expand code chunk below for details).

```
# find "influential experiments"
# the code code was run and saved, here just being sourced to save time
# cd.exp.q1.xs.0.5<-cooks.distance.rma.mv(q1.xs.0.5, 
#                       cluster = sterile_ref_long$Experiment, #whereas influential studies = 51, 172
#                       reestimate = F,
#                       parallel = "multicore")
# write.csv(cd.exp.q1.xs.0.5, "cd.exp.q1.xs.0.5.csv")
cd.exp.q1.xs.0.5 <- read.csv("cd.exp.q1.xs.0.5.csv")
colnames(cd.exp.q1.xs.0.5) <- c("expt", "cooks_dist")

# cd.exp.q1.xs.0.5[cd.exp.q1.xs.0.5$cooks_dist>0.45, ] # influential experiment = 33; Hendricks et al.
#sterile_ref_long[sterile_ref_long$Experiment ==33,] %>% view()

# plot(cd.exp.q1.xs.0.5, type="o", pch=19, xlab="Experiment", ylab="Cook's Distance")
# abline(0.45, 0, col = "red")

# remove influential experiment 33
sterile_ref_no_33 <- sterile_ref_long %>% filter(Experiment != 33)
q1.xs.0.5.no33 <- run_metamod(which_df = sterile_ref_no_33, 
                              which_response_column = "absolute_FDSD_effects",
                              moderators = " ~ (set-1)",
                              impute_cov_r = 0.5)
# check if there are new influence experiments:
# cooks.distance.rma.mv(q1.xs.0.5.no33, 
#                       cluster = sterile_ref_trim$Experiment, 
#                       reestimate = F,
#                       parallel = "multicore") 
# no other influence experiments

coef_test(q1.xs.0.5.no33, vcov = "CR2") # check if different from zero
```

```
##   Coef. Estimate     SE t-stat d.f. p-val (Satt) Sig.
## 1 setFD    0.760 0.1005   7.56 51.4       <0.001  ***
## 2 setSD    0.188 0.0193   9.73 49.5       <0.001  ***
```

```
conf_int(q1.xs.0.5.no33, vcov = "CR2") # take look at the confidence interval
```

```
##    Coef Estimate     SE d.f. Lower 95% CI Upper 95% CI
## 1 setFD    0.760 0.1005 51.4        0.558        0.961
## 2 setSD    0.188 0.0193 49.5        0.149        0.227
```

```
q1.xs.trim.wald <- Wald_test(q1.xs.0.5.no33, constraints = constrain_equal(1:2), vcov = "CR2")


# find "influential points"
# cd.points.q1.xs.0.5<-cooks.distance.rma.mv(q1.xs.0.5, parallel = "multicore",
#                                            reestimate = F)
# write.csv(cd.points.q1.xs.0.5, "cd.points.q1.xs.0.5.csv")
cd.points.q1.xs.0.5 <- read.csv("cd.points.q1.xs.0.5.csv")
colnames(cd.points.q1.xs.0.5) <- c("expt", "cooks_dist")
# cd.points.q1.xs.0.5[cd.points.q1.xs.0.5$cooks_dist>0.45, ] # no influential points
```

###### Make plots with & without influential points/expts

```
# make dataframe to plot results after deletion
q1.lr.deletion.plot <- bind_rows(conf_int(q1.lr.0.5, vcov = "CR2"),
                              #conf_int(q1.lr.0.5.no143, vcov = "CR2"),
                              conf_int(q1.lr.0.5.no143.138, vcov = "CR2")) %>%
  data.frame() %>% 
  rownames_to_column("which")


q1.lr.deletion.plot <-
  q1.lr.deletion.plot %>%
  mutate(set = c("FD", "SD", "FD", "SD"),
         set = ifelse(set == "FD", "Fitness difference", "(De)stabilization"))

q1.xs.deletion.plot <- bind_rows(conf_int(q1.xs.0.5, vcov = "CR2"),
                              conf_int(q1.xs.0.5.no33, vcov = "CR2")) %>%
  data.frame() %>% 
  rownames_to_column("which")

q1.xs.deletion.plot <-
  q1.xs.deletion.plot %>%
  mutate(set = c("FD", "SD", "FD", "SD"),
         set = ifelse(set == "FD", "Fitness difference", "(De)stabilization"))


q1.deletion.plot <- bind_rows(q1.lr.deletion.plot, q1.xs.deletion.plot) 
q1.deletion.plot <- q1.deletion.plot %>% 
  mutate(dataset = rep(c("Full", "Full", "Influential\n experiment\n removed", "Influential\n experiment\n removed"), 2))%>%
  mutate(yval= rev(c(1, 1.7, 1.3, 2,
                     2.5, 3.2, 2.8, 3.5)))

q1_deletion_plot <- ggplot(q1.deletion.plot, aes(y = beta, x = yval)) +
  geom_ribbon(aes(x = seq(0.75, 2.25, length.out = 8), ymin = -Inf, ymax = Inf), fill = "grey90") + 
  geom_point(size = 5, aes(color = set, shape = dataset)) +
  geom_errorbar(aes(ymin = CI_L, ymax = CI_U), width = 0, size = .4) +
  geom_vline(xintercept = 2.25, size = 0.25) +
  scale_color_manual(name = "", values = c("#CC79A7", "#009E73")) + 
  scale_x_continuous(breaks = c(1.5, 2.75), 
                     expand = c(0,0), limits = c(0.75, 3.9), 
                     labels = c("Sterile reference",
                                "Live reference")) + 
  xlab("") +
  ylab("Effect size") +
  theme_plots() + 
  theme(axis.text.y = element_text(size = 11, color = "black"),
        legend.position = "right",
        plot.title = element_text(face = "bold"),
        plot.title.position = "plot") + 
    ggpubr::geom_bracket(xmin = c(1, 1.4, 2.5, 2.8), 
                         xmax = c(1.6, 2, 3.1, 3.4), 
                         y.position = c(1.1, 1.05, 1.1, 1.05),
                         label = c("***", "***", "*", "**"), tip.length = c(0.01, 0.01),
                         vjust = 0.1) +
  coord_flip() + 
  labs(title = "Influence analysis")

q1_deletion_plot
```

```
# ggsave("figures/q1_deletion_plot.png", q1_deletion_plot, bg = "transparent", height = 5, width = 8)
```

##### Moderator analyses

Now, explore how experimental moderators affect the outcomes

###### Live ref soils Moderator analysis

**Inoculation fraction: High/Low**

```
# Make columns to help conduct the analyses
live_ref_long <-
  live_ref_long %>%
  mutate(inoc_frac_binary = ifelse(`Fraction inoculum` < 0.2501, "low", "high"),
         set_inoc_frac = paste0(set, "_", inoc_frac_binary),
         set_trainingEnv = paste0(set, "_", `Training environment`),
         set_testingEnv = paste0(set, "_", `Testing environment`),
         set_testingCommunity = paste0(set, "_", `Testing community`))

# # Check which comparisons are possible to make in this dataset:
# table(live_ref$`Training environment`) # 14 pairs in field, 58 in lab
# table(live_ref$`Testing environment`) # 3 pairs in the field, 69 in lab
# table(live_ref$`Testing community`) # 2 pairs population; 15 pairs community, 55 pairs individual
# table(live_ref_long$inoc_frac_binary)/2 # 17 pairs with low, 55 pairs with high
# 
# nrow(unique(live_ref_long[which(live_ref_long$`Training environment` == "Field"),"Experiment"]))
# nrow(unique(live_ref_long[which(live_ref_long$`Training environment` == "Lab"),"Experiment"]))
# nrow(unique(live_ref_long[which(live_ref_long$inoc_frac_binary == "low"),"Experiment"]))
# nrow(unique(live_ref_long[which(live_ref_long$inoc_frac_binary == "high"),"Experiment"]))
# nrow(unique(live_ref_long[which(live_ref_long$`Testing community` == "Population"),"Experiment"]))
# nrow(unique(live_ref_long[which(live_ref_long$`Testing community` == "Community"),"Experiment"]))
# nrow(unique(live_ref_long[which(live_ref_long$`Testing community` == "Individual"),"Experiment"]))

q1.lr.inocFrac <- run_metamod(which_df = live_ref_long, 
                              which_response_column = "absolute_FDSD_effects",
                              moderators = " ~ (set_inoc_frac-1)",
                              impute_cov_r = 0.5)
# coef_test(q1.lr.inocFrac, vcov = "CR2") %>% # check if different from zero
#   kable() %>%
#   kable_styling(full_width = F)
conf_int(q1.lr.inocFrac, vcov = "CR2") %>% # take look at the confidence interval
  kable() %>%
  kable_styling(full_width = F)
```

|  | beta | SE | df | CI\_L | CI\_U |
| --- | --- | --- | --- | --- | --- |
| set\_inoc\_fracFD\_high | 0.6447761 | 0.1100202 | 12.054273 | 0.4051824 | 0.8843698 |
| set\_inoc\_fracFD\_low | 0.4829630 | 0.0590816 | 1.678832 | 0.1759622 | 0.7899639 |
| set\_inoc\_fracSD\_high | 0.4054537 | 0.0807271 | 11.602875 | 0.2288946 | 0.5820129 |
| set\_inoc\_fracSD\_low | 0.1870284 | 0.0359641 | 1.680444 | 0.0003673 | 0.3736896 |

```
Wald_test(q1.lr.inocFrac, constraints = constrain_equal(c(1,3)), vcov = "CR2") # Check if FD diff from SD in High inoc
```

```
##  test Fstat df_num df_denom   p_val sig
##   HTZ  10.5      1     9.19 0.00984  **
```

```
Wald_test(q1.lr.inocFrac, constraints = constrain_equal(c(2,4)), vcov = "CR2") # Check if FD diff from SD in low inoc
```

```
##  test Fstat df_num df_denom p_val sig
##   HTZ  12.7      1     1.26 0.132
```

---

**Conditioning phase environment: Greenhouse/Field**

```
q1.lr.trainEnv <- run_metamod(which_df = live_ref_long, 
                              which_response_column = "absolute_FDSD_effects",
                              moderators = " ~ (set_trainingEnv-1)",
                              impute_cov_r = 0.5)

# coef_test(q1.lr.trainEnv, vcov = "CR2") %>% # check if different from zero
#   kable() %>%
#   kable_styling(full_width = F)
conf_int(q1.lr.trainEnv, vcov = "CR2") %>% # take look at the confidence interval
  kable() %>%
  kable_styling(full_width = F)
```

|  | beta | SE | df | CI\_L | CI\_U |
| --- | --- | --- | --- | --- | --- |
| set\_trainingEnvFD\_Field | 0.3760611 | 0.1401516 | 3.579264 | -0.0317825 | 0.7839048 |
| set\_trainingEnvFD\_Lab | 0.7106848 | 0.1060617 | 9.490630 | 0.4726340 | 0.9487356 |
| set\_trainingEnvSD\_Field | 0.3952396 | 0.0599903 | 3.601112 | 0.2211453 | 0.5693339 |
| set\_trainingEnvSD\_Lab | 0.3714087 | 0.0951298 | 9.729855 | 0.1586459 | 0.5841715 |

```
Wald_test(q1.lr.trainEnv, constraints = constrain_equal(c(1,3)), vcov = "CR2") # Field: FD vs SD
```

```
##  test  Fstat df_num df_denom p_val sig
##   HTZ 0.0171      1     2.95 0.904
```

```
Wald_test(q1.lr.trainEnv, constraints = constrain_equal(c(2,4)), vcov = "CR2") # Lab: FD vs SD
```

```
##  test Fstat df_num df_denom  p_val sig
##   HTZ  57.6      1     5.45 <0.001 ***
```

---

**Response phase growth type: Individual/Population/Community**

```
# Testing community ----
q1.lr.testComm <- run_metamod(which_df = live_ref_long, 
                              which_response_column = "absolute_FDSD_effects",
                              moderators = " ~ (set_testingCommunity-1)",
                              impute_cov_r = 0.5)
# coef_test(q1.lr.testComm, vcov = "CR2") %>% # check if different from zero
  # kable() %>%
  # kable_styling(full_width = F)
conf_int(q1.lr.testComm, vcov = "CR2") %>% # take look at the confidence interval
  kable() %>%
  kable_styling(full_width = F)
```

|  | beta | SE | df | CI\_L | CI\_U |
| --- | --- | --- | --- | --- | --- |
| set\_testingCommunityFD\_Community | 0.5418709 | 0.2002920 | 1.970319 | -0.3324770 | 1.4162188 |
| set\_testingCommunityFD\_Individual | 0.6726773 | 0.1123367 | 10.190987 | 0.4230098 | 0.9223448 |
| set\_testingCommunityFD\_Population | 0.3552481 | 0.0735266 | 1.000013 | -0.5789660 | 1.2894621 |
| set\_testingCommunitySD\_Community | 0.2430481 | 0.1031686 | 1.940225 | -0.2142171 | 0.7003133 |
| set\_testingCommunitySD\_Individual | 0.4116234 | 0.0895896 | 9.882843 | 0.2116842 | 0.6115626 |
| set\_testingCommunitySD\_Population | 0.1959575 | 0.1218909 | 1.000043 | -1.3526544 | 1.7445695 |

```
Wald_test(q1.lr.testComm, constraints = constrain_equal(c(1,4)), vcov = "CR2") # Q1 FD v SD, community
```

```
##  test Fstat df_num df_denom p_val sig
##   HTZ  1.31      1     1.86 0.379
```

```
Wald_test(q1.lr.testComm, constraints = constrain_equal(c(2,5)), vcov = "CR2") # Q1 FD v SD, individual
```

```
##  test Fstat df_num df_denom   p_val sig
##   HTZ  20.4      1     5.94 0.00412  **
```

```
Wald_test(q1.lr.testComm, constraints = constrain_equal(c(3,6)), vcov = "CR2") # Q1 FD v SD, population
```

```
##  test Fstat df_num df_denom p_val sig
##   HTZ  10.6      1     1.01 0.189
```

***Now, put it all together to make the plot.***

```
q1.lr.coefs.plot <- bind_rows(conf_int(q1.lr.0.5, vcov = "CR2"),
                              conf_int(q1.lr.trainEnv, vcov = "CR2"),
                              conf_int(q1.lr.inocFrac, vcov =  "CR2"),
                              conf_int(q1.lr.testComm, vcov = "CR2")) %>%
  data.frame() %>% 
  rownames_to_column("which")
q1.lr.coefs.plot <-
  q1.lr.coefs.plot %>%
  mutate(yval = rev(c(0.9,1.4,1.9, 1.1,1.6,2.1, 
                      2.4,2.9,     2.6,3.1, 
                      3.4,3.9,     3.6,4.1, 
                      4.4,4.6)),
         set = c("FD", "SD", "FD", "FD", "SD", "SD","FD", "FD", "SD", "SD",
                 "FD", "FD", "FD", "SD", "SD", "SD"),
         set = ifelse(set == "FD", "Fitness difference", "(De)stabilization"))


q1.mods.lr.plot <-
  ggplot(q1.lr.coefs.plot, aes(y = beta, x = yval)) +
  geom_ribbon(aes(x = seq(0.75, 2.25, length.out = 16), ymin = -Inf, ymax = Inf), fill = "grey90") + 
  geom_ribbon(aes(x = seq(3.25, 4.25, length.out = 16), ymin = -Inf, ymax = Inf), fill = "grey90") + 

  geom_point(size = 4, aes(fill = set), shape = 21) +
  geom_errorbar(aes(ymin = CI_L, ymax = CI_U), width = 0, size = .4) +
  geom_vline(xintercept = c(2.25, 3.25, 4.25), size = 0.25) + 
  geom_vline(xintercept = c(1.25, 1.75, 2.75, 3.75), size = 0.25, linetype = 2) + 
  scale_fill_manual(name = "", values = c("#CC79A7", "#009E73")) + 
  scale_x_continuous(breaks = c(1.5, 2.75, 3.75, 4.5), 
                     expand = c(0,0), limits = c(0.75, 4.75), 

                     labels = c("Response\nphase\ngrowth\ntype",
                                "Inoculation\nFraction",
                                "Training\nphase\nlocation",
                                "Overall")) + 
  annotate("text", y = -Inf, x = c(1.25, 1.75, 2.25, 2.75, 3.25, 3.75, 4.25, Inf),
           label = c(
             ' Community (15 pairs / 3 experiments)',
             ' Individual (55 pairs / 13 experiments)',
             ' Population (2 pairs / 2 experiments)',
             ' Low (<25%) (17 pairs / 3 experiments)',
             ' High (>25%) (55 pairs / 15 experiments)',
             ' Greenhouse (58 pairs / 15 experiments)', 
             ' Field (14 pairs / 3 experiments)', 
             ' All pairs (72 pairs / 18 experiments)'), 
           hjust = 0, vjust = 1.5,
           color = "grey25", fontface = "italic", size = 3.5) +
  xlab("") +
  ylab("Effect size") +
  theme_plots() + 
  theme(axis.text.y = element_text(size = 11, color = "black"),
        legend.position = "bottom",
        plot.title = element_text(face = "bold"),
        plot.title.position = "plot") + 
    ggpubr::geom_bracket(xmin = c(1.4, 2.9, 3.4, 4.4), 
                         xmax = c(1.6, 3.1, 3.6, 4.6), 
                         y.position = 1.25,
                         label = c("**", "***","**", "**"), tip.length = c(0.01, 0.01),
                         vjust = 0.5) +
  coord_flip() + 
  labs(title = "(A) Live soil reference moderator analysis")

q1.mods.lr.plot
```

###### Sterile soil moderator analysis

```
# Inoculation fraction, low/high
sterile_ref_long <-
  sterile_ref_long %>%
  mutate(`Training environment` = ifelse(`Training environment` == "LabField", "Lab", `Training environment`)) %>%
  mutate(inoc_frac_binary = ifelse(`Fraction inoculum` < 0.2501, "low", "high"),
         set_inoc_frac = paste0(set, "_", inoc_frac_binary),
         set_trainingEnv = paste0(set, "_", `Training environment`),
         set_testingEnv = paste0(set, "_", `Testing environment`),
         set_controlType = paste0(set, "_", control_type),
         Study_sterilization_method = ifelse(Study_sterilization_method %in% 
                                               c("not specified", "pasteurization", "steam-pasteurization", "steaming"), 
                                             "other", 
                                             Study_sterilization_method),
         set_sterileMethod = paste0(set, "_", Study_sterilization_method),
         set_communityType = paste0(set, "_", `Testing community`))

# Check which comparisons are possible to make in this dataset:
# table(sterile_ref$`Training environment`) # 177 pairs in field, 235 in lab, 34 in LabField
# table(sterile_ref$`Testing environment`) # 2 pairs in the field, 444 in lab
# table(sterile_ref_long$inoc_frac_binary)/2 # 117 pairs with low, 329 pairs with high
# table(sterile_ref$control_type)
```

**Inoculation fraction: High/Low**

```
q1.xs.inocFrac <- run_metamod(which_df = sterile_ref_long, 
                              which_response_column = "absolute_FDSD_effects",
                              moderators = " ~ (set_inoc_frac-1)",
                              impute_cov_r = 0.5)
# coef_test(q1.xs.inocFrac, vcov = "CR2") # check if different from zero
conf_int(q1.xs.inocFrac, vcov = "CR2") %>% # take look at the confidence interval
  kable() %>%
  kable_styling(full_width = F)
```

|  | beta | SE | df | CI\_L | CI\_U |
| --- | --- | --- | --- | --- | --- |
| set\_inoc\_fracFD\_high | 0.7434680 | 0.1284906 | 17.83454 | 0.4733396 | 1.0135965 |
| set\_inoc\_fracFD\_low | 0.7476425 | 0.1362295 | 33.69125 | 0.4706974 | 1.0245875 |
| set\_inoc\_fracSD\_high | 0.2670052 | 0.0651052 | 18.25240 | 0.1303597 | 0.4036507 |
| set\_inoc\_fracSD\_low | 0.1857668 | 0.0172844 | 34.95795 | 0.1506761 | 0.2208574 |

```
Wald_test(q1.xs.inocFrac, constraints = constrain_equal(c(1,3)), vcov = "CR2") #high: FD vs SD
```

```
##  test Fstat df_num df_denom   p_val sig
##   HTZ  13.9      1     17.3 0.00166  **
```

```
Wald_test(q1.xs.inocFrac, constraints = constrain_equal(c(2,4)), vcov = "CR2") # Low: FD vs SD
```

```
##  test Fstat df_num df_denom  p_val sig
##   HTZ  20.4      1     32.5 <0.001 ***
```

---

**Conditioning phase environment: Greenhouse/Field**

```
# Training environment -------
q1.xs.trainEnv <- run_metamod(which_df = sterile_ref_long, 
                              which_response_column = "absolute_FDSD_effects",
                              moderators = " ~ (set_trainingEnv-1)",
                              impute_cov_r = 0.5)

# coef_test(q1.xs.trainEnv, vcov = "CR2") # check if different from zero
conf_int(q1.xs.trainEnv, vcov = "CR2") %>% # take look at the confidence interval
  kable() %>%
  kable_styling(full_width = F)
```

|  | beta | SE | df | CI\_L | CI\_U |
| --- | --- | --- | --- | --- | --- |
| set\_trainingEnvFD\_Field | 0.6407833 | 0.0891627 | 34.36207 | 0.4596532 | 0.8219135 |
| set\_trainingEnvFD\_Lab | 0.9114088 | 0.2018883 | 17.60269 | 0.4865700 | 1.3362476 |
| set\_trainingEnvSD\_Field | 0.1967454 | 0.0280773 | 36.00329 | 0.1398021 | 0.2536887 |
| set\_trainingEnvSD\_Lab | 0.2384962 | 0.0444441 | 18.75195 | 0.1453902 | 0.3316022 |

```
Wald_test(q1.xs.trainEnv, constraints = constrain_equal(1:3), vcov = "CR2") # Field: FD vs SD
```

```
##  test Fstat df_num df_denom  p_val sig
##   HTZ  24.6      2       35 <0.001 ***
```

```
Wald_test(q1.xs.trainEnv, constraints = constrain_equal(2:4), vcov = "CR2") # Lab: FD vs SD
```

```
##  test Fstat df_num df_denom   p_val sig
##   HTZ  5.95      2     25.4 0.00757  **
```

---

**Response phase reference soil type environment: CmS/CS/FS/GS**

- Recall that CmS = mixed conditioned soils; CS = sterilized conditioned soil; FS = sterilized field soil; GS = sterilized greenhouse soil

```
# Control type ---------
q1.xs.controlType <- run_metamod(which_df = sterile_ref_long, 
                                 which_response_column = "absolute_FDSD_effects",
                                 moderators = " ~ (set_controlType-1)",
                                 impute_cov_r = 0.5)

# coef_test(q1.xs.controlType, vcov = "CR2") # check if different from zero
conf_int(q1.xs.controlType, vcov = "CR2") %>% # take look at the confidence interval
  kable() %>%
  kable_styling(full_width = F)
```

|  | beta | SE | df | CI\_L | CI\_U |
| --- | --- | --- | --- | --- | --- |
| set\_controlTypeFD\_CmS | 0.2575516 | 0.0552553 | 1.944070 | 0.0131307 | 0.5019724 |
| set\_controlTypeFD\_CS | 0.6006857 | 0.1069559 | 25.887008 | 0.3807880 | 0.8205834 |
| set\_controlTypeFD\_FS | 0.7496226 | 0.1725701 | 6.562040 | 0.3359728 | 1.1632725 |
| set\_controlTypeFD\_GS | 1.4579523 | 0.1707753 | 9.477152 | 1.0745765 | 1.8413281 |
| set\_controlTypeSD\_CmS | 0.4365872 | 0.2706576 | 1.991339 | -0.7328256 | 1.6060001 |
| set\_controlTypeSD\_CS | 0.1703291 | 0.0270217 | 32.015169 | 0.1152887 | 0.2253696 |
| set\_controlTypeSD\_FS | 0.3016669 | 0.0470920 | 6.175626 | 0.1872267 | 0.4161070 |
| set\_controlTypeSD\_GS | 0.2222953 | 0.0303956 | 11.619813 | 0.1558279 | 0.2887628 |

```
Wald_test(q1.xs.controlType, constraints = constrain_equal(c(1,5)), vcov = "CR2") # CmS: Fd vs SD
```

```
##  test Fstat df_num df_denom p_val sig
##   HTZ 0.422      1     1.92 0.585
```

```
Wald_test(q1.xs.controlType, constraints = constrain_equal(c(2,6)), vcov = "CR2") # CS: FD vs SD
```

```
##  test Fstat df_num df_denom  p_val sig
##   HTZ  22.2      1       23 <0.001 ***
```

```
Wald_test(q1.xs.controlType, constraints = constrain_equal(c(3,7)), vcov = "CR2") # FS: FD vs SD
```

```
##  test Fstat df_num df_denom  p_val sig
##   HTZ  5.37      1     6.16 0.0586   .
```

```
Wald_test(q1.xs.controlType, constraints = constrain_equal(c(4,8)), vcov = "CR2") # GS: FD vs SD
```

```
##  test Fstat df_num df_denom  p_val sig
##   HTZ  72.8      1     8.26 <0.001 ***
```

---

**Sterilization method: Autoclaving, Gamma Radiation, Heating, or Other**

```
# Sterilization method ---------
q1.xs.sterileMethod <- run_metamod(which_df = sterile_ref_long, 
                              which_response_column = "absolute_FDSD_effects",
                              moderators = " ~ (set_sterileMethod-1)",
                              impute_cov_r = 0.5)
# coef_test(q1.xs.sterileMethod, vcov = "CR2") # check if different from zero
conf_int(q1.xs.sterileMethod, vcov = "CR2") %>% # take look at the confidence interval
  kable() %>%
  kable_styling(full_width = F)
```

|  | beta | SE | df | CI\_L | CI\_U |
| --- | --- | --- | --- | --- | --- |
| set\_sterileMethodFD\_autoclaving | 0.4323789 | 0.0994516 | 20.065028 | 0.2249697 | 0.6397882 |
| set\_sterileMethodFD\_gamma irradiation | 1.0147852 | 0.1665735 | 7.636759 | 0.6274625 | 1.4021080 |
| set\_sterileMethodFD\_heating | 1.9391413 | 0.0205284 | 3.994673 | 1.8821153 | 1.9961672 |
| set\_sterileMethodFD\_other | 0.3852328 | 0.1140017 | 8.180240 | 0.1233493 | 0.6471164 |
| set\_sterileMethodSD\_autoclaving | 0.1499915 | 0.0165809 | 28.988289 | 0.1160792 | 0.1839039 |
| set\_sterileMethodSD\_gamma irradiation | 0.3179508 | 0.0856622 | 10.503562 | 0.1283166 | 0.5075851 |
| set\_sterileMethodSD\_heating | 0.3166535 | 0.0100848 | 3.997874 | 0.2886478 | 0.3446591 |
| set\_sterileMethodSD\_other | 0.1959404 | 0.0498763 | 8.629237 | 0.0823690 | 0.3095119 |

```
Wald_test(q1.xs.sterileMethod, constraints = constrain_equal(c(1,5)), vcov = "CR2") # Autoclaving: FD vs SD
```

```
##  test Fstat df_num df_denom   p_val sig
##   HTZ  9.09      1     19.6 0.00697  **
```

```
Wald_test(q1.xs.sterileMethod, constraints = constrain_equal(c(2,6)), vcov = "CR2") # Gamma irradiation: FD vs SD
```

```
##  test Fstat df_num df_denom   p_val sig
##   HTZ  19.2      1      7.4 0.00282  **
```

```
Wald_test(q1.xs.sterileMethod, constraints = constrain_equal(c(3,7)), vcov = "CR2") # Heating: FD vs SD
```

```
##  test Fstat df_num df_denom  p_val sig
##   HTZ  8084      1     3.99 <0.001 ***
```

```
Wald_test(q1.xs.sterileMethod, constraints = constrain_equal(c(4,8)), vcov = "CR2") # Other: FD vs SD
```

```
##  test Fstat df_num df_denom  p_val sig
##   HTZ  3.98      1     8.12 0.0806   .
```

---

**Response phase growth type: Individual/Population/Community**

```
# Community type ---------
q1.xs.communityType <- run_metamod(which_df = sterile_ref_long, 
                                   which_response_column = "absolute_FDSD_effects",
                                   moderators = " ~ (set_communityType-1)",
                                   impute_cov_r = 0.5)
# coef_test(q1.xs.communityType, vcov = "CR2") # check if different from zero
conf_int(q1.xs.communityType, vcov = "CR2") %>% # take look at the confidence interval
  kable() %>%
  kable_styling(full_width = F)
```

|  | beta | SE | df | CI\_L | CI\_U |
| --- | --- | --- | --- | --- | --- |
| set\_communityTypeFD\_Community | 1.8341309 | 0.0615156 | 6.089070 | 1.6841398 | 1.9841220 |
| set\_communityTypeFD\_Individual | 0.5261654 | 0.0876983 | 24.511765 | 0.3453647 | 0.7069661 |
| set\_communityTypeFD\_Population | 0.7921600 | 0.3486605 | 2.859651 | -0.3488865 | 1.9332065 |
| set\_communityTypeSD\_Community | 0.3859826 | 0.0476435 | 8.219688 | 0.2766254 | 0.4953397 |
| set\_communityTypeSD\_Individual | 0.1834898 | 0.0299204 | 37.119545 | 0.1228719 | 0.2441077 |
| set\_communityTypeSD\_Population | 0.1523392 | 0.0214212 | 5.900417 | 0.0997082 | 0.2049702 |

```
Wald_test(q1.xs.communityType, constraints = constrain_equal(1:2), vcov = "CR2")
```

```
##  test Fstat df_num df_denom  p_val sig
##   HTZ   149      1     13.8 <0.001 ***
```

```
Wald_test(q1.xs.communityType, constraints = constrain_equal(3:4), vcov = "CR2")
```

```
##  test Fstat df_num df_denom p_val sig
##   HTZ  1.33      1     3.22 0.327
```

```
Wald_test(q1.xs.communityType, constraints = constrain_equal(c(1,4)), vcov = "CR2")
```

```
##  test Fstat df_num df_denom  p_val sig
##   HTZ   300      1     6.48 <0.001 ***
```

```
Wald_test(q1.xs.communityType, constraints = constrain_equal(c(2,5)), vcov = "CR2")
```

```
##  test Fstat df_num df_denom  p_val sig
##   HTZ  15.7      1     26.1 <0.001 ***
```

```
Wald_test(q1.xs.communityType, constraints = constrain_equal(c(3,6)), vcov = "CR2")
```

```
##  test Fstat df_num df_denom p_val sig
##   HTZ  3.61      1     3.13  0.15
```

```
Wald_test(q1.xs.0.5, constraints = constrain_equal(c(1,2)), vcov = "CR2")
```

```
##  test Fstat df_num df_denom  p_val sig
##   HTZ  33.1      1     50.7 <0.001 ***
```

**Now, put it all together into a plot**

```
q1.xs.coefs.plot <- bind_rows(conf_int(q1.xs.0.5, vcov = "CR2"),
                              conf_int(q1.xs.trainEnv, vcov = "CR2"),
                              conf_int(q1.xs.sterileMethod, vcov = "CR2"),
                              conf_int(q1.xs.inocFrac, vcov = "CR2"),
                              conf_int(q1.xs.controlType, vcov = "CR2"),
                              conf_int(q1.xs.communityType, vcov = "CR2")) %>%
  data.frame() %>% 
  rownames_to_column("which")
q1.xs.coefs.plot <-
  q1.xs.coefs.plot %>%
  mutate(yval = rev(c(c(0.9,1.4,1.9)+0.05, c(1.1,1.6,2.1)-0.05, # Community type
                      c(2.4,2.9,3.4,3.9)+0.05, c(2.6,3.1,3.6,4.1)-0.05, # Control type
                      c(4.4,4.9)+0.05, c(4.6,5.1)-0.05, # Inoculation high/low
                      c(5.4,5.9,6.4,6.9)+0.05, c(5.6,6.1,6.6,7.1)-0.05, # Sterile method
                      c(7.4,7.9)+0.05, c(7.6,8.1)-0.05, # Training Environment
                      8.4+0.05,8.6-0.05) # Overall
                    ),
         set = ifelse(stringr::str_detect(which, "FD"), "FD", "SD"),
         set = ifelse(set == "FD", "Fitness difference", "(De)stabilization"))


q1.mods.xs.plot <-
  ggplot(q1.xs.coefs.plot, aes(y = beta, x = yval)) +
  coord_flip() +
  geom_ribbon(aes(x = seq(0.75, 2.25, length.out = 32), ymin = -Inf, ymax = Inf), fill = "grey90") + 
  geom_ribbon(aes(x = seq(4.25, 5.25, length.out = 32), ymin = -Inf, ymax = Inf), fill = "grey90") + 
  geom_ribbon(aes(x = seq(7.25, 8.25, length.out = 32), ymin = -Inf, ymax = Inf), fill = "grey90") + 
  geom_point(size = 4, aes(fill = set), shape = 21) +
  geom_errorbar(aes(ymin = CI_L, ymax = CI_U), width = 0, size = .4) +
  geom_vline(xintercept = c(2.25, 4.25, 5.25, 7.25, 8.25), size = 0.25) + 
  geom_vline(xintercept = c(1.25, 1.75, 2.75, 3.25, 3.75, 4.75,  5.75, 6.25, 6.75,7.75), 
             size = 0.25, linetype = 2) + 
  scale_fill_manual(name = "", values = c("#CC79A7", "#009E73")) + 
  scale_x_continuous(breaks = c(1.5, 3.25, 4.75, 6.25, 7.75, 8.5), 
                     expand = c(0,0), limits = c(0.75, 8.75), 
                     labels = c("Response\nphase\ngrowth\ntype", 
                                "Sterile\nsoil\nsource", 
                                "Inoculation\nfraction", 
                                "Soil\nsterilization\nmethod", 
                                "Training\nphase\nlocation",
                                "Overall")) + 
  annotate("text", y = -Inf, x = c(1.25, 1.75, 2.25, 2.75, 
                                   3.25, 3.75, 4.25,
                                   4.75, 5.25, 5.75, 6.25,
                                   6.75,7.25, 7.75,8.25, Inf),
           label = c(
             ' Population (37 pairs / 9 experiments)',
             ' Individual (189 pairs / 47 experiments)',
             ' Community (220 pairs / 11 experiments)',
             ' Greenhouse soil (216 pairs / 15 experiments)',
             ' Field soil (20 pairs / 10 experiments)',
             ' Phase 1 soil (species specific) (195 pairs / 39 experiments)',
             ' Phase 1 soil (mixed species) (15 pairs / 3 experiments)',
             ' Low (<25%) (329 pairs / 44 experiments)',
             ' High (>25%) (117 pairs / 23 experiments)',
             ' Other (17 pairs / 11 experiments)',
             ' Heating (189 pairs / 5 experiments)',
             ' Gamma irradiation (122 pairs / 13 experiments)',
             ' Autoclaving (118 pairs / 38 experiments)',
             ' Greenouse (269 pairs / 22 experiments)',
             ' Field (177 pairs / 45 experiments)',
             ' All pairs (446 pairs / 67 experiments)'
             ), 
           hjust = 0, vjust = 1.5,
           color = "grey25", fontface = "italic", size = 3.5) +
  xlab("") +
  ylab("Effect size") +
  theme_plots() + 
  theme(axis.text.y = element_text(size = 11, color = "black"),
        legend.position = "bottom",
        plot.title = element_text(face = "bold"),
        plot.title.position = "plot") + 
    ggpubr::geom_bracket(xmin = c(8.45, 7.95,7.45, 7.05,6.55,6.05, 4.95,4.45, 3.45,2.45, 1.95,1.45),
                         xmax = c(8.55, 8.05,7.55, 6.95,6.45,5.95, 5.05,4.55, 3.55,2.55, 2.05,1.55),
                         y.position = c(1.5,1.5,1.5,1.5,1.5, 2.1, 1.5,1.5,1.5, 2.1, 2.1, 1.5),
                         tip.length = c(0.01,0.01),
                         vjust = 1,
                         label = c("***", "***", "**", "**", "**", "***", "**", "***", "***", "***", "***", "***")) +
  labs(title = "(B) Sterile soil reference moderator analysis")

q1.mods.xs.plot
```

```
# table(sterile_ref_long$`Training environment`) # 177 pairs in field, 235 in lab, 34 in LabField
# table(sterile_ref$`Testing environment`) # 2 pairs in the field, 444 in lab
# table(sterile_ref_long$inoc_frac_binary)/2 # 117 pairs with low, 329 pairs with high
# table(sterile_ref_long$`Testing community`)/2 # 440 community, 378 individual, 74 population
# table(sterile_ref_long$Study_sterilization_method)/2 # 236 pairs autoclave, 244 pairs gamma irradiation, 378 pairs heating, 34 pairs 'other'
# table(sterile_ref$control_type) # 15 CmS, 195 CS, 20 FS, 216 GS
# nrow(sterile_ref)
# 
# nrow(unique(sterile_ref_long[which(sterile_ref_long$control_type == "CmS"),"Experiment"]))
# nrow(unique(sterile_ref_long[which(sterile_ref_long$control_type == "CS"),"Experiment"]))
# nrow(unique(sterile_ref_long[which(sterile_ref_long$control_type == "GS"),"Experiment"]))
# nrow(unique(sterile_ref_long[which(sterile_ref_long$control_type == "FS"),"Experiment"]))
# 
# nrow(unique(sterile_ref_long[which(sterile_ref_long$`Training environment` == "Field"),"Experiment"]))
# nrow(unique(sterile_ref_long[which(sterile_ref_long$`Training environment` == "Lab"),"Experiment"]))
# nrow(unique(sterile_ref_long[which(sterile_ref_long$inoc_frac_binary == "low"),"Experiment"]))
# nrow(unique(sterile_ref_long[which(sterile_ref_long$inoc_frac_binary == "high"),"Experiment"]))
# 
# nrow(unique(sterile_ref_long[which(sterile_ref_long$`Testing community` == "Population"),"Experiment"]))
# nrow(unique(sterile_ref_long[which(sterile_ref_long$`Testing community` == "Individual"),"Experiment"]))
# nrow(unique(sterile_ref_long[which(sterile_ref_long$`Testing community` == "Community"),"Experiment"]))
# 
# nrow(unique(sterile_ref_long[which(sterile_ref_long$Study_sterilization_method == "heating"),"Experiment"]))
# nrow(unique(sterile_ref_long[which(sterile_ref_long$Study_sterilization_method == "gamma irradiation"),"Experiment"]))
# nrow(unique(sterile_ref_long[which(sterile_ref_long$Study_sterilization_method == "autoclaving"),"Experiment"]))
# nrow(unique(sterile_ref_long[which(sterile_ref_long$Study_sterilization_method == "other"),"Experiment"]))
```

*For the supplement of the main paper, we put these plots together into one multipanel figure (expand following code block for details).*

```
q1_mods_plot <- 
  q1.mods.lr.plot + q1.mods.xs.plot + 
  plot_layout(guides = "collect") &
  theme(legend.position = "bottom")
q1_mods_plot
```

```
if(save_figures) {
  ggsave("figures/q1_mods_plot.png", q1_mods_plot, 
         bg = "transparent", height = 10, width = 10)
}
```

#### Question 2

Across species pairs, what is the net effect of microbes on competitive outcomes? In other words, how frequently does the balance of microbially-mediated stabilization (or destabilization) and fitness differences predict species coexistence, exclusion, or priority effects

```
## Make a base plot that can be modified for each sub question  

base_coex_plot <- 
  ggplot() +
  geom_ribbon(aes(x = seq(0, 5, length.out = 100),
                  ymin = rep(0, length.out = 100),
                  ymax = seq(0, 5, length.out = 100)),
              fill = alpha("#99D594", .5)) +
  geom_ribbon(aes(x = seq(0, -5, length.out = 100),
                  ymin = rep(0, length.out = 100),
                  ymax = seq(0, 5, length.out = 100)),
              fill = alpha("#FFFFBF", 1)) +
  annotate("segment", x = 0, y = 0, xend = 5, yend = 5, linetype = 2) +
  annotate("segment", x = 0, y = 0, xend = -5, yend = 5, linetype = 2) +
  geom_hline(yintercept = 0) +
  geom_vline(xintercept = 0) +
  xlab(bquote(atop("",
                   -frac(1,2)~(m["1A"]-m["1B"]-m["2A"]+m["2B"]) ))) +
  ylab(bquote(atop("Fitness difference",
                   frac(1,2)~(m["1A"]+m["1B"]-m["2A"]-m["2B"])))) +
  annotate("text", x = 0.2, y = -Inf, hjust = 0, vjust = 0, label = "Stabilization", color = "black", size = 4) +
  annotate("text", x = -0.2, y = -Inf, hjust = 1, vjust = 0, label = "Destabilization",  color = "black", size = 4) +
  annotate("segment", x = 0.2, xend = 1.25, y = -.1, yend = -.1, 
           colour = "grey25", size = 0.5, arrow = arrow(length = unit(0.075, "inches"))) + 
  annotate("segment", x = -0.2, xend = -1.25, y = -.1, yend = -.1, 
           colour = "grey25", size = 0.5, arrow = arrow(length = unit(0.075, "inches"))) +
  theme_plots() + 
  theme(axis.line = element_blank())
```

##### Start sampling approach

To do this sampling we use the custom function `simulation_ci`, which we imported in sourcing the function file above.

###### Live soil

```
FL_simulation_out <- simulation_ci(data_with_SDFD = live_ref)
live_ref_sims <- FL_simulation_out$outcomes_by_pair
live_ref_sims_by_run <- FL_simulation_out$outcomes_by_run

# Define a new function to summarize outcomes
percent_outcomes <- function(outcomes_by_run) {
  format_range <- function(range) {
    paste0(range[2], " (", range[1], "-", range[3],")")
  }
  range.exclusion <-  percent(quantile(outcomes_by_run$n_reps_exclusion/rowSums(outcomes_by_run), 
                                       c(0.05, 0.5, 0.95)), accuracy = .1) %>% format_range
  range.coex <- percent(quantile(outcomes_by_run$n_reps_coex/rowSums(outcomes_by_run), 
                                 c(0.05, 0.5, 0.95)), accuracy = .1) %>% format_range
  range.pes <- percent(quantile(outcomes_by_run$n_reps_pes/rowSums(outcomes_by_run), 
                                c(0.05, 0.5, 0.95)), accuracy = .1) %>% format_range
  return(list(exclusion = range.exclusion, coex = range.coex, pes = range.pes))
}

percent_outcomes(live_ref_sims_by_run)
```

```
## $exclusion
## [1] "69.4% (62.5%-76.4%)"
## 
## $coex
## [1] "22.2% (16.7%-29.2%)"
## 
## $pes
## [1] "6.9% (4.2%-11.1%)"
```

```
x_min <- with(live_ref_sims, min(mean_SD))-0.1
x_max <- with(live_ref_sims, max(mean_SD))+0.1
y_min <- with(live_ref_sims, min(abs(mean_FD)))-0.15
y_max <- with(live_ref_sims, max(abs(mean_FD)))+0.15

(scatterpie_FL <- base_coex_plot +
  scale_x_continuous(limits = c(x_min, x_max)) +
  scale_y_continuous(limits = c(y_min, y_max)) +
  annotate("segment", x = 0, y = 0, xend =  min(x_max, y_max),
           yend = min(x_max, y_max), 
           linetype = 2) +
  annotate("segment", x = 0, y = 0, xend =  max(x_min, -y_max),
           yend =  min(-x_min, y_max),
           linetype = 2) +
  geom_scatterpie(aes(x=mean_SD, y=abs(mean_FD), r = 0.07), 
                  data=live_ref_sims, size = 0.15,
                  cols=c("n_reps_exclusion", "n_reps_coex", "n_reps_pes")) +
  scale_fill_manual(values = c( "#D55E00", "#0072B2","white"),
                    labels =  c( "exclusion", "coexistence", "priority effect"),
                    name = "") +
    guides(fill = "none") +
    labs(title = "(A) Species pairs with reference growth in live soil") + 
  coord_fixed() +
  theme(axis.line = element_line(size = 0),
        legend.position = "none",
        legend.background = element_rect(colour = "grey50", size = .3),
        legend.text = element_text(size = 13),
        legend.title = element_text(size = 8, face = "bold"),
        axis.title = element_text(size = 12))
)
```

###### Sterile soil

```
sterile_simulation_out <- simulation_ci(data_with_SDFD = sterile_ref)
sterile_ref_sims <- sterile_simulation_out$outcomes_by_pair
sterile_ref_sims_by_run <- sterile_simulation_out$outcomes_by_run

percent_outcomes(sterile_ref_sims_by_run)
```

```
## $exclusion
## [1] "81.2% (78.7%-83.6%)"
## 
## $coex
## [1] "12.8% (10.8%-14.8%)"
## 
## $pes
## [1] "6.1% (4.5%-7.6%)"
```

```
x_min <- with(sterile_ref_sims, min(mean_SD))-1
x_max <- with(sterile_ref_sims, max(mean_SD))+1
y_min <- with(sterile_ref_sims, min(abs(mean_FD)))-0.15
y_max <- with(sterile_ref_sims, max(abs(mean_FD)))+0.15

(scatterpie_XS_all <- 
    base_coex_plot +
    scale_x_continuous(limits = c(x_min, x_max)) +
    scale_y_continuous(limits = c(y_min, y_max)) +
    annotate("segment", x = 0, y = 0, xend =  min(x_max, y_max),
             yend = min(x_max, y_max), 
             linetype = 2) +
    annotate("segment", x = 0, y = 0, xend = max(x_min, -y_max),
             yend = min(-x_min, y_max),
             linetype = 2) +
    
    geom_scatterpie(aes(x=mean_SD, y=abs(mean_FD), r = 0.12), 
                    data= sterile_ref_sims, size = 0.15,
                    cols=c("n_reps_exclusion", "n_reps_coex", "n_reps_pes")) +
    scale_fill_manual(values = c( "#D55E00", "#0072B2","white"),
                      labels =  c( "exclusion", "coexistence","priority effect"),
                      name = "") +
    coord_fixed() +
    labs(title = "(B) Species pairs with reference growth in sterile soil") + 
    theme(axis.line = element_line(size = 0),
          legend.justification =c(1,0), 
          legend.position = "bottom",
          # legend.position = c(0.3,0.1),
          legend.text = element_text(size = 12.5),
          legend.background = element_rect(colour = "grey50"),
          legend.title = element_text(size = 8, face = "bold"))
)
```

*For the main text, we assembled these panels together into one figure, see code below for details.*

```
# Combine the two plots together for Figure 4 of the paper
scatterpies_combined <- 
  scatterpie_FL + scatterpie_XS_all + 
  plot_layout(guides = "collect", widths = c(1,1)) & 
  theme(legend.position = "bottom")
if(save_figures) {
  ggsave("figures/scatterpies_combined.png", 
         plot = scatterpies_combined, height = 5.5, width = 11)
}
```

Now we can do a sort of “Moderator analysis” for this sampling approach.

```
# "Moderator analysis"

FL_simulation_out_FILow <- simulation_ci(data_with_SDFD = live_ref %>% filter(`Fraction inoculum` < 0.251))
# percent_outcomes(FL_simulation_out_FILow$outcomes_by_run)

FL_simulation_out_FIHigh <- simulation_ci(data_with_SDFD = live_ref %>% filter(`Fraction inoculum` > 0.25))
# percent_outcomes(FL_simulation_out_FIHigh$outcomes_by_run)

FL_simulation_out_TrainGreenhouse <- simulation_ci(data_with_SDFD = live_ref %>% filter(`Training environment` == "Lab"))
# percent_outcomes(FL_simulation_out_TrainGreenhouse$outcomes_by_run)

FL_simulation_out_TrainField <- simulation_ci(data_with_SDFD = live_ref %>% filter(`Training environment` == "Field"))
# percent_outcomes(FL_simulation_out_TrainField$outcomes_by_run)


FL_simulation_out_ResponseInd <- simulation_ci(data_with_SDFD = live_ref %>% filter(`Testing community` == "Individual"))
# percent_outcomes(FL_simulation_out_ResponseInd$outcomes_by_run)

FL_simulation_out_ResponseComm <- simulation_ci(data_with_SDFD = live_ref %>% filter(`Testing community` == "Community"))
# percent_outcomes(FL_simulation_out_ResponseComm$outcomes_by_run)

FL_simulation_out_ResponsePop <- simulation_ci(data_with_SDFD = live_ref %>% filter(`Testing community` == "Population"))
# percent_outcomes(FL_simulation_out_ResponsePop$outcomes_by_run)


XS_simulation_out_FILow <- simulation_ci(data_with_SDFD = sterile_ref %>% filter(`Fraction inoculum` < 0.251))
# percent_outcomes(XS_simulation_out_FILow$outcomes_by_run)

XS_simulation_out_FIHigh <- simulation_ci(data_with_SDFD = sterile_ref %>% filter(`Fraction inoculum` > 0.25))
# percent_outcomes(XS_simulation_out_FIHigh$outcomes_by_run)

XS_simulation_out_TrainGreenhouse <- simulation_ci(data_with_SDFD = sterile_ref %>% filter(`Training environment` == "Lab"))
# percent_outcomes(XS_simulation_out_TrainGreenhouse$outcomes_by_run)

XS_simulation_out_TrainField <- simulation_ci(data_with_SDFD = sterile_ref %>% filter(`Training environment` == "Field"))
# percent_outcomes(XS_simulation_out_TrainField$outcomes_by_run)


XS_simulation_out_ResponseInd <- simulation_ci(data_with_SDFD = sterile_ref %>% filter(`Testing community` == "Individual"))
# percent_outcomes(XS_simulation_out_ResponseInd$outcomes_by_run)

XS_simulation_out_ResponseComm <- simulation_ci(data_with_SDFD = sterile_ref %>% filter(`Testing community` == "Community"))
# percent_outcomes(XS_simulation_out_ResponseComm$outcomes_by_run)

XS_simulation_out_ResponsePop <- simulation_ci(data_with_SDFD = sterile_ref %>% filter(`Testing community` == "Population"))
# percent_outcomes(XS_simulation_out_ResponsePop$outcomes_by_run)

XS_simulation_out_TrainGreenhouse <- simulation_ci(data_with_SDFD = sterile_ref %>% filter(`Training environment` == "Lab"))
# percent_outcomes(XS_simulation_out_TrainGreenhouse$outcomes_by_run)

XS_simulation_out_TrainField <- simulation_ci(data_with_SDFD = sterile_ref %>% filter(`Training environment` == "Field"))
# percent_outcomes(XS_simulation_out_TrainField$outcomes_by_run)


XS_simulation_out_ResponseInd <- simulation_ci(data_with_SDFD = sterile_ref %>% filter(`Testing community` == "Individual"))
# percent_outcomes(XS_simulation_out_ResponseInd$outcomes_by_run)

XS_simulation_out_ResponseComm <- simulation_ci(data_with_SDFD = sterile_ref %>% filter(`Testing community` == "Community"))
# percent_outcomes(XS_simulation_out_ResponseComm$outcomes_by_run)

XS_simulation_out_ResponsePop <- simulation_ci(data_with_SDFD = sterile_ref %>% filter(`Testing community` == "Population"))
# percent_outcomes(XS_simulation_out_ResponsePop$outcomes_by_run)

XS_simulation_out_SterileAutoclave <- simulation_ci(data_with_SDFD = sterile_ref %>% filter(Study_sterilization_method == "autoclaving"))
# percent_outcomes(XS_simulation_out_SterileAutoclave$outcomes_by_run)

XS_simulation_out_SterileAutoclave <- simulation_ci(data_with_SDFD = sterile_ref %>% filter(Study_sterilization_method == "autoclaving"))
# percent_outcomes(XS_simulation_out_SterileAutoclave$outcomes_by_run)

XS_simulation_out_SterileGamma <- simulation_ci(data_with_SDFD = sterile_ref %>% filter(Study_sterilization_method == "gamma irradiation"))
# percent_outcomes(XS_simulation_out_SterileGamma$outcomes_by_run)

XS_simulation_out_SterileHeat <- simulation_ci(data_with_SDFD = sterile_ref %>% filter(Study_sterilization_method == "heating"))
# percent_outcomes(XS_simulation_out_SterileHeat$outcomes_by_run)

XS_simulation_out_SterileOther <- simulation_ci(data_with_SDFD = sterile_ref %>% filter(!(Study_sterilization_method %in%  
                                                                                         c("autoclaving", "gamma irradiation", "heating"))))
# percent_outcomes(XS_simulation_out_SterileOther$outcomes_by_run)


XS_simulation_out_SterileSoilGreenhouse <- simulation_ci(data_with_SDFD = sterile_ref %>% filter(control_type == "GS"))
# percent_outcomes(XS_simulation_out_SterileSoilGreenhouse$outcomes_by_run)

XS_simulation_out_SterileSoilField <- simulation_ci(data_with_SDFD = sterile_ref %>% filter(control_type == "FS"))
# percent_outcomes(XS_simulation_out_SterileSoilField$outcomes_by_run)

XS_simulation_out_SterileSoilCS <- simulation_ci(data_with_SDFD = sterile_ref %>% filter(control_type == "CS"))
# percent_outcomes(XS_simulation_out_SterileSoilCS$outcomes_by_run)

XS_simulation_out_SterileSoilCmS <- simulation_ci(data_with_SDFD = sterile_ref %>% filter(control_type == "CmS"))
# percent_outcomes(XS_simulation_out_SterileSoilCmS$outcomes_by_run)
#
```

###### **Table of outcomes for live soil reference**

```
# library(kableExtra)
options(kableExtra.latex.load_packages = FALSE)
table_of_things <-
  rbind(c("Overall",(t(unlist(percent_outcomes(live_ref_sims_by_run))))),

      c("Low",t(unlist(percent_outcomes(FL_simulation_out_FILow$outcomes_by_run)))),
      c("High",t(unlist(percent_outcomes(FL_simulation_out_FIHigh$outcomes_by_run)))),

      c("Field",t(unlist(percent_outcomes(FL_simulation_out_TrainField$outcomes_by_run)))),
      c("Greenhouse",t(unlist(percent_outcomes(FL_simulation_out_TrainGreenhouse$outcomes_by_run)))),
      
      c("Individual",t(unlist(percent_outcomes(FL_simulation_out_ResponseInd$outcomes_by_run)))),
      c("Population",t(unlist(percent_outcomes(FL_simulation_out_ResponsePop$outcomes_by_run)))),
      c("Community",t(unlist(percent_outcomes(FL_simulation_out_ResponseComm$outcomes_by_run))))
    
      ) %>%
  as_tibble() %>%
  mutate(category = c("Overall", 
                      "Inoculation fraction", "Inoculation fraction",
                      "Training phase location", "Training phase location",
                      "Response phase growth type", "Response phase growth type", "Response phase growth type")) %>%
  rename("Method" = V1,
         "Exclusion" = V2,
         "Coexistence" = V3,
         "Priority effects" = V4) %>%
  select(category, Method, Coexistence, `Priority effects`, Exclusion) %>%
  kable(format = "latex", booktabs = T,align = "llccc") %>%
  # kable_styling(latex_options = "striped",
  #                           stripe_index = c(2,3, 6,7,8)) %>%
  column_spec(c(1,3,4,5), width = "1.1in")  %>%
  row_spec(c(2:3,6:8) - 1, extra_latex_after = "\\rowcolor{gray!10}") %>%
  collapse_rows(1, latex_hline = "none")
```

```
## Warning: The `x` argument of `as_tibble.matrix()` must have unique column names if `.name_repair` is omitted as of tibble 2.0.0.
## Using compatibility `.name_repair`.
## This warning is displayed once every 8 hours.
## Call `lifecycle::last_lifecycle_warnings()` to see where this warning was generated.
```

```
if(save_figures) {
  saveRDS(object = table_of_things, file = "figures/table_S1.Rds")  
}

# Code to print table in HTML:
rbind(c("Overall",(t(unlist(percent_outcomes(live_ref_sims_by_run))))),
      
      c("Low",t(unlist(percent_outcomes(FL_simulation_out_FILow$outcomes_by_run)))),
      c("High",t(unlist(percent_outcomes(FL_simulation_out_FIHigh$outcomes_by_run)))),
      
      c("Field",t(unlist(percent_outcomes(FL_simulation_out_TrainField$outcomes_by_run)))),
      c("Greenhouse",t(unlist(percent_outcomes(FL_simulation_out_TrainGreenhouse$outcomes_by_run)))),
      
      c("Individual",t(unlist(percent_outcomes(FL_simulation_out_ResponseInd$outcomes_by_run)))),
      c("Population",t(unlist(percent_outcomes(FL_simulation_out_ResponsePop$outcomes_by_run)))),
      c("Community",t(unlist(percent_outcomes(FL_simulation_out_ResponseComm$outcomes_by_run))))
    
      ) %>%
  as_tibble() %>%
  mutate(category = c("Overall", 
                      "Inoculation fraction", "Inoculation fraction",
                      "Training phase location", "Training phase location",
                      "Response phase growth type", "Response phase growth type", "Response phase growth type")) %>%
  rename("Method" = V1,
         "Exclusion" = V2,
         "Coexistence" = V3,
         "Priority effects" = V4) %>%
  select(category, Method, Coexistence, `Priority effects`, Exclusion) %>%
  kable(align = "llccc")
```

| category | Method | Coexistence | Priority effects | Exclusion |
| --- | --- | --- | --- | --- |
| Overall | Overall | 22.2% (16.7%-29.2%) | 6.9% (4.2%-11.1%) | 69.4% (62.5%-76.4%) |
| Inoculation fraction | Low | 17.6% (5.9%-29.4%) | 5.9% (0.0%-17.6%) | 76.5% (58.8%-88.2%) |
| Inoculation fraction | High | 23.6% (18.2%-30.9%) | 7.3% (3.6%-12.7%) | 67.3% (60.0%-76.4%) |
| Training phase location | Field | 28.6% (14.3%-42.9%) | 21.4% (7.1%-28.6%) | 50.0% (35.7%-71.4%) |
| Training phase location | Greenhouse | 22.4% (15.5%-27.6%) | 3.4% (1.7%-8.6%) | 74.1% (67.2%-81.0%) |
| Response phase growth type | Individual | 23.6% (16.4%-30.9%) | 7.3% (3.6%-12.7%) | 69.1% (60.0%-76.4%) |
| Response phase growth type | Population | 0.0% (0.0%-50.0%) | 0.0% (0.0%-50.0%) | 50.0% (0.0%-100.0%) |
| Response phase growth type | Community | 20.0% (6.7%-33.3%) | 6.7% (0.0%-13.3%) | 73.3% (60.0%-86.7%) |

###### **Table of outcomes for sterile soil reference**

```
table_of_things_sterile <-
  rbind(c("Overall",(t(unlist(percent_outcomes(sterile_ref_sims_by_run))))),
        
        c("Low",t(unlist(percent_outcomes(XS_simulation_out_FILow$outcomes_by_run)))),
        c("High",t(unlist(percent_outcomes(XS_simulation_out_FIHigh$outcomes_by_run)))),
        
        c("Field",t(unlist(percent_outcomes(XS_simulation_out_TrainField$outcomes_by_run)))),
        c("Greenhouse",t(unlist(percent_outcomes(XS_simulation_out_TrainGreenhouse$outcomes_by_run)))),
        
        c("Individual",t(unlist(percent_outcomes(XS_simulation_out_ResponseInd$outcomes_by_run)))),
        c("Population",t(unlist(percent_outcomes(XS_simulation_out_ResponsePop$outcomes_by_run)))),
        c("Community",t(unlist(percent_outcomes(XS_simulation_out_ResponseComm$outcomes_by_run)))),
        
        c("Autoclaving",t(unlist(percent_outcomes(XS_simulation_out_SterileAutoclave$outcomes_by_run)))),
        c("Gamma irradiation",t(unlist(percent_outcomes(XS_simulation_out_SterileGamma$outcomes_by_run)))),
        c("Heat",t(unlist(percent_outcomes(XS_simulation_out_SterileHeat$outcomes_by_run)))),
        c("Other",t(unlist(percent_outcomes(XS_simulation_out_SterileOther$outcomes_by_run)))),

        c("Grenhouse soil",t(unlist(percent_outcomes(XS_simulation_out_SterileSoilGreenhouse$outcomes_by_run)))),
        c("Field soil",t(unlist(percent_outcomes(XS_simulation_out_SterileSoilGreenhouse$outcomes_by_run)))),
        c("Conditioned soil (species-specific)",t(unlist(percent_outcomes(XS_simulation_out_SterileSoilCS$outcomes_by_run)))),
        c("Conditioned soil (mixed)",t(unlist(percent_outcomes(XS_simulation_out_SterileSoilCS$outcomes_by_run))))
        
        
  ) %>%
  as_tibble() %>%
  mutate(category = c("Overall", 
                      "Inoculation fraction", "Inoculation fraction",
                      "Training phase location", "Training phase location",
                      "Response phase growth type", "Response phase growth type", "Response phase growth type",
                      rep("Sterilization method", 4),
                      rep("Type of soil sterilized", 4))) %>%
  rename("Method" = V1,
         "Exclusion" = V2,
         "Coexistence" = V3,
         "Priority effects" = V4) %>%
  select(category, Method, Coexistence, `Priority effects`, Exclusion) %>%
  kable(format = "latex", booktabs = T,align = "llccc") %>%
  # kable_styling(latex_options = "striped",
  #                           stripe_index = c(2,3, 6,7,8)) %>%
  column_spec(c(1,2,3,4,5), width = "1.1in")  %>%
  row_spec(c(2:3,6:8, 13:16) - 1, extra_latex_after = "\\rowcolor{gray!10}") %>%
  collapse_rows(1, latex_hline = "none") 

if(save_figures) {
  saveRDS(object = table_of_things_sterile, file = "figures/table_S2.Rds")  
}


rbind(c("Overall",(t(unlist(percent_outcomes(sterile_ref_sims_by_run))))),
      
      c("Low",t(unlist(percent_outcomes(XS_simulation_out_FILow$outcomes_by_run)))),
      c("High",t(unlist(percent_outcomes(XS_simulation_out_FIHigh$outcomes_by_run)))),
      
      c("Field",t(unlist(percent_outcomes(XS_simulation_out_TrainField$outcomes_by_run)))),
      c("Greenhouse",t(unlist(percent_outcomes(XS_simulation_out_TrainGreenhouse$outcomes_by_run)))),
      
      c("Individual",t(unlist(percent_outcomes(XS_simulation_out_ResponseInd$outcomes_by_run)))),
      c("Population",t(unlist(percent_outcomes(XS_simulation_out_ResponsePop$outcomes_by_run)))),
      c("Community",t(unlist(percent_outcomes(XS_simulation_out_ResponseComm$outcomes_by_run)))),
      
      c("Autoclaving",t(unlist(percent_outcomes(XS_simulation_out_SterileAutoclave$outcomes_by_run)))),
      c("Gamma irradiation",t(unlist(percent_outcomes(XS_simulation_out_SterileGamma$outcomes_by_run)))),
      c("Heat",t(unlist(percent_outcomes(XS_simulation_out_SterileHeat$outcomes_by_run)))),
      c("Other",t(unlist(percent_outcomes(XS_simulation_out_SterileOther$outcomes_by_run)))),
      
      c("Grenhouse soil",t(unlist(percent_outcomes(XS_simulation_out_SterileSoilGreenhouse$outcomes_by_run)))),
      c("Field soil",t(unlist(percent_outcomes(XS_simulation_out_SterileSoilGreenhouse$outcomes_by_run)))),
      c("Conditioned soil (species-specific)",t(unlist(percent_outcomes(XS_simulation_out_SterileSoilCS$outcomes_by_run)))),
      c("Conditioned soil (mixed)",t(unlist(percent_outcomes(XS_simulation_out_SterileSoilCS$outcomes_by_run))))
      
      
) %>%
  as_tibble() %>%
  mutate(category = c("Overall", 
                      "Inoculation fraction", "Inoculation fraction",
                      "Training phase location", "Training phase location",
                      "Response phase growth type", "Response phase growth type", "Response phase growth type",
                      rep("Sterilization method", 4),
                      rep("Type of soil sterilized", 4))) %>%
  rename("Method" = V1,
         "Exclusion" = V2,
         "Coexistence" = V3,
         "Priority effects" = V4) %>%
  select(category, Method, Coexistence, `Priority effects`, Exclusion) %>%
  kable(booktabs = T, align = "llccc")
```

| category | Method | Coexistence | Priority effects | Exclusion |
| --- | --- | --- | --- | --- |
| Overall | Overall | 12.8% (10.8%-14.8%) | 6.1% (4.5%-7.6%) | 81.2% (78.7%-83.6%) |
| Inoculation fraction | Low | 11.6% (9.4%-14.0%) | 6.7% (4.9%-8.5%) | 81.8% (79.0%-84.5%) |
| Inoculation fraction | High | 16.2% (12.8%-20.5%) | 4.3% (1.7%-6.8%) | 79.5% (75.2%-83.8%) |
| Training phase location | Field | 12.4% (9.6%-15.8%) | 7.3% (4.5%-10.2%) | 79.7% (75.7%-83.6%) |
| Training phase location | Greenhouse | 9.4% (7.2%-11.5%) | 4.7% (2.6%-6.8%) | 86.0% (83.0%-88.9%) |
| Response phase growth type | Individual | 21.2% (17.5%-24.9%) | 7.9% (5.3%-11.1%) | 70.9% (66.1%-75.1%) |
| Response phase growth type | Population | 8.1% (2.7%-13.5%) | 5.4% (2.6%-13.5%) | 86.5% (78.4%-94.6%) |
| Response phase growth type | Community | 6.4% (4.5%-8.6%) | 4.1% (2.3%-5.9%) | 89.5% (86.8%-92.3%) |
| Sterilization method | Autoclaving | 22.9% (17.8%-28.0%) | 9.3% (5.9%-13.6%) | 67.8% (61.9%-73.7%) |
| Sterilization method | Gamma irradiation | 14.8% (11.5%-18.9%) | 4.1% (1.6%-7.4%) | 80.3% (76.2%-84.4%) |
| Sterilization method | Heat | 4.2% (2.1%-6.3%) | 4.2% (2.6%-6.9%) | 91.5% (88.4%-94.2%) |
| Sterilization method | Other | 23.5% (11.8%-35.3%) | 11.8% (0.0%-23.5%) | 70.6% (52.9%-82.4%) |
| Type of soil sterilized | Grenhouse soil | 6.0% (3.7%-8.3%) | 5.1% (2.8%-7.4%) | 88.9% (86.1%-92.1%) |
| Type of soil sterilized | Field soil | 6.0% (3.7%-8.3%) | 5.1% (2.8%-7.4%) | 88.9% (86.1%-92.1%) |
| Type of soil sterilized | Conditioned soil (species-specific) | 16.9% (13.3%-20.5%) | 6.7% (4.1%-9.2%) | 76.4% (72.8%-80.5%) |
| Type of soil sterilized | Conditioned soil (mixed) | 16.9% (13.3%-20.5%) | 6.7% (4.1%-9.2%) | 76.4% (72.8%-80.5%) |

##### Meta-analysis approach

Here we do separate meta-analysis models for species pairs each each dataset with stabilizing or destabilizing effects of soil microbes

##### Run meta-analysis for live ref soil

```
live_ref_long_q2 <- 
  live_ref_long %>% 
  # Get rid of the rows where SD < 0
  filter(set == "FD" | (set == "SD" & mean_value > 0)) %>%
  # Eliminate the species pairs whose SDs were eliminated in the previous step
  group_by(Experiment_SpPair) %>% 
  mutate(n_rows = n()) %>% 
  filter(n_rows > 1) %>% 
  # Clean up the dataset
  select(-n_rows)

q2.lr.0.5 <- run_metamod(which_df = live_ref_long_q2, 
                         which_response_column = "absolute_FDSD_effects", 
                         moderators = " ~ (set-1)",
                         impute_cov_r = 0.5)
```

**Live reference soil, species pairs with stabilizing effects of soil microbes**

```
# check if different from zero
coef_test(q2.lr.0.5, vcov = "CR2") %>% 
  kable() %>%
  kable_styling(full_width = 0)
```

|  | beta | SE | tstat | df | p\_Satt |
| --- | --- | --- | --- | --- | --- |
| setFD | 0.7203153 | 0.0966905 | 7.449697 | 10.35514 | 0.0000180 |
| setSD | 0.4385484 | 0.1093915 | 4.008982 | 11.20481 | 0.0019813 |

```
# check if values are different from one another
Wald_test(q2.lr.0.5, constraints = constrain_equal(1:2),  vcov = "CR2") %>% 
  kable() %>%
  kable_styling(full_width = 0)
```

| test | Fstat | delta | df\_num | df\_denom | p\_val |
| --- | --- | --- | --- | --- | --- |
| HTZ | 22.05369 | 1 | 1 | 6.662667 | 0.0025311 |

```
live_ref_long_q2_destab <- 
  live_ref_long %>% 
  # Get rid of the rows where SD < 0
  filter(set == "FD" | (set == "SD" & mean_value < 0)) %>%
  # Eliminate the species pairs whose SDs were eliminated in the previous step
  group_by(Experiment_SpPair) %>% 
  mutate(n_rows = n()) %>% 
  filter(n_rows > 1) %>% 
  # Clean up the dataset
  select(-n_rows)

q2.lr.0.5.destab <- run_metamod(which_df = live_ref_long_q2_destab, 
                         which_response_column = "absolute_FDSD_effects", 
                         moderators = " ~ (set-1)",
                         impute_cov_r = 0.5)
```

**Live reference soil, species pairs with destabilizing effects of soil microbes**

```
# check if different from zero
coef_test(q2.lr.0.5.destab, vcov = "CR2") %>% 
  kable() %>%
  kable_styling(full_width = 0)
```

|  | beta | SE | tstat | df | p\_Satt |
| --- | --- | --- | --- | --- | --- |
| setFD | 0.5294017 | 0.1815242 | 2.916425 | 6.010139 | 0.0267034 |
| setSD | 0.2808761 | 0.0578024 | 4.859245 | 5.144896 | 0.0042949 |

```
# check if values are different from one another
Wald_test(q2.lr.0.5.destab, constraints = constrain_equal(1:2),  vcov = "CR2") %>% 
  kable() %>%
  kable_styling(full_width = 0)
```

| test | Fstat | delta | df\_num | df\_denom | p\_val |
| --- | --- | --- | --- | --- | --- |
| HTZ | 2.708292 | 1 | 1 | 5.720508 | 0.1533417 |

###### Run meta-analysis for sterile ref soil

```
sterile_ref_long_q2 <- 
  sterile_ref_long %>% 
  # Get rid of the rows where SD < 0
  filter(set == "FD" | (set == "SD" & mean_value > 0)) %>%
  # Eliminate the species pairs whose SDs were eliminated in the previous step
  group_by(Experiment_SpPair) %>% 
  mutate(n_rows = n()) %>% 
  filter(n_rows > 1) %>% 
  # Clean up the dataset
  select(-n_rows)

q2.xs.0.5 <- run_metamod(which_df = sterile_ref_long_q2, 
                         which_response_column = "absolute_FDSD_effects", 
                         moderators = " ~ (set-1)",
                         impute_cov_r = 0.5)
```

**Sterile reference soils, species pairs with stabilizing effects of soil microbes**

```
# check if different from zero
coef_test(q2.xs.0.5, vcov = "CR2") %>% 
  kable() %>%
  kable_styling(full_width = F)
```

|  | beta | SE | tstat | df | p\_Satt |
| --- | --- | --- | --- | --- | --- |
| setFD | 0.7866931 | 0.1176843 | 6.684776 | 40.04465 | 1e-07 |
| setSD | 0.1992079 | 0.0282220 | 7.058601 | 45.02829 | 0e+00 |

```
# check if values are different from one another
Wald_test(q2.xs.0.5, constraints = constrain_equal(1:2),  vcov = "CR2") %>% 
  kable() %>%
  kable_styling(full_width = F)
```

| test | Fstat | delta | df\_num | df\_denom | p\_val |
| --- | --- | --- | --- | --- | --- |
| HTZ | 25.85547 | 1 | 1 | 39.92169 | 9.1e-06 |

```
sterile_ref_long_q2_destab <- 
  sterile_ref_long %>% 
  # Get rid of the rows where SD < 0
  filter(set == "FD" | (set == "SD" & mean_value < 0)) %>%
  # Eliminate the species pairs whose SDs were eliminated in the previous step
  group_by(Experiment_SpPair) %>% 
  mutate(n_rows = n()) %>% 
  filter(n_rows > 1) %>% 
  # Clean up the dataset
  select(-n_rows)

q2.xs.0.5.destab <- run_metamod(which_df = sterile_ref_long_q2_destab, 
                         which_response_column = "absolute_FDSD_effects", 
                         moderators = " ~ (set-1)",
                         impute_cov_r = 0.5)
```

**Sterile reference soil, species pairs with destabilizing effects of soil microbes**

```
# check if different from zero
coef_test(q2.xs.0.5.destab, vcov = "CR2") %>% 
  kable() %>%
  kable_styling(full_width = F)
```

|  | beta | SE | tstat | df | p\_Satt |
| --- | --- | --- | --- | --- | --- |
| setFD | 0.8935225 | 0.1319193 | 6.773249 | 34.87081 | 1e-07 |
| setSD | 0.2251813 | 0.0340159 | 6.619884 | 29.69725 | 3e-07 |

```
# check if values are different from one another
Wald_test(q2.xs.0.5.destab, constraints = constrain_equal(1:2),  vcov = "CR2") %>% 
  kable() %>%
  kable_styling(full_width = F)
```

| test | Fstat | delta | df\_num | df\_denom | p\_val |
| --- | --- | --- | --- | --- | --- |
| HTZ | 36.313 | 1 | 1 | 34.04953 | 8e-07 |

###### Make plots

**For pairs with stabilizing effects (both reference types)**

```
q2b_plots_df_stabilizing <- bind_rows(
  data.frame(conf_int(q2.xs.0.5, vcov = "CR2")),
  data.frame(conf_int(q2.lr.0.5, vcov = "CR2"))
)

q2b_plots_df_stabilizing <- 
  q2b_plots_df_stabilizing %>%
  mutate(yval = c(1.2, 0.8, 2.2, 1.8),
         which_source = rep(c("XS", "LR"), each = 2),
         effect_type = rep(c("Fitness difference", 
                 "(De)stabilization"), 2))

q2.stab.prediction_intervals <- bind_rows(data.frame(predict.rma(q2.xs.0.5)[1:2,]),
                                  data.frame(predict.rma(q2.lr.0.5)[1:2,])) %>%
  mutate(which_source = rep(c("XS", "LR"), each = 2),
         effect_type = c("(De)stabilization", "Fitness difference", "(De)stabilization", "Fitness difference"))

q2b_plots_df_stabilizing <- left_join(x = q2b_plots_df_stabilizing, y = q2.stab.prediction_intervals, by = c("effect_type", "which_source"))


(q2_plot_stabilizing <- 
    ggplot(q2b_plots_df_stabilizing) +
    ggbeeswarm::geom_quasirandom(data = sterile_ref_long_q2,
                                 aes(x = absolute_FDSD_effects, y = yval, size = scale, color = effect_type),
                                 groupOnX = F, alpha = 1, pch = 21, fill = "transparent", stroke = 0.25) +
    ggbeeswarm::geom_quasirandom(data = live_ref_long_q2,
                                 aes(x = absolute_FDSD_effects, y = yval, size = scale, color = effect_type),
                                 groupOnX = F, alpha = 1, pch = 21, fill = "transparent", stroke = 0.5) +
    geom_point(aes(x = beta, y = yval, fill = effect_type),
               size = 5, stroke = 1.25, shape = 21) + 
    geom_errorbarh(aes(y = yval, xmin = CI_L, xmax = CI_U), height = 0, size = 0.8) +
    geom_hline(yintercept = 1.5, size = .25) + 
    geom_vline(xintercept = 0, linetype = 2, size = .5, color = "grey25") +
    annotate("text", x = Inf, y = 1, label = paste0("K = ", nrow(sterile_ref_long_q2)/2, 
                                                    " species pairs\n (",
                                                    length(unique(sterile_ref_long_q2$Experiment)),
                                                    " experiments)"), hjust = 1, size = 4) +
    annotate("text", x = Inf, y = 2, label = paste0("K = ", nrow(live_ref_long_q2)/2, 
                                                    " species pairs\n (",
                                                    length(unique(live_ref_long_q2$Experiment)),
                                                    " experiments)"), hjust = 1, size = 4) + 
  scale_fill_manual(name = "", values = rev(c(alpha("#009E73", 0.6), alpha("#CC79A7", 0.75)))) +
  scale_color_manual(name = "", values = rev(c("#009E73", "#CC79A7", "#009E73","#CC79A7"))) +
  scale_y_continuous(name = "", breaks = c(1,2), labels = c("Sterile\nreference\nsoil","Live\nreference\nsoil")) +
  scale_x_continuous(limits = c(-1.1, 5.05), breaks = c(0, 2,4)) +
  scale_size_continuous(name = "Precision (1/SE)", breaks = c(10, 20, 30)) + 
  xlab("Effect size") + 
  theme_plots() +
  
  annotate("segment", x = 2.15, xend = Inf, y = c(1.5, 1.7), yend = c(1.5,1.7), size = .25) +
  annotate("segment", x = c(2.15, Inf), xend = c(2.15, Inf), y = 1.5, yend = 1.7, size = .25)  + 
  geom_point(aes(x = 2.27, y = 1.65), size = 4, shape = 21, fill = c("#009E73")) + 
  geom_point(aes(x = 2.27, y = 1.55), size = 4, shape = 21, fill = c("#CC79A7")) + 
  annotate("text", x = 2.4, y = 1.65, label = "Fitness difference", hjust = 0, size = 4) +
  annotate("text", x = 2.4, y = 1.55, label = "Stabilizing effects", hjust = 0, size = 4) +  
  
  guides(fill = "none", color = "none") + 
  labs(title = "(A) Species pairs with stabilizing effects of soil microbes") + 
  theme(axis.text.y = element_text(color = "black", size = 11),
        legend.position= c(1, 0), 
        legend.direction = "horizontal",
        legend.justification = c(1, 0),
        plot.title.position = "plot",
        plot.title = element_text(face = "bold"))
)
```

**For pairs with destabilizing effects (both reference types)**

```
q2b_plots_df_destabilizing <- bind_rows(
  data.frame(conf_int(q2.xs.0.5.destab, vcov = "CR2")),
  data.frame(conf_int(q2.lr.0.5.destab, vcov = "CR2"))
)

q2b_plots_df_destabilizing <- 
  q2b_plots_df_destabilizing %>%
  mutate(yval = c(1.2, 0.8, 2.2, 1.8),
         which_source = rep(c("XS", "LR"), each = 2),
         effect_type = rep(c("Fitness difference", 
                 "(De)stabilization"), 2))

q2.destab.prediction_intervals <- bind_rows(data.frame(predict.rma(q2.xs.0.5.destab)[1:2,]),
                                          data.frame(predict.rma(q2.lr.0.5.destab)[1:2,])) %>%
  mutate(which_source = rep(c("XS", "LR"), each = 2),
         effect_type = c("(De)stabilization", "Fitness difference", "(De)stabilization", "Fitness difference"))

q2b_plots_df_destabilizing <- left_join(x = q2b_plots_df_destabilizing, 
                                        y = q2.destab.prediction_intervals, 
                                        by = c("effect_type", "which_source"))

(q2_plot_destabilizing <- ggplot(q2b_plots_df_destabilizing) +
  ggbeeswarm::geom_quasirandom(data = sterile_ref_long_q2_destab,
                               aes(x = absolute_FDSD_effects, y = yval, size = scale, color = effect_type),
                               groupOnX = F, alpha = 1, pch = 21, fill = "transparent", stroke = 0.25) +
  ggbeeswarm::geom_quasirandom(data = live_ref_long_q2_destab,
                               aes(x = absolute_FDSD_effects, y = yval, size = scale, color = effect_type),
                               groupOnX = F, alpha = 1, pch = 21, fill = "transparent", stroke = 0.5) +
  geom_point(aes(x = beta, y = yval, fill = effect_type),
             size = 5, stroke = 1.25, shape = 21) + 
  geom_errorbarh(aes(y = yval, xmin = CI_L, xmax = CI_U), height = 0, size = 0.8) +
  geom_hline(yintercept = 1.5, size = .25) + 
  geom_vline(xintercept = 0, linetype = 2, size = .5, color = "grey25") +
    annotate("text", x = Inf, y = 1, label = paste0("K = ", nrow(sterile_ref_long_q2_destab)/2, 
                                                    " species pairs\n (",
                                                    length(unique(sterile_ref_long_q2_destab$Experiment)),
                                                    " experiments)"), hjust = 1, size = 4) +
    annotate("text", x = Inf, y = 2, label = paste0("K = ", nrow(live_ref_long_q2_destab)/2, 
                                                    " species pairs\n (",
                                                    length(unique(live_ref_long_q2_destab$Experiment)),
                                                    " experiments)"), hjust = 1, size = 4) + 

  scale_fill_manual(name = "", values = rev(c(alpha("#009E73", 0.6), alpha("#CC79A7", 0.75)))) +
  scale_color_manual(name = "", values = rev(c("#009E73", "#CC79A7", "#009E73","#CC79A7"))) +
  scale_y_continuous(name = "", breaks = c(1,2), labels = c("Sterile\nreference\nsoil", "Live\nreference\nsoil")) +
  scale_x_continuous(limits = c(-1, 5.05), breaks = c(0, 2,4)) +
  scale_size_continuous(name = "Precision (1/SE)", breaks = c(10, 20, 30)) + 
  xlab("Effect size") + 
  theme_plots() +
    
    annotate("segment", x = 1.9, xend = Inf, y = c(1.5, 1.7), yend = c(1.5,1.7), size = .25) +
    annotate("segment", x = c(1.9, Inf), xend = c(1.9, Inf), y = 1.5, yend = 1.7, size = .25)  + 
    geom_point(aes(x = 2.1, y = 1.65), size = 4, shape = 21, fill = c("#009E73")) + 
    geom_point(aes(x = 2.1, y = 1.55), size = 4, shape = 21, fill = c("#CC79A7")) + 
    annotate("text", x = 2.25, y = 1.65, label = "Fitness difference", hjust = 0, size = 4) +
    annotate("text", x = 2.25, y = 1.55, label = "Destabilizing effects", hjust = 0, size = 4) +  

  guides(fill = "none", color = "none") + 
  labs(title = "(B) Species pairs with destabilizing effects of soil microbes") + 
  theme(axis.text.y = element_text(color = "black", size = 11),
        legend.position= c(1, 0), 
        legend.direction = "horizontal",
        legend.justification = c(1, 0),
        plot.title.position = "plot",
        plot.title = element_text(face = "bold"))
)
```

*For the supplemental figures we put these two panels together into one figure, see code below for details.*

```
q2b_plot <- q2_plot_stabilizing + q2_plot_destabilizing
# if(save_figures = T) {
  ggsave("figures/q2_meta-plot.png", width = 11, height = 5)
# }
```

#### Supplemental analyses

##### Univariate version of the Q1 models

```
q1.lr.univariate.fd <- rma.mv(yi = absolute_FDSD_effects, V = var_value, 
                              random = ~ 1|Experiment, 
                              data = live_ref_long %>% filter(set == "FD"))

q1.lr.univariate.sd <- rma.mv(yi = absolute_FDSD_effects, V = var_value, 
                              random = ~ 1|Experiment, 
                              data = live_ref_long %>% filter(set == "SD"))


q1.xs.univariate.fd <- rma.mv(yi = absolute_FDSD_effects, V = var_value, 
                              random = ~ 1|Experiment, 
                              data = sterile_ref_long %>% filter(set == "FD"))
q1.xs.univariate.sd <- rma.mv(yi = absolute_FDSD_effects, V = var_value, 
                              random = ~ 1|Experiment, 
                              data = sterile_ref_long %>% filter(set == "SD"))


q1_multi_v_uni <-
  bind_rows(
  data.frame(conf_int(q1.xs.0.5, "CR2")),
  data.frame(conf_int(q1.xs.univariate.fd, "CR2")),
  data.frame(conf_int(q1.xs.univariate.sd, "CR2")),
  data.frame(conf_int(q1.lr.0.5, "CR2")),
  data.frame(conf_int(q1.lr.univariate.fd, "CR2")),
  data.frame(conf_int(q1.lr.univariate.sd, "CR2"))

) %>%
  mutate(which_analysis = rep(c("Multivariate", "Multivariate", "Univariate", "Univariate"),2),
         which_metric = rep(c("Fitness difference", "(De)stabilization"), 4),
         which_ref = c("Sterile", "Sterile", "Sterile", "Sterile", "Live", "Live", "Live", "Live"),
         yval = c(2.25, 1.85, 2.15, 1.75,
                  1.25, 0.85, 1.15, 0.75))

q1_multi_v_uni %>% 
  as_tibble() %>%
  kable %>%
  kable_styling(full_width = F)
```

| beta | SE | df | CI\_L | CI\_U | which\_analysis | which\_metric | which\_ref | yval |
| --- | --- | --- | --- | --- | --- | --- | --- | --- |
| 0.7481241 | 0.0998507 | 52.28616 | 0.5477851 | 0.9484631 | Multivariate | Fitness difference | Sterile | 2.25 |
| 0.2129726 | 0.0245918 | 54.80079 | 0.1636856 | 0.2622596 | Multivariate | (De)stabilization | Sterile | 1.85 |
| 0.6118938 | 0.0755687 | 64.60269 | 0.4609551 | 0.7628325 | Univariate | Fitness difference | Sterile | 2.15 |
| 0.2010285 | 0.0224690 | 58.74263 | 0.1560641 | 0.2459929 | Univariate | (De)stabilization | Sterile | 1.75 |
| 0.6308658 | 0.0909004 | 14.07145 | 0.4359967 | 0.8257348 | Multivariate | Fitness difference | Live | 1.25 |
| 0.3638120 | 0.0721101 | 13.98695 | 0.2091378 | 0.5184863 | Multivariate | (De)stabilization | Live | 0.85 |
| 0.5965602 | 0.0977646 | 15.64835 | 0.3889292 | 0.8041911 | Univariate | Fitness difference | Live | 1.15 |
| 0.3649092 | 0.0631866 | 15.63165 | 0.2307026 | 0.4991157 | Univariate | (De)stabilization | Live | 0.75 |

Now, we can make a plot:

```
q1_multi_v_uni_plot <-
  ggplot(q1_multi_v_uni) +
  # facet_wrap(.~which_ref) + 
  geom_errorbarh(aes(xmin = CI_L, xmax = CI_U, y = yval), height = 0, size = 0.25) + 
  geom_point(aes(x = beta, y = yval, color = which_metric, shape = which_analysis), size = 4) + 
  scale_color_manual(name = "Effect size", values = c("#CC79A7", "#009E73")) +
  scale_shape_manual(name = "Analysis type", values = c(17,19)) + 
  ylab("") + xlab("Effect size") + 
  scale_y_continuous(breaks = c(1,2), labels = c("Live\nreference\nsoil", "Sterile\nreference\nsoil")) + 
  scale_x_continuous(limits = c(-0.1, 1)) +
  geom_hline(yintercept = 1.5) + 
    geom_vline(xintercept = 0, linetype = 2) + 

  theme_plots() +
  theme(axis.text.y = element_text(size = 12, color = "black"),
        # legend.position = "top",
        legend.direction = "vertical")

q1_multi_v_uni_plot
```

```
if(save_figures) {
  ggsave("figures/q1_Uni_Vs_Multivariate.png", 
         q1_multi_v_uni_plot, height = 3, width = 6)
}
```

##### Univariate analysis of the strength of (de)stabilization

Instead of analyzing the *magnitude* of the (de)stabilizing effect (by taking the absolute value), here we analyze the actual value of the (de)stabilization metric. This is akin to the meta-analysis model of \(I\_S\) in Crawford et al. (2019)’s paper.

```
# Univariate analysis of the (De)stabilization metric

combined <- bind_rows(live_ref, sterile_ref)
combined_SD_efect <- rma.mv(mean_SD, V = var_SD, 
                            random = list(~1|Experiment, 
                                          ~1|species_pair), 
                            data = combined)

combined_SD_efect
```

```
## 
## Multivariate Meta-Analysis Model (k = 518; method: REML)
## 
## Variance Components:
## 
##             estim    sqrt  nlvls  fixed        factor 
## sigma^2.1  0.0531  0.2304     81     no    Experiment 
## sigma^2.2  0.0837  0.2893    263     no  species_pair 
## 
## Test for Heterogeneity:
## Q(df = 517) = 2782.1503, p-val < .0001
## 
## Model Results:
## 
## estimate      se    zval    pval   ci.lb   ci.ub 
##   0.1000  0.0381  2.6259  0.0086  0.0254  0.1746  ** 
## 
## ---
## Signif. codes:  0 '***' 0.001 '**' 0.01 '*' 0.05 '.' 0.1 ' ' 1
```

Now we can make an orchard plot of the (de)stabilization metric.

```
combined$Precision <- (1/sqrt(combined$var_SD))

overall_stabilization_plot <- 
  ggplot(combined) + 
  ggbeeswarm::geom_beeswarm(aes(x = mean_SD,y = 0, size = Precision),
                            groupOnX = F, alpha = 1, stroke = 0.25, fill = "transparent",
                            shape = 21, color = "#CC79A7") +
  geom_point(aes(y = 0, x = combined_SD_efect$b), shape = 21, 
             size = 5, fill = "#CC79A7", stroke = 1.25) + 
  geom_errorbarh(aes(y = 0, xmin = combined_SD_efect$ci.lb, 
                     xmax = combined_SD_efect$ci.ub), height = 0) +
  geom_vline(xintercept = 0, linetype = "dashed") +
  
  xlab("(De)stabilization") + 
  ylab("") + 
  scale_y_continuous(breaks = c(), limits = c(-0.5, 0.5)) +
  theme_plots() +
  annotate("text", x = 0.1, y = -Inf, hjust = 0, vjust = -1,
           label = "stabilizing", fontface = "italic", color = "grey25") +
  annotate("text", x = -0.1, y = -Inf, hjust = 1, vjust = -1, 
           label = "destabilizing", fontface = "italic", color = "grey25") +
  annotate("segment", x = 0.1, xend = 0.5, y = -0.5, yend = -0.5, 
           colour = "grey25", size = 0.5, arrow = arrow(length = unit(0.075, "inches"))) + 
  annotate("segment", x = -0.1, xend = -0.5, y = -0.5, yend = -0.5, 
           colour = "grey25", size = 0.5, arrow = arrow(length = unit(0.075, "inches"))) +
  
  theme(legend.position = c(1,0),
        legend.justification = c(1, 0),
        legend.direction = "horizontal")

overall_stabilization_plot
```

```
if(save_figures) {
  ggsave("figures/overall_stabilization_plot.png", 
         overall_stabilization_plot, width = 9, height = 2)
}
```

##### Evaluate sensitivity of Q1 models to the imputed value of within-experient correlation

Here we run models as in the main text, but we impute covariance matrices with assumed within-experiment (de)stabilization-fitness difference correlations of -0.5, 0.1, or 0.9. (The main text models are run with assumed correlations of 0.5.). Expand code below to see details.

```
q1.lr.0.1 <- run_metamod(which_df = live_ref_long, 
                         which_response_column = "absolute_FDSD_effects",
                         moderators = " ~ (set-1)",
                         impute_cov_r = 0.1)
q1.lr.0.9 <- run_metamod(which_df = live_ref_long, 
                         which_response_column = "absolute_FDSD_effects",
                         moderators = " ~ (set-1)",
                         impute_cov_r = 0.9)
q1.lr.neg.0.5 <- run_metamod(which_df = live_ref_long, 
                         which_response_column = "absolute_FDSD_effects",
                         moderators = " ~ (set-1)",
                         impute_cov_r = -0.5)


q1.xs.0.1 <- run_metamod(which_df = sterile_ref_long, 
                         which_response_column = "absolute_FDSD_effects", 
                         moderators = " ~ (set-1)",
                         impute_cov_r = 0.1)
q1.xs.0.9 <- run_metamod(which_df = sterile_ref_long, 
                         which_response_column = "absolute_FDSD_effects", 
                         moderators = " ~ (set-1)",
                         impute_cov_r = 0.9)
q1.xs.neg.0.5 <- run_metamod(which_df = sterile_ref_long, 
                         which_response_column = "absolute_FDSD_effects", 
                         moderators = " ~ (set-1)",
                         impute_cov_r = -0.5)

varying_imputed_r <- 
  bind_rows(data.frame(conf_int(q1.xs.neg.0.5, vcov = "CR2")),
            data.frame(conf_int(q1.xs.0.1, vcov = "CR2")),
            data.frame(conf_int(q1.xs.0.5, vcov = "CR2")),
            data.frame(conf_int(q1.xs.0.9, vcov = "CR2")),
            
            data.frame(conf_int(q1.lr.neg.0.5, vcov = "CR2")),
            data.frame(conf_int(q1.lr.0.1, vcov = "CR2")),
            data.frame(conf_int(q1.lr.0.5, vcov = "CR2")),
            data.frame(conf_int(q1.lr.0.9, vcov = "CR2"))) %>%
  mutate(imputed_cov = rep(rep(c(-0.5, 0.1, 0.5, 0.9), each = 2),2),
         effect_type = rep(rep(c("Fitness difference","(De)stabilization"),4),2),
         yval = c(2.2,1.8, 2.15,1.75, 2.1,1.7, 2.05,1.65,
                  1.2,0.8, 1.15,0.75, 1.1,0.7, 1.05,0.65))

varying_imputed_r_plot <- 
  ggplot(varying_imputed_r) +
  geom_point(aes(x = beta, y = yval, shape = as.factor(imputed_cov),
                 color = effect_type), size = 4) +
  geom_errorbarh(aes(xmin = CI_L, xmax = CI_U, y = yval), height = 0) +
  scale_color_manual(name = "Effect size", values = c("#CC79A7", "#009E73"))+
  scale_y_continuous(name = "", breaks = c(1,2),
                     labels = c("Live\nreference\nsoil",
                                "Sterile\nreference\nsoil")) +
  geom_hline(yintercept = 1.5) + 
  scale_shape_manual(name = "Assumed correlation", values = c(15,17,18,19)) +
  theme_plots() + 
  xlab("Effect size") +
  xlim(c(-.1, 1)) +
  geom_vline(xintercept = 0, linetype = 2) +
  theme(axis.text = element_text(color = "black", size = 12))

varying_imputed_r_plot
```

```
if(save_figures){
  ggsave("figures/varying_imputed_r.png", 
         varying_imputed_r_plot, height = 4, width = 7)
}
```

#### Make conceptual figures

*First, assume negative values for all \(m\) terms*

```
psf_model <- function(time, init, params) {
  with (as.list(c(time, init, params)), {
    # description of parameters
    
    # Differential equations
    dpA <- pA*(1-pA)*((m1A-m1B)*pAlpha + (m2A-m2B)*(1-pAlpha))
    dpAlpha <- pAlpha*(1-pAlpha)*(pA-v*(1-pA))
    
    # Return dN1 and dN2
    return(list(c(dpA, dpAlpha)))
  })
}

metrics <- function(params) {
  IS <- with(as.list(params), {m1A - m2A - m1B + m2B})
  FD <- with(as.list(params), {(1/2)*(m1A+m2A) - (1/2)*(m1B+m2B)})
  SD <- (-1/2)*IS
  return(c(IS = IS, SD = SD, FD = FD))
}

params_coex = c(m1A = -.85, m1B = -.4, m2A = -.1, m2B = -.75, v = 1)
params_excl = c(m1A = -.55, m1B = -.7, m2A = -.1, m2B = -.85, v = 1)
params_priority = c(m1A = -0.2, m1B = -0.5, m2A = -0.8, m2B = -0.15, v = 1)
params_excl2 = c(m1A = -.1, m1B = -0.9, m2A = -.2, m2B = -.55, v = 1)
time <- seq(0, 75)
init = c(pA = 0.3, pAlpha = 0.3)

out_coex <- data.frame(deSolve::ode(func = psf_model,
                                    y = init, times = time, parms = params_coex)) 
out_excl <- data.frame(deSolve::ode(func = psf_model,
                                    y = init, times = time, parms = params_excl)) 
out_priority <- data.frame(deSolve::ode(func = psf_model,
                                        y = init, times = time, parms = params_priority)) 

out_priority2 <- data.frame(deSolve::ode(func = psf_model,
                                         y = c(pA = 0.7, pAlpha = 0.7), times = time, parms = params_priority)) 
out_excl2 <- data.frame(deSolve::ode(func = psf_model,
                                     y = init, times = time, parms = params_excl2)) 
out_excl2b <- data.frame(deSolve::ode(func = psf_model,
                                      y = c(pA = 0.7, pAlpha = 0.7), times = time, parms = params_excl2)) 


# Plotting --------
make_sim_plot <- function(desolve_out) {
  desolve_out$pB = 1-desolve_out$pA
  out_long <- 
    desolve_out %>%
    select(-pAlpha) %>%
    pivot_longer(cols = pA:pB)
  sim1 <- 
    ggplot(out_long) +
    geom_path(aes(x = time, y = value, color = name), size = 1) +
    scale_color_manual(values = c("#CC79A7", "#999999")) +
    ylim(c(0,1)) + 
    # ylab("Plant species frequency") + 
    ylab("") + 
    xlab("time") + 
    scale_x_continuous(breaks = c(0, 50, 100)) +
    theme_plots() + 
    theme(axis.title = element_text(size = 10))
  
  return(sim1)
}

make_params_plot <- function(params) {
  
  color_func <- function(x) {
    ifelse(x < 0, "darkred", "royalblue4")
  }
  df <- data.frame(x = c(0,0,1,1),
                   y = c(0,1,0,1),
                   type = c("M", "P", "M", "P"))
  
  params_plot <- 
    ggplot(df) +
    annotate("text", x = 0, y = 1.15,  size = 3.15, label = "Plant 1") + 
    annotate("text", x = 1, y = 1.15,  size = 3.15, label = "Plant 2") + 
    annotate("text", x = 0, y = -0.15, size = 3.15, label = "\nSoil\nmicrobes A") + 
    annotate("text", x = 1, y = -0.15, size = 3.15, label = "\nSoil\nmicrobes B") + 
    geom_segment(aes(x = 0, xend = 0, y = 0.1, yend = 0.9),
                 arrow = arrow(length = unit(0.03, "npc")),
                 size = abs(params["m1A"]), 
                 color = alpha(color_func(params["m1A"]), 1)) +
    geom_segment(aes(x = 0.05, xend = 0.95, y = 0.1, yend = 0.9),
                 arrow = arrow(length = unit(0.03, "npc")),
                 size = abs(params["m2A"]), 
                 color = alpha(color_func(params["m1B"]),1)) +
    geom_segment(aes(x = 0.95, xend = 0.05, y = 0.1, yend = 0.9),
                 arrow = arrow(length = unit(0.03, "npc")),
                 size = abs(params["m1B"]), 
                 color = alpha(color_func(params["m2A"]), 1)) +
    geom_segment(aes(x = 1, xend = 1, y = 0.1, yend = 0.9),
                 arrow = arrow(length = unit(0.03, "npc"), type = "closed"),
                 size = abs(params["m2B"]), 
                 color = alpha(color_func(params["m2B"]), 1)) +
    
    # Plant cultivation of microbes
    geom_segment(aes(x = -0.35, xend = -0.35, y = 0.9, yend = 0.1), size = 0.15, linetype = 1,
                 arrow = arrow(length = unit(0.03, "npc"))) +
    geom_segment(aes(x = 1.35, xend = 1.35, y = 0.9, yend = 0.1), size = 0.15, linetype = 1,
                 arrow = arrow(length = unit(0.03, "npc"))) +
    
    annotate("text", x = 0, y = 0.5, 
             label = TeX(paste0(params["m1A"])), 
             angle = 90, vjust = -0.25, size = 3) + 
    annotate("text", x = 1, y = 0.5, 
             label = TeX(paste0(params["m2B"])), 
             angle = 90, vjust = 1.5, size = 3) + 
    annotate("text", x = 0.75, y = 0.75, 
             label = TeX(paste0(params["m2A"])), 
             label = TeX(paste0("m_{2A} = ", params["m2A"])), 
             angle = 40, vjust = -0.25, size = 3) + 
    annotate("text", x = 0.25, y = 0.75, 
             label = TeX(paste0(params["m1B"])), 
             angle = -40, vjust = -0.25, size = 3) + 
    xlim(c(-0.4, 1.4)) + 
    coord_cartesian(ylim = c(-0.25, 1.25), clip = "off") + 
    theme_void() +
    theme(legend.position = "none",
          plot.caption = element_text(hjust = 0.5, size = 10))
  return(params_plot)
}


plot_traj1 <- make_sim_plot(out_coex) + ylab("")
plot_traj2 <- make_sim_plot(out_excl) + ylab("")
plot_traj3 <- {make_sim_plot(out_priority) + ylab("")} + 
  {make_sim_plot(out_priority2) + ylab("")}
plot_traj4 <- {make_sim_plot(out_excl2) + ylab("")} + 
  {make_sim_plot(out_excl2b) + ylab("")}


(pp2 <- 
  base_coex_plot + 
  labs(
    x = "<span style = 'font-size:10pt;color:grey25;'>
    *Do plants grow better (destabilization) or worse (stabilization)<br>
    in soils conditioned by conspecifics vs. heterospecifics?*
    </span>",

    y = "Fitness differences<br>
    <span style = 'font-size:10pt;color:grey25;''>
    *Do species differ in their average responses<br>
    to conditioned vs. reference soil communities?*
    </span>",
    
  ) +  
  annotate("text", x = 0.9, y = 0.25, hjust = 0.95, 
           label = "Coexistence", size = 5) +
  annotate("text", x = 0.1, y = Inf, vjust = 2, hjust = 0, 
           label = "Species\nexclusion", size = 5) +
  annotate("text", x = -0.1, y = Inf, vjust = 2, hjust = 1, 
           label = "Species\nexclusion", size = 5) +
  annotate("text", x = -0.9, y = 0.35, hjust = 0, 
           label = "Priority\neffects", size = 5) +
    # geom_point(#"text", label = "outcome\n1", fontface = "italic", color = "midnightblue",
    #          aes(x = metrics(params_excl2)["SD"], 
    #              y = abs(metrics(params_excl2)["FD"])) + 
    
  annotate("text", label = "outcome\n1", fontface = "italic", color = "midnightblue",
           x = metrics(params_excl2)["SD"], 
           y = abs(metrics(params_excl2)["FD"])) + 
  annotate("text", label = "outcome\n2", fontface = "italic", color = "midnightblue",
           x = metrics(params_priority)["SD"], 
           y = abs(metrics(params_priority)["FD"])) + 
  annotate("text", label = "outcome\n3", fontface = "italic", color = "midnightblue",
           x = metrics(params_excl)["SD"], 
           y = metrics(params_excl)["FD"]) + 
  annotate("text", label = "outcome\n4", fontface = "italic", color = "midnightblue",
           x = metrics(params_coex)["SD"], 
           y = metrics(params_coex)["FD"]) + 
  annotate("text", label = c("-1", "-0.5", "0.5", "1"), 
           x = c(-0.95, -0.5, 0.5, 0.95), y = -.03, color = "grey25", size = 3) +
  scale_x_continuous(limits = c(-1,1), expand = c(0,0)) + 
  scale_y_continuous(limits = c(-0.15,1), expand = c(0,0)) + 
  annotate("segment", x = 0.2, xend = 0.7, y = -.07, yend = -.07,
           colour = "grey25", size = 0.5, arrow = arrow(length = unit(0.075, "inches"))) +
  annotate("segment", x = -0.2, xend = -.7, y = -.07, yend = -.07,
           colour = "grey25", size = 0.5, arrow = arrow(length = unit(0.075, "inches"))) +
  
  theme(axis.line = element_line(size = 0),
        plot.margin = margin(r = 20, t = 0, b = 10, l = 20),
        axis.text.x = element_blank(),
        axis.title.x = element_textbox_simple(
          width = NULL, halign = 0.5),
        axis.title.y = element_textbox_simple(
          width = NULL, halign = 0.5, orientation = "left-rotated")))
```

```
panelB <- {make_params_plot(params_excl2)    +labs(title = "outcome 1 (Exclusion)")} + {plot_traj4} 
panelC <- {make_params_plot(params_priority) +labs(title = "outcome 2 (Priority effects)")} + {plot_traj3} 
panelD <- {make_params_plot(params_excl)     +labs(title = "outcome 3 (Exclusion)")} + {plot_traj2}  
panelE <- {make_params_plot(params_coex)     +labs(title = "outcome 4 (Coexistence)")} + {plot_traj1}  

layout <- "
BBBBBBBAAAAAAADDDDDDD
BBBBBBBAAAAAAADDDDDDD
BBBBBBBAAAAAAADDDDDDD
BBBBBBBAAAAAAADDDDDDD
CCCCCCCAAAAAAAEEEEEEE
CCCCCCCAAAAAAAEEEEEEE
CCCCCCCAAAAAAAEEEEEEE
CCCCCCCAAAAAAAEEEEEEE
"

(full_plot <-
    {pp2 } + 
    {panelB} + 
    {panelC} + 
    {panelD} + 
    {panelE} + 
    plot_layout(design = layout) & 
    theme(legend.position = "none", # plot.title.position = "plot",
          plot.subtitle = element_text(face = "italic", color = "midnightblue", size = 12),
          plot.title = element_text(face = "italic", color = "midnightblue", size = 11),
          plot.tag = element_text(size = 12))
)
```

```
if(save_figures) {
  ggsave('figures/fig1.png', width = 13/1.15, height = 5/1.15, bg = "transparent")
  ggsave('figures/fig1.pdf', width = 13/1.15, height = 5/1.15, bg = "transparent")
}
```

Make the conceptual figure, but this time, show all four outcomes using positive values of all \(m\) terms.

```
params_coex = c(m1A = .2*2, m1B = 0.3*2, m2A = .4*2, m2B = .13*2, v = 1)
params_excl = c(m1A = .3*2, m1B = .2*2, m2A = 0.5*2, m2B = .15*2, v = 1)
params_priority = c(m1A = .9, m1B = 0.4, m2A = 0.38, m2B = 0.5, v = 1)
params_excl2 = c(m1A = .8, m1B = 0.1, m2A = 0.5, m2B = .15, v = 1)
time <- seq(0, 75)
init = c(pA = 0.3, pAlpha = 0.3)

out_coex <- data.frame(deSolve::ode(func = psf_model,
                                    y = init, times = time, parms = params_coex)) 
out_excl <- data.frame(deSolve::ode(func = psf_model,
                                    y = init, times = time, parms = params_excl)) 
out_priority <- data.frame(deSolve::ode(func = psf_model,
                                        y = init, times = time, parms = params_priority)) 

out_priority2 <- data.frame(deSolve::ode(func = psf_model,
                                         y = c(pA = 0.7, pAlpha = 0.7), times = time, parms = params_priority)) 
out_excl2 <- data.frame(deSolve::ode(func = psf_model,
                                     y = init, times = time, parms = params_excl2)) 
out_excl2b <- data.frame(deSolve::ode(func = psf_model,
                                      y = c(pA = 0.7, pAlpha = 0.7), times = time, parms = params_excl2)) 


# Plotting --------
plot_traj1 <- make_sim_plot(out_coex) + ylab("")
plot_traj2 <- make_sim_plot(out_excl) + ylab("")
plot_traj3 <- {make_sim_plot(out_priority) + ylab("")} + 
  {make_sim_plot(out_priority2) + ylab("")}
plot_traj4 <- {make_sim_plot(out_excl2) + ylab("")} + 
  {make_sim_plot(out_excl2b) + ylab("")}


pp2 <- 
    base_coex_plot + 
    labs(
      x = "<span style = 'font-size:10pt;color:grey25;'>
    *Do plants grow better (destabilizing) or worse (stabilizing)<br>
    in soils conditioned by conspecifics vs. heterospecifics?*
    </span>",
      
      y = "Fitness differences<br>
    <span style = 'font-size:10pt;color:grey25;''>
    *Do species differ in their average responses<br>
    to conditioned vs. reference soil communities?*
    </span>",
      
    ) +  
    annotate("text", x = 0.9, y = 0.25, hjust = 0.95, 
             label = "Coexistence", size = 5) +
    annotate("text", x = 0.1, y = Inf, vjust = 2, hjust = 0, 
             label = "Species\nexclusion", size = 5) +
    annotate("text", x = -0.1, y = Inf, vjust = 2, hjust = 1, 
             label = "Species\nexclusion", size = 5) +
    annotate("text", x = -0.9, y = 0.35, hjust = 0, 
             label = "Priority\neffects", size = 5) +

    annotate("text", label = "outcome\n1", fontface = "italic", color = "midnightblue",
             x = metrics(params_excl2)["SD"], 
             y = abs(metrics(params_excl2)["FD"])) + 
    annotate("text", label = "outcome\n2", fontface = "italic", color = "midnightblue",
             x = metrics(params_priority)["SD"], 
             y = abs(metrics(params_priority)["FD"])) + 
    annotate("text", label = "outcome\n3", fontface = "italic", color = "midnightblue",
             x = metrics(params_excl)["SD"], 
             y = metrics(params_excl)["FD"]) + 
    annotate("text", label = "outcome\n4", fontface = "italic", color = "midnightblue",
             x = metrics(params_coex)["SD"], 
             y = metrics(params_coex)["FD"]) + 
    annotate("text", label = c("-1", "-0.5", "0.5", "1"), 
             x = c(-0.95, -0.5, 0.5, 0.95), y = -.03, color = "grey25", size = 3) +
    scale_x_continuous(limits = c(-1,1), expand = c(0,0)) + 
    scale_y_continuous(limits = c(-0.15,1), expand = c(0,0)) + 
    annotate("segment", x = 0.2, xend = 0.7, y = -.07, yend = -.07,
             colour = "grey25", size = 0.5, arrow = arrow(length = unit(0.075, "inches"))) +
    annotate("segment", x = -0.2, xend = -.7, y = -.07, yend = -.07,
             colour = "grey25", size = 0.5, arrow = arrow(length = unit(0.075, "inches"))) +
    
    theme(axis.line = element_line(size = 0),
          plot.margin = margin(r = 20, t = 0, b = 10, l = 20),
          axis.text.x = element_blank(),
          axis.title.x = element_textbox_simple(
            width = NULL, halign = 0.5),
          axis.title.y = element_textbox_simple(
            width = NULL, halign = 0.5, orientation = "left-rotated"))


panelB <- {make_params_plot(params_excl2)    +labs(title = "outcome 1 (Exclusion)")} + {plot_traj4} 
panelC <- {make_params_plot(params_priority) +labs(title = "outcome 2 (Priority effects)")} + {plot_traj3} 
panelD <- {make_params_plot(params_excl)     +labs(title = "outcome 3 (Exclusion)")} + {plot_traj2}  
panelE <- {make_params_plot(params_coex)     +labs(title = "outcome 4 (Coexistence)")} + {plot_traj1}  


(full_plot <-
    {pp2} + 
    {panelB} + 
    {panelC} + 
    {panelD} + 
    {panelE} + 
    plot_layout(design = layout) & 
    theme(legend.position = "none", # plot.title.position = "plot",
          plot.subtitle = element_text(face = "italic", color = "midnightblue", size = 12),
          plot.title = element_text(face = "italic", color = "midnightblue", size = 11),
          plot.tag = element_text(size = 12))
)
```

```
if(save_figures) {
  ggsave('figures/fig1_MutualismsVersion.png', width = 13/1.15, height = 5/1.15, bg = "transparent")
  ggsave('figures/fig1_MutualismsVersion.pdf', width = 13/1.15, height = 5/1.15, bg = "transparent")
}
```

#### Other visualizations

##### Catterpillar-like plots

```
# For Live soil reference --------- 
live_ref_plot <- 
  live_ref %>%
  # remove unneeded columns
  select(Experiment, Study, author, #Species_pair = `Species pair`,
         mean_FD, sem_FD, mean_SD, sem_SD) %>%
  group_by(Study, author) %>%
  # make new column for absolute value of FD and SD
  mutate(n = n(), abs_mean_FD = abs(mean_FD), abs_mean_SD = abs(mean_SD)) %>%
  # Arrange within group, descending by FD
  arrange(-abs_mean_FD, .by_group = T) %>%
  # Arrange so that groups (studies) with most rows are at top
  arrange(-n) %>%
  ungroup %>% 
  # Add a new column with just the row number
  mutate(row_num = rev(row_number()))


helper <- live_ref_plot %>% 
  group_by(Study, author) %>% 
  summarize(mean_row = ceiling(mean(row_num)),
            min_row = min(row_num))
```

```
## `summarise()` has grouped output by 'Study'. You can override using the `.groups` argument.
```

```
fx_plot <- 
  ggplot(data = live_ref_plot) +
  scale_y_continuous(limits = c(0, 73), breaks = helper$mean_row, 
                     labels = helper$author, expand = c(0,0)) + 
  scale_x_continuous(limits = c(-1.5, 3.5)) + 
  geom_hline(yintercept = helper$min_row - .5, size = .1) +
  geom_vline(xintercept = 0, linetype = 1, size = .25, color = "grey50") +
  geom_pointrange(aes(x = abs_mean_SD, y = row_num,
                      xmin = abs_mean_SD-2*sem_SD,
                      xmax = abs_mean_SD+2*sem_SD),
                  color =  alpha("black", .6)) +
  geom_pointrange(aes(x = abs_mean_FD, y = row_num,
                      xmin = abs_mean_FD-2*sem_FD,
                      xmax = abs_mean_FD+2*sem_FD),
                  color = alpha("Sienna", .6)) +
  scale_color_manual(values = c("sienna", "black"), name = "",
                     labels = c("Fitness difference",
                                "(De)stabilization") ) + 
  geom_hline(yintercept = 0) + 
  ylab("") +
  xlab("") +
  # theme_gsk() +
  theme_bw() + 
  theme(axis.line.y= element_blank(), 
        legend.position = "top",
        panel.grid = element_blank(),
        axis.text.y = element_text(angle = 45, size = 8),
        axis.text.x = element_blank(),
        axis.ticks.x = element_blank())  

fx_plot
```

```
# For sterile reference soil ----------
XS_ref_plot <- 
  sterile_ref %>%
  # remove unneeded columns
  filter(mean_SD > -4) %>%
  select(Experiment, Study, author, #Species_pair = `Species pair`,
         mean_FD, sem_FD, mean_SD, sem_SD, control_type,study_control_type) %>%
  group_by(Study, author) %>%
  # make new column for absolute value of FD and SD
  mutate(n = n(), abs_mean_FD = abs(mean_FD), abs_mean_SD = abs(mean_SD)) %>%
  # Arrange within group, descending by FD
  arrange(-abs_mean_FD, .by_group = T) %>%
  # Arrange so that groups (studies) with most rows are at top
  arrange(-n) %>%
  ungroup %>% 
  # Add a new column with just the row number
  mutate(row_num = rev(row_number()))

XS_helper <- XS_ref_plot %>% 
  group_by(Study, author, study_control_type) %>% 
  summarize(mean_row = ceiling(mean(row_num)),
            min_row = min(row_num))
```

```
## `summarise()` has grouped output by 'Study', 'author'. You can override using the `.groups` argument.
```

```
xs_plot <- 
  ggplot(data = XS_ref_plot) +
  scale_y_continuous(limits = c(0, nrow(XS_ref_plot) + 1), 
                     breaks = XS_helper$mean_row, labels = XS_helper$author, expand = c(0,0)) + 
  scale_x_continuous(limits = c(-1.5, 6.5)) + 
  geom_hline(yintercept = XS_helper$min_row - .5, size = .1) +
  geom_vline(xintercept = 0, linetype = 1, size = .25, color = "grey50") +
  geom_pointrange(aes(x = abs_mean_SD, y = row_num,
                      xmin = abs_mean_SD-2*sem_SD,
                      xmax = abs_mean_SD+2*sem_SD),
                  color =  alpha("black", .6)) +
  geom_pointrange(aes(x = abs_mean_FD, y = row_num,
                      xmin = abs_mean_FD-2*sem_FD,
                      xmax = abs_mean_FD+2*sem_FD),
                  color = alpha("Sienna", .6)) +
    # geom_point(aes(x = -1.5, y = row_num,fill = control_type), shape = 21, stroke = 0, size = 2) +
  scale_color_manual(values = c("sienna", "black"), name = "",
                     labels = c("Fitness difference",
                                "(De)stabilization") ) + 
  geom_hline(yintercept = 0) + 
  ylab("") +
  xlab("") +
  theme_bw() + 
  theme(axis.line.y= element_blank(), 
        #legend.position = "top",
        panel.grid = element_blank(),
        axis.text.y = element_text(angle = 45, size = 8),
        axis.text.x = element_blank(),
        axis.ticks.x = element_blank())

xs_plot
```

```
## Warning: Removed 3 rows containing missing values (geom_segment).
```

```
combined_catterpillars <- xs_plot + fx_plot

if(save_figures) {
  ggsave("figures/effectSize-catterpillars.png", combined_catterpillars,
         height = 12, width = 6)
}
```
